# Potato yield can be predicted using drone-captured and environmental measurements early in the growing season

**DOI:** 10.64898/2026.03.09.709817

**Authors:** Angelika Vižintin, Maja Zagorščak, Eva Turk, Maja Križnik, Marko Petek, Katja Stare, Bernhard Wurzinger, Maroof Ahmed Shaikh, Guus Heselmans, Josef Söllinger, Anže Županič, Pieter-Jelte Lindenbergh, Jeroen Bakker, Robert Graveland, Ivana Imerovski, Salome Prat, Christian Bachem, Markus Teige, Bianca Doevendans, Alexandra Ribarits, Jan Zrimec, Kristina Gruden

## Abstract

Accurate pre-harvest prediction of crop yield informs variety selection, optimizes management, and accelerates breeding. As potato is the world’s leading non-grain staple, here we evaluate a diverse panel of varieties in a three-year field trial across five European locations. Canopy development and environmental parameters are monitored throughout the growing season using drone-based imaging, in-field sensors and gene expression measurements, while tuber yield and quality traits are quantified at harvest. We show that these data enable the identification of climate-resilient, high-yielding genotypes and support the development of machine learning models that explain over 80% of yield variation in independent test sets. Strikingly, measurements collected within the first two months after planting achieve predictive performance comparable to models trained on full-season data. Model interrogation further shows that over 70% of yield variation can already be predicted based on a simple five-parameter linear equation. Our framework thus demonstrates the potential of integrative field phenotyping and data-driven modeling to improve variety selection across heterogeneous environments.

## 1. Introduction

The ability to accurately predict tuber yield and quality across diverse potato cultivars and environmental conditions has been a central objective in agriculture research. Potato as the world’s third most important food crop feeds over a billion people globally, and analytical tools to understand its yield as affected by the environment and growing conditions can help growers to further improve and optimize its productivity ^1,2^. Typically, for high yield, quick emergence and canopy development are desired at the start of the growing season. During growth, various factors can affect tuber sizing and dry matter, including environmental parameters and biotic stress, as well as agricultural management practices employed by growers to comply with different target market requirements ^3,4^. Monitoring plant growth and growing conditions is therefore becoming essential to acquire data, based on which crop yield can be studied, modelled and predicted ^5–7^. Predicting potato yield during the growing season enables farmers to make informed decisions regarding crop management practices like irrigation, fertilization, and pest control. It also supports breeding programs in the identification of varieties that perform well under diverse environmental conditions and stress factors, and enhances market stability and food security.

Technological advances have created unprecedented opportunities to collect detailed environmental and crop data, which can be integrated into yield prediction models. Apart from various in-field sensors that routinely collect environmental variables to characterize growing conditions, the monitoring of crop growth has been revolutionised over the past decade by the increasing use of uncrewed aerial vehicles, or ‘drones’, equipped with different visible (VIS), near-infrared (NIR), and thermal sensors ^8^, assessing crop growth, vigour and stress. Drone-collected vegetation metrics, coupled with measurements of environmental parameters such as temperature, precipitation and soil properties, have been shown to improve yield prediction across a range of crop species, including potato ^1,5,9^, rice ^10,11^ and maize ^12–14^. Data-driven methods for crop yield prediction typically encompass statistical and machine learning (ML) approaches ^6,15^. With the increasing availability of data from drone and in-field sensor-based technologies ^6,16^, the predictive power and accuracy of crop yield prediction are also expected to improve ^1,6^. However, comparing the performance of multiple cultivars across locations, environmental conditions, and growing seasons (years), requires a large number of field trials with standardized measurements of pre- and post-harvest vegetation, environmental, yield and quality parameters, collected across varieties and time points ^17^.

From the perspective of predictive model complexity, previous studies in grapevine and potato crops have shown that crop yield can be accurately predicted using simple equations comprising only a handful of variables measured during or before the growing season ^18,19^. For instance, the “Bordeaux equation” is a well-known formula developed to predict the quality and price of a Bordeaux wine vintage based on weather data. The equation uses a linear model in which quality is expressed as a function of winter rainfall, average growing season temperature, and harvest rainfall ^18^. Similarly, as early as 1978, potato yield was shown to be predictable using a linear equation with temperature and solar radiation as key variables ^19^. However, despite the evidence that final post-harvest crop yield can be predicted from plant development and environmental data captured during the growing season, it remains unclear how early such predictions can be made and how small of a set of selected variables is sufficient to enable such predictions.

Here, we present an extensive 3 year field trial conducted across 5 locations in Europe, involving 44 different potato varieties representing diverse genotypes. This broad genotypic representation is essential, given that different genotypes inherently exhibit distinct yield potentials and responses to environmental stresses. Vegetation metrics were captured using drones, and environmental variables collected during the growing season, while potato yield and quality indices were measured at harvest. Focusing on tuber yield and quality, we show the ability to identify resilient and stable-yielding varieties across the different locations and growing conditions. Tuber yield and quality are found to be strongly associated with the drone-captured vegetation and environmental data, with machine learning (ML) models achieving highly predictive accuracy for tuber yield (*R*^2^ ∼ 0.8), prompting further investigation using ML algorithms. We find that a performance within ∼1% of the maximum can also be achieved using explanatory data collected during only the first two months of plant growth. Finally, we query the data-derived ML models to obtain the most informative and thus highly predictive features (vegetation and environmental variables) for potato yield, demonstrating that simple 5-parameter linear equations can already explain over ∼70% of the variation of tuber yield. The developed approach thus represents an important stepping stone towards precision agriculture for potato yield and quality prediction. It demonstrates the potential of field trial data to facilitate the analysis and selection of best-performing varieties across diverse conditions, and to revolutionize farming by enabling early (already within 2 months) and straightforward (only a couple of key measured variables) yield predictions with a high accuracy.

## 2. Results

### 2.1 Potato field trials across Europe capture plant growth and yield diversity

To capture the effects of potato environmental and growth conditions on tuber yield and quality, we conducted potato field trials for three years at five European locations (Figure 1A, Supp. table S1.1: Netherlands - Zeeland; Austria - Fuchsenbigl and Großnondorf; Spain - Silla-Valencia; and Serbia - Vojvodina). A total of 44 tetraploid potato varieties were included in the study (Supp. table S1.2, Methods M1), selected to represent diversity in maturity, parental lineage, market use, and resilience to environmental stress (Supp. table S1.3). The varieties were grown under irrigated and non-irrigated conditions, with approximately half of the trials conducted under irrigation (Supp. table S1.1). The experimental design for field trials in Valencia, Vojvodina and Zeeland was to plant each variety in duplicate plots (i.e., two distinct sections per variety), yielding up to 88 plots per field trial, while at Fuchsenbigl and Großnondorf, ∼13 varieties were cultivated in quadruple plots (Supp. tables S1.1, S1.2). This yielded a total of 1,165 plots studied across all field trials (Figure 1B).

**Figure 1.**
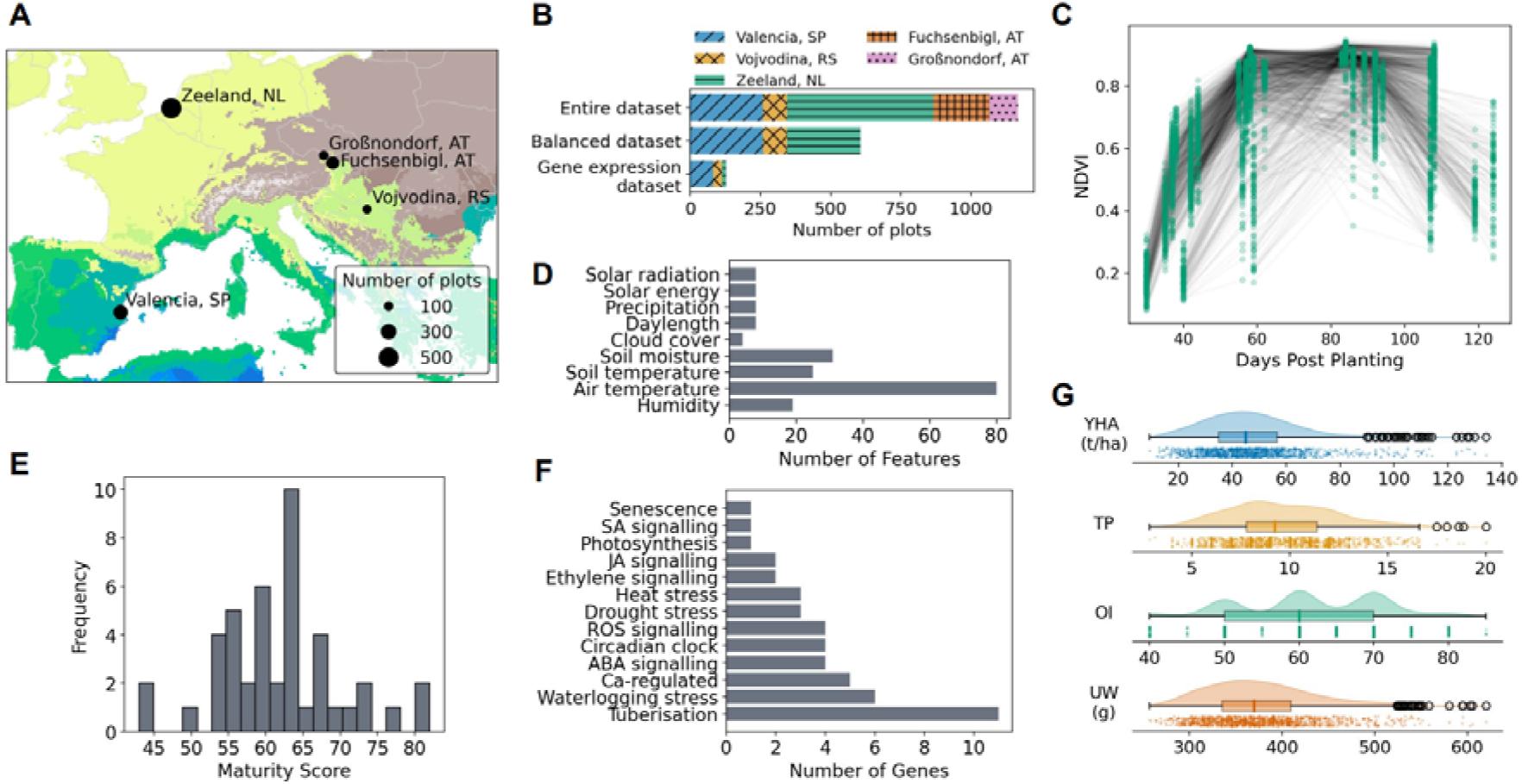
Potato field trials across Europe capture plant growth and yield diversity. (A) Locations of field trials with number of plots per location, alongside Köppen-Geiger climate maps for 1901–2099 ^22^: The corresponding climate classifications of the field trial locations are: Dfb (brown): continental, fully humid, warm summer (Fuchsenbigl and Großnondorf), Bsk (blue-green): dry, steppe, cold (Valencia), Cfa (light green): temperate, fully humid, hot summer (Vojvodina), Cfb (yellow): temperate, fully humid, warm summer (Zeeland). (B) Bar plot illustrating the number of plots contributed by each field trial location to the three main datasets: the entire dataset (*N* = 1,165 plots), the balanced dataset (*N* = 606 plots) used for predicting tuber yield and quality variables, and the gene expression dataset (*N* = 128 plots). (C) Scatter plot of drone-derived normalised difference vegetation index (NDVI) as a function of days post-planting. Lines connect the measurements from the same plot. (D) Count of environmental features (calculated from collected environmental variables) by category. The features were derived from two primary sources: in-field soil sensors (soil temperature and moisture) and weather station data (all other environmental variables). (E) Histogram of maturity scores across varieties. (F) Count of genes selected for expression analysis by pathway annotation. Acronyms: abscisic acid (ABA), jasmonic acid (JA), salicylic acid (SA). (G) Raincloud plots illustrating the distributions of tuber yield and quality variables: yield per hectare (YHA, tons per hectare), number of tubers per plant (TP), overall impression (OI) scored on a scale from 10 (very bad) to 90 (very good), and underwater weight (UW, grams).

To monitor plant growth and development throughout the growing season, multiple drone flights were conducted over each trial field on several dates (Supplementary Table S1.4, Methods M1). Vegetation metrics collected across all field trials included percent of vegetation cover, the normalised difference vegetation index (NDVI, Figure 1C), the weighted difference vegetation index (WDVI), and the chlorophyll index (CI_RED) based on NIR and RED bands (Methods M1). Vegetation metrics from drone flight dates closest to 40, 60, 90 and 120 days post-planting (dpp) were included in the dataset (Supp. table S1.4). In addition, environmental data were recorded using in-field sensors measuring soil temperature and moisture that were complemented with data from nearby weather stations to capture a comprehensive spectrum of environmental parameters during the growing season. To generate predictive features, we segmented the full growing season data into four time windows (0–40, 40–60, 60–90, and 90–120 dpp) to match the measurement periods with the vegetation metrics. Various summary statistics (e.g. mean, median, maximum) were then calculated for each environmental variable within each window, resulting in a total of 191 environmental features (Figure 1D, Supp. table S1.5, Methods M2). Moreover, based on previous experiments, a maturity score was assigned to each variety to reflect earliness, with early varieties receiving high scores and late varieties receiving low scores (Supp. table S1.3, Figure 1E). Furthermore, gene expression of 50 genes was measured from leaves at the tuber initiation stage for 16 varieties in five field trials (128 plots in total, Figure 1B, Supp. table S1.1, Supp. table S1.2, Methods M3). These included 3 housekeeping genes and 47 targets associated with physiological processes related to plant development and stress responses with a particular focus on the regulation of tuberisation ^20,21^ (Figure 1F, Supp. table S1.6). Finally, the post-harvest plant performance was assessed at the end of each growing season in each field trial using four tuber yield and quality variables, including yield per hectare (YHA, in tons per hectare), number of tubers per plant (TP), overall impression (OI, an expert quality assessment), and underwater weight (UW, indicative of starch content, in grams) (Figure 1G, Methods M1).

To ensure data quality and balanced representation across conditions, a subset of the initial dataset was selected (Methods M4). Data from field trials conducted at Fuchsenbigl and Großnondorf were excluded from this selected dataset, as they captured only 13 of the total 44 varieties and vegetation metrics were available only for a single drone flight date, making these trials unsuitable for model training. Importantly, to mitigate location bias and reduce the risk of overfitting, we included data from only one field trial per year from Zeeland, the most overrepresented location. This stringent selection yielded a balanced dataset comprising 606 plots, which was used for correlation analysis and predictive modelling (Figure 1B, Supp. table S1.1).

### 2.2 Differences in environmental conditions are reflected across all collected data

The different field trials were conducted under varying environmental conditions (Figure 2A, Supp. figure S2.1). Overall, irrigated plots showed similar or lower soil temperatures and similar or higher soil moisture than non-irrigated plots when conducted simultaneously at a specific location, namely Fuchsenbigl and Zeeland (Figure 2A, Methods M5, Supp. table S1.1), suggesting a cooling and moisturizing effect of irrigation. However, while differences between irrigated and non-irrigated plots were statistically significant at Fuchsenbigl, for both mean daily soil temperature (Wilcoxon rank-sum test *p*-value ≤ 8.9×10^−14^) and soil moisture (Wilcoxon rank-sum test *p*-value ≤ 4.2×10^−3^), they were largely non-significant in the Zeeland trials (Supp. Table S2.1). In the latter, the only significant difference was observed for soil temperature in the 2021 trials, where the irrigated plot was significantly cooler (Wilcoxon rank-sum test *p*-value = 1.7×10^−3^). Since the Zeeland field trials were the northernmost, with the highest amount of precipitation, the lowest impact of irrigation was expected (Fig.1A). Principal Component Analysis (PCA) performed on all environmental features showed that plots from different field trials occupied distinct regions in the projection space (Methods M5). The exceptions were trials from Zeeland and Fuchsenbigl from the same year, which overlapped, indicating similar environmental conditions despite differences in irrigation settings (Figure 2B). A similar pattern of separation was observed when t-distributed Stochastic Neighbor Embedding (t-SNE) was applied to the drone-captured vegetation metrics: plots from nearly every trial occupied distinct regions in the projection space (Figure 2C, Supp. table S1.1, Methods M5). The only exception was the irrigated and non-irrigated Zeeland trials from the same year, which showed substantial overlap. Furthermore, PCA of all measured gene expression values across samples also revealed separation by location (Figure 2D), likely reflecting site-specific differences in environmental conditions. Overall, the analyses indicated that the environmental conditions differed across field trials, with the exception of the concurrent irrigated and non-irrigated trials in Zeeland and Fuchsenbigl.

**Figure 2.**
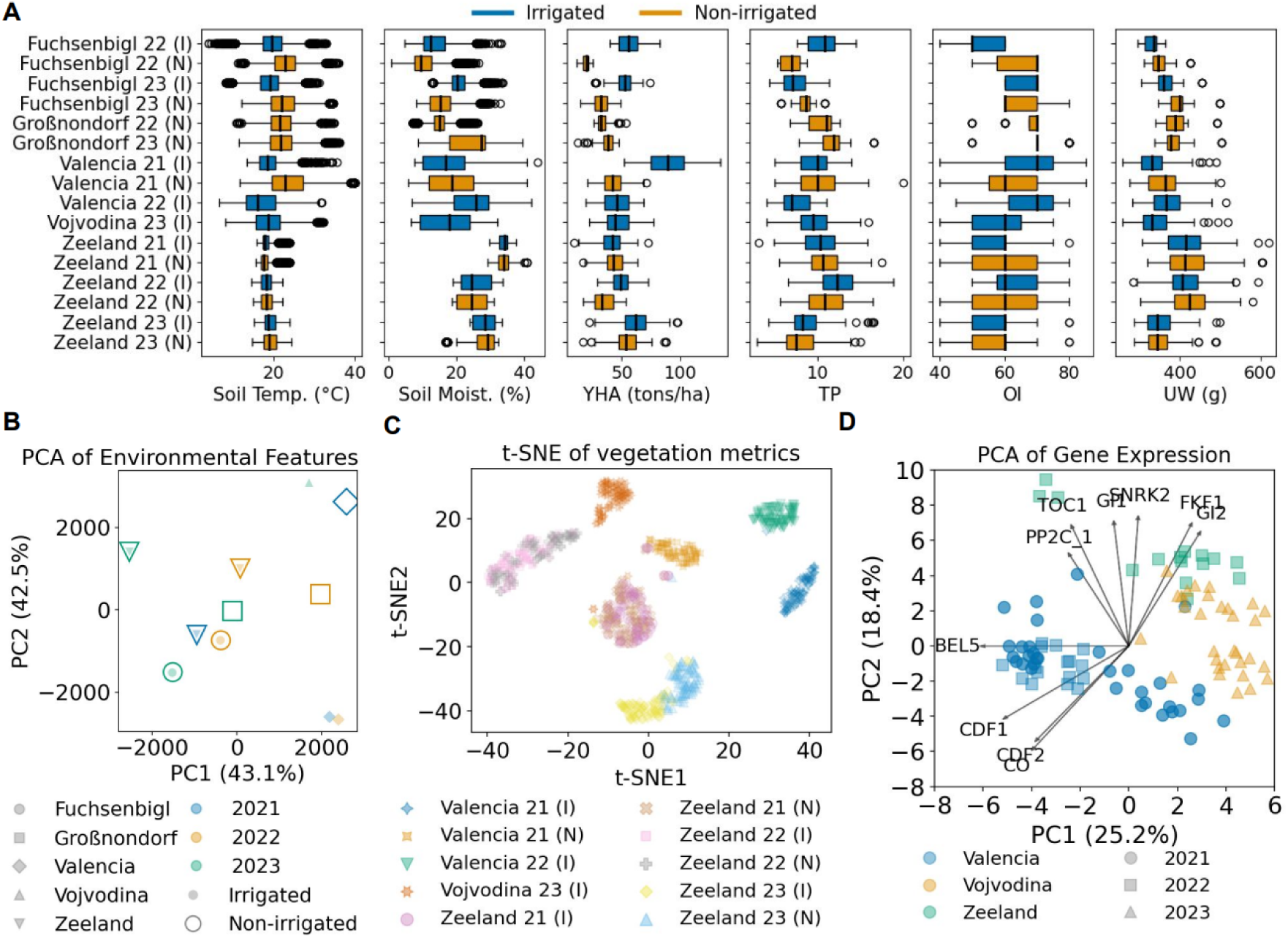
Differences in environmental conditions are reflected across all collected data. (A) Boxplots showing distributions of soil temperature (°C), soil moisture (%, measured at 10 cm except for Fuchsenbigl and Großnondorf, where it was measured at 5 cm), yield per hectare (YHA, tons per hectare), number of tubers per plant (TP), overall impression (OI) scored on a scale from 10 (very bad) to 90 (very good), and underwater weight (UW, grams). Blue boxes indicate irrigated field trials (I), while orange boxes indicate non-irrigated trials (N). (B) Principal Component Analysis (PCA) of environmental features across all field trials. Each point represents a unique field trial, positioned according to the first two principal components (PC1 and PC2, respectively), which together explain 85.6% of the variance. (C) t-SNE visualization of vegetation metrics. Each point represents a plot from a field trial, projected into two dimensions (t-SNE1 and t-SNE2). Colors and markers distinguish between field trials, with (I) denoting irrigated and (N) denoting non-irrigated conditions. The positions of the points in the projected space illustrate similarities in vegetation profiles between plots and trials. Note that Fuchsenbigl and Großnondorf trials were excluded from this analysis due to having vegetation metrics data only for a single drone flight. (D) PCA biplot of log_2_-transformed gene expression data. Each point represents a plot, positioned according to the first two principal components (PC1 and PC2), which together explain 43.6% of the total variance. The vectors originating from the center represent the top 10 genes contributing most strongly to the observed variance, where the length and direction of a vector indicate the gene’s overall contribution to sample separation and its correlation with the first two principal components.

The observed environmental differences across field trials were also reflected in the final tuber yield and quality variables (Figure 2A). While the median YHA across the entire dataset was 45.1 t/ha, the value varied substantially among field trials, ranging from a minimum of 19.8 t/ha in the non-irrigated Fuchsenbigl trial in 2022 to a maximum of 89.5 t/ha in the irrigated Valencia trial in 2021. A similar trend was observed for the remaining yield and quality variables (Figure 2A). YHA and TP were typically higher under irrigated conditions, with statistically significant differences (Wilcoxon Rank-Sum *p*-value ≤ 0.016,

Supp. Table S2.2) across trials conducted simultaneously at a specific location, except for the two trials conducted in Zeeland in 2021. Median OI was consistent across the Zeeland trials, but fluctuated in Fuchsenbigl. Specifically, the median OI for the irrigated trial was significantly lower in 2022 (Wilcoxon Rank-Sum test *p*-value = 1.5×10^-8^), whereas in 2023 the differences were not significant (Supp. Table S2.2). The median UW did not differ among the field trials in Zeeland but was significantly higher in non-irrigated trials in Fuchsenbigl (Wilcoxon Rank-Sum test *p*-value ≤ 5.2×10^-4^, Supp. Table S2.2). Collectively, these results demonstrate that site-specific environmental conditions are strongly reflected across all collected datasets, not only in environmental features, but also in the drone-derived vegetation metrics, gene expression profiles, as well as yield and quality traits.

### 2.3 High performing varieties show less than 35% variation in yield across diverse field conditions

We next analysed the yield and quality traits to evaluate the performance and stability of the tested varieties across the field trials. To this end, we computed the weighted median and weighted relative standard deviation (RSD) for each variety across the four tuber yield and quality variables: yield per hectare (YHA, Figure 3A), number of tubers per plant (TP, Figure 3B), overall impression (Figure 3C), and underwater weight (Figure 3D) (Methods M5). The weighted median reflects the typical performance/value of a variety for the given trait, whereas the weighted RSD captures its consistency across trials conducted under different environmental conditions, including differences in heat and drought stress. Thus, a high weighted median (upper part of the plots) represents varieties with higher YHA, TP, OI or UW, while a low RSD (left part of the plots) indicates varieties that maintain stable performance under different growing conditions. The weighted medians of YHA per variety across all field trials ranged from 27.3 t/ha (for Charlotte) to 68.5 t/ha (for Arizona). For UW, the median values clearly followed the expected market segmentation: starch varieties exhibited the highest median UW (462.0 - 501.0 g), followed by crisp and French fry varieties, with table varieties demonstrating the lowest median UW (303.0 - 405.0 g). The range of RSD values differed markedly among the four tuber yield and quality variables, with YHA exhibiting the widest range (0.20 - 0.47) and UW the narrowest (0.07 - 0.16) (Supp. figure S3.1).

**Figure 3.**
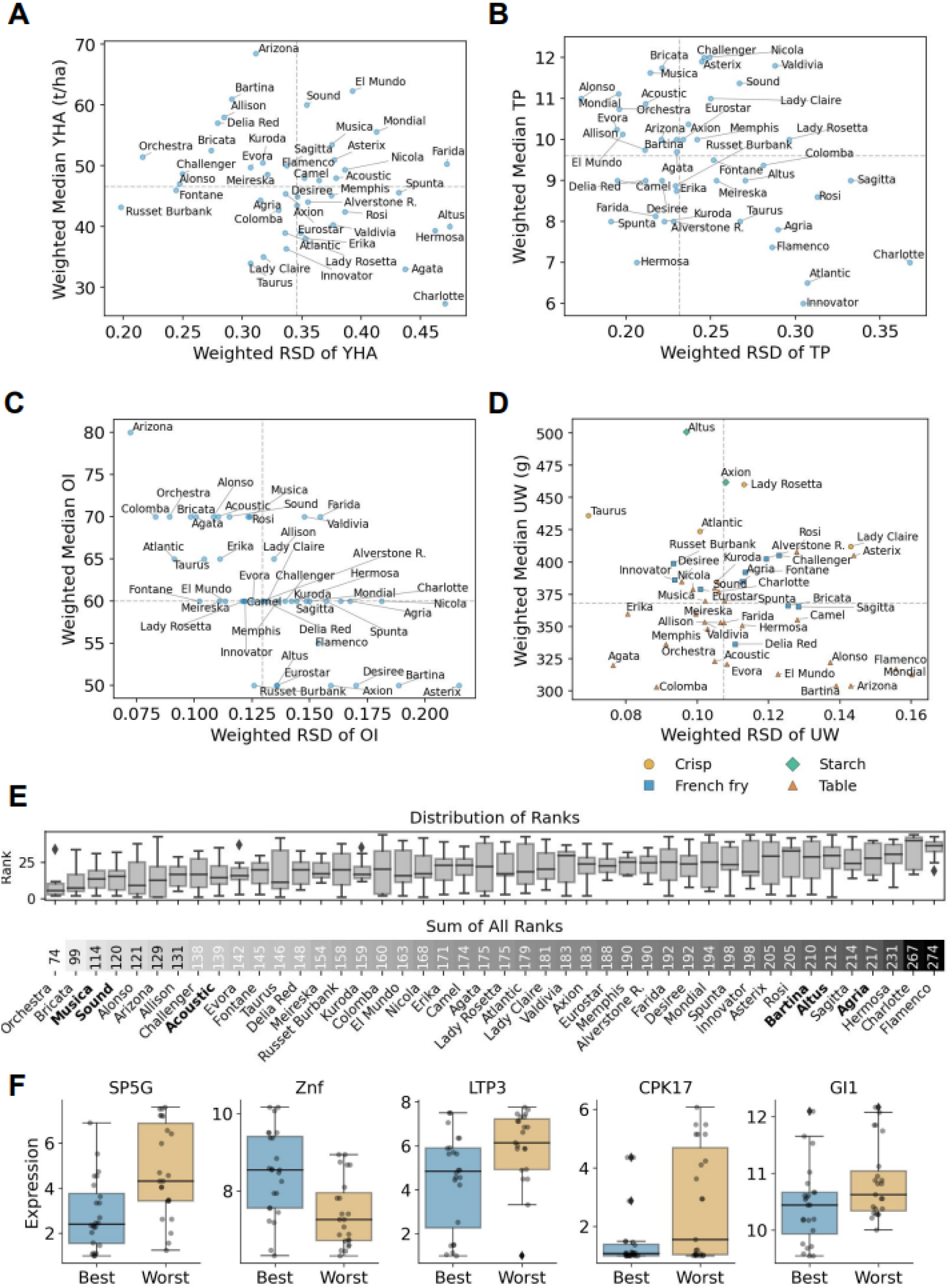
High performing varieties show less than 35% variation in yield across diverse field conditions. (A) Weighted Median YHA vs. Weighted Relative Standard Deviation (RSD) of YHA. (B) Weighted Median TP vs. Weighted RSD of TP. (C) Weighted Median OI vs. Weighted RSD of OI. (D) Weighted Median UW vs. Weighted RSD of UW with marker symbols and colors indicating the market type of the potato varieties. (E) Summary of rank metrics across varieties. The upper panel displays boxplots for the distribution of median and RSD ranks across the four tuber yield and quality traits allowing comparison of consistency and relative performance among varieties. Varieties were ranked such that those with the highest median value for a trait received a rank of 1, whereas for the RSD of a trait, the variety with the lowest RSD received a rank of 1. The lower panel shows a heatmap of the sum of ranks for each variety. In bold, we highlight the three varieties with the lowest (best overall performance) and the three varieties with the highest (worst overall performance) sum of ranks as compared with available gene expression data. (F) Boxplots of log_2_-transformed expression values for genes showing significant differences between the best-performing (blue) and worst-performing (orange) varieties, and individual data points overlaid in gray.

To enable comparison across all four traits, we also ranked the varieties separately for each trait, with independent rankings for the median values and for the RSD (Supp. figure S3.2, Methods M5). Specifically, varieties were ranked such that those with the highest median value for a trait received a rank of 1. For stability, the variety with the lowest RSD received a rank of 1, reflecting greater consistency across trials (see the distribution of ranks per variety in Figure 3E). The ranks were then summed per variety, with lower total scores indicating a potentially more stable and high-performing variety. Analysis of rank distributions across the four traits (Supp. table S3.1) showed that no variety consistently achieved top ranks for all traits. Both the rank range (difference between maximum and minimum rank) and the standard deviation per variety were relatively high, indicating that overall performance is strongly trait-dependent and that there is no universally stable top performer across all traits. The variety Memphis exhibited the lowest rank range (16) and standard deviation (6.5); however, it still spanned more than one third of the possible ranks (16 of 44), reflecting a relatively broad spread. In the present dataset, the overall highest-ranked variety (Figure 3E: Orchestra, range 2–34) and worst variety (Flamenco, range 19–43) had overlapping rank distributions, illustrating the substantial variability of performance across traits.

Next, to assess whether differences between high- and low-performing varieties can be explained by specific signaling or transcriptional module activation, we compared the three highest-ranked (Musica, Sound, Acoustic) and three lowest-ranked (Agria, Altus, Bartina) varieties with available gene expression data (Fig. 3E, Methods M5). Initially, differences in the distributions of log_2_-expression values between the highest- and lowest-performing varieties across the combined field trials were assessed (Supp. table S3.2). This revealed that the genes *StSP5G* (Self-pruning 5G), *StLTP3* (Lipid transfer protein 3), *StCPK17* (Calcium-dependent protein kinase 17), and *StGI1* (Gigantea 1) had significantly lower median expression values in the best performing varieties (Wilcoxon rank-sum test *p*-value = 0.001, 0.019, 0.029, and 0.036, respectively, Figure 3F), while the median expression value for *ZNF* was significantly lower in the worst performing varieties (Wilcoxon rank-sum test *p*-value = 0.003, Figure 3F). Next, a differential expression profile was generated by computing the difference between the median log_2_-transformed expression values of the best and worst performing varieties within each field trial (Supp. figure S3.3). Lower expression of *StSP5G* and *StGI1* in better performing plants was expected as they are both negative regulators of tuberisation ^23,24^. Less information is available on the function of the *StZNF* (Zinc finger transcription factor)*, StLTP3* and *StCPK17* genes. *StZNF* was included as drought-regulated and *StLTP3* as waterlogging-regulated based on publicly available stress-specific transcriptomics datasets, while *StCPK17* was included as a Ca^2+^-regulated protein kinase ^25^. Here we found these genes to be linked with potato performance and thus to represent potentially interesting future breeding targets. The results also demonstrate that with a sufficiently large and geographically diverse dataset, the field trial data enable robust performance ranking and the identification of resilient, stable-performing varieties across diverse conditions.

### 2.4 Analysis of vegetation and environmental data demonstrates their predictive value for plant performance

We next examined the correlations among the various collected data types. Prior to this, we split the balanced dataset into training and test sets based on variety (Methods M4). This was done to ensure that all correlation analyses were based solely on a training dataset, thereby preventing information from the test set from influencing feature selection and model development (i.e. to avoid data leakage). Specifically, four varieties (10%) were held out as the test set, and the remaining 40 varieties (90%) were used for training. We applied the same partitioning strategy to the gene expression dataset to ensure consistency in variety allocation. Correlation analysis was performed between: (i) the tuber yield and quality traits, i.e. the targets, to explore relationships among target variables; (ii) features and target variables, to gain insights into the relationships between environmental and vegetation features and target variables and to identify features with predictive potential; (iii) features, to analyze feature-feature interactions and to guide feature selection by identifying and reducing redundancy (Methods M5).

Firstly, among the four tuber yield and quality traits, when aggregated by variety, we observed moderate correlations between median YHA and median UW (*ρ* = –0.63, *p*-value = 1.5×10^−5^), median YHA and the median TP (*ρ* = 0.46, *p*-value = 2.9×10^−3^) and median UW and median OI (*ρ* = 0.45, p-value = 3.7×10^−3^) (Supp. figure S4.1, Supp. Table S4.1). Secondly, correlation analysis between the targets (i.e. tuber yield and quality variables), and vegetation metrics (Supp. tables S4.2-4.5) revealed moderate positive correlations between YHA and vegetation metrics measured at 90 and 120 days post planting (*ρ* = 0.39- 0.52, *p*-value < 10^−16^). Similarly, moderate positive correlations were observed between the number of tubers per plant and vegetation metrics measured at 60 days post-planting (*ρ* = 0.45–0.51, *p*-value < 10^−16^). Considering the gene expression data, moderate correlations were observed between YHA and the expression of some genes including *StRBA* (Rubisco activase), *StCAT1* (Catalase 1), and *StPPDK* (Pyruvate orthophosphate dikinase) (*ρ* = 0.59, 0.58, and 0.53, respectively, *p*-value ≤ 2.4×10^−9^) (Supp. tables S4.6-4.9), which are involved in photosynthesis, ROS signalling, and response to waterlogging.

Finally, correlation analyses among features identified numerous pairs of highly correlated features (Supp. figure S4.2). Strong correlations were observed among vegetation indices (NDVI, WDVI and CIRED) measured on the same day (Supp. figure S4.3 and Supp. table S4.10) and environmental features derived from the same variable within the same time period (Supp. file S1, Supp. file S2). This indicated potential collinearity among the features. Correlation analysis between maturity scores and per-variety median vegetation metrics revealed strong to moderate negative correlations with late-season measurements (90 and 120 dpp, *ρ* = -0.84 to -0.65, *p*-value ≤ 1.2×10^−5^, Supp. table S4.11). These findings confirm that vegetation metrics effectively capture the transition from active growth to harvest readiness (senescence), as earlier maturing varieties (with higher maturity scores) enter senescence sooner. Furthermore, gene expression analysis revealed strong co-expression between gene pairs across multiple pathways (Supp. figure S4.4, Supp. table S4.12), highlighting a tightly integrated regulatory network. Within the core pathways, strong correlations were detected among circadian clock components (e.g. *StCO* (Constans) and *StGI2*, *StGI1* and *StTOC1*, *ρ* = −0.85 and 0.84, *p*-value < 10^−16^) and tuberisation genes (e.g. *StCDF1* (Cycling DOF Factor 1) and *StCDF2* (Cycling DOF Factor 2), *StCDF2* and *StFKF1* (*Flavin-binding Kelch Repeat F-Box 1*), *ρ* = 0.88 and −0.84, *p*-value < 10^−16^). Strong co-expression also extended across functional categories, indicating robust linkages between circadian clock and tuberisation genes (e.g. *StFKF1* and *StGI2*, *StCDF1/StCDF2* and *StGI2*, *ρ* = 0.93 and −0.77, *p*-value < 10^−16^) as well as circadian clock and ABA signaling genes (e.g. *StGI1* or *StTOC1 (*Timing of CAB Expression 1) and *StSNRK2* (SNF1 (sucrose non-fermenting 1)-related protein kinase 2), *ρ* = 0.84 and 0.79, *p*-value < 10^−16^). Additionally, gene expression levels showed strong-to-moderate correlations with vegetation metrics measured at 40 and 60 dpp, corresponding to the period of tissue sampling (Figure 4A, Supp. table 4.13). For instance, negative correlations were found for the tuberisation genes *StBEL5* (BEL1-like transcription factor 5) and *StTPL* (Topless) with vegetation metrics at 40 dpp (*ρ* = −0.82 to −0.74 and *ρ* = −0.72 to −0.68, *p*-value < 10^−16^). The observed associations between plant features monitored during the growing season and final yield demonstrate that these features can be used as predictors of plant performance.

**Figure 4.**
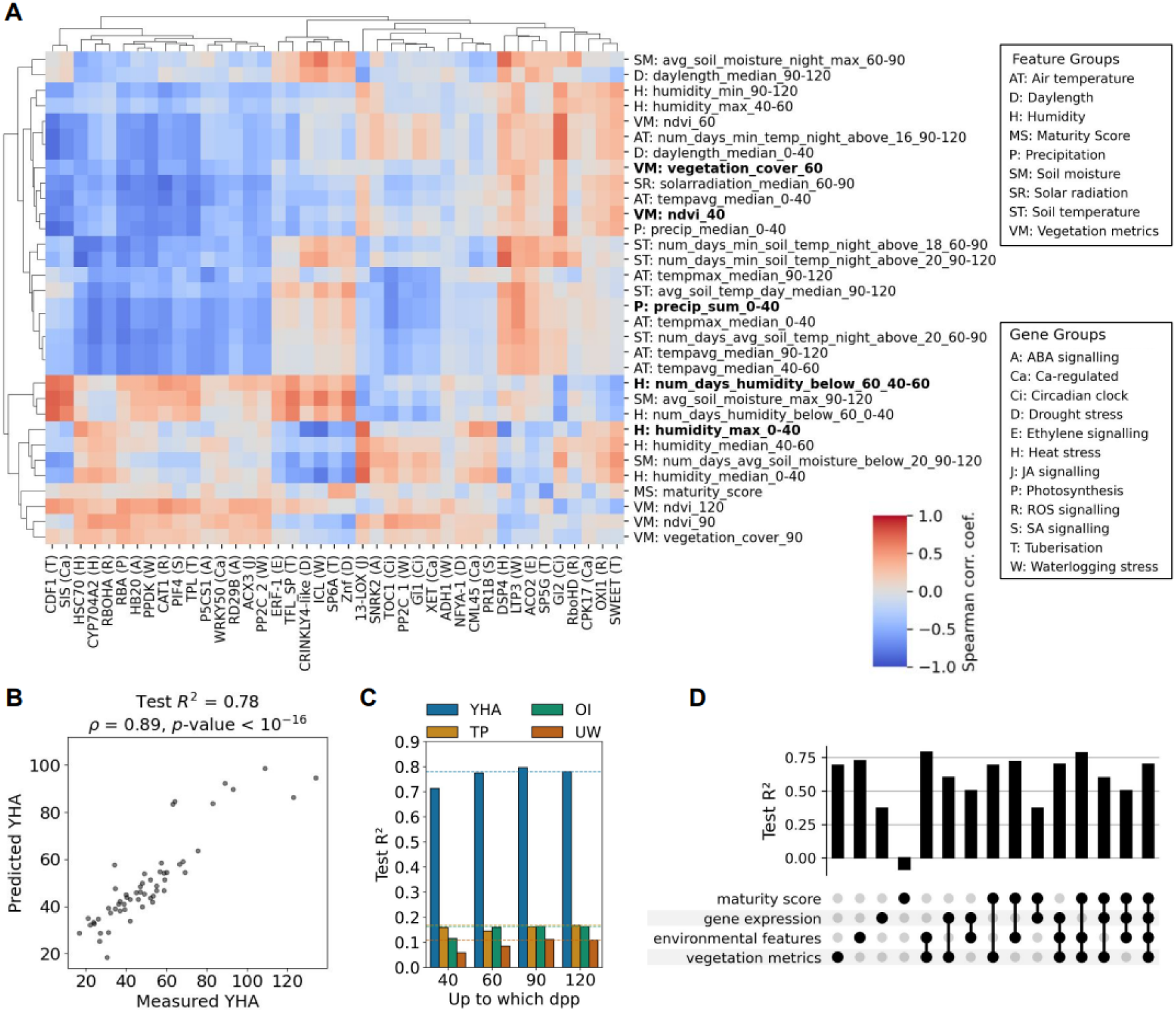
Early-season field data from the first two months of growth enables robust prediction of tuber yield. (A) Heatmap showing Spearman correlation coefficients between the selected set of non-highly correlated environmental/vegetation features and the non-highly correlated genes (*N* = 112, Methods M5). The five features included in the linear regression equation for yield per hectare are highlighted in bold. Acronyms used for feature categories: AT: Air temperature, D: Daylength, H: Humidity, MS: Maturity score, P: Precipitation, SM: Soil moisture, SR: Solar radiation, ST: Soil temperature, and VM: Vegetation metrics. Acronyms used for gene processes annotation: A: Abscisic acid signalling, Ca: Ca-regulated, Ci: Circadian clock, D: Drought stress, E: Ethylene signalling, H: Heat stress, J: Jasmonic acid signalling, P: Photosynthesis, R: ROS signalling, R: SA: Salicylic acid signalling, Se: Senescence, T: Tuberisation, W: Waterlogging stress regulated (Supp. table 1.6). (B) Performance of kernel ridge regression model in predicting tuber yield per hectare (YHA) using vegetation metrics, environmental data, and maturity scores, across the full growing season. The scatter plot shows observed versus predicted YHA values for the test set. (C) Barplot showing the performance of regression models in predicting four tuber yield and quality traits: yield per hectare (YHA, blue), number of tubers per plant (TP, orange), overall impression (OI, green), and underwater weight (UW, red), after training on data subsets corresponding to different time points after planting (Supp. Table S5.2, Methods M6). Models were trained on subsets of selected features available up to 40, 60, 90, or 120 days post planting (dpp) and evaluated on the test set. The dashed horizontal lines indicate the performance with features up to 120 dpp. (D) UpSet plot displaying test *R²* values for kernel ridge regression models predicting yield per hectare (YHA) using different combinations of feature groups.

### 2.5 Tuber yield can be predicted using field data from the first two months of growth

We next applied machine learning (ML) to build models that can predict yield and quality traits from the measured plant growth, environmental and gene expression data (Figure 1). To create a non-redundant feature set and avoid multicollinearity, we filtered the initial features (vegetation metrics, environmental measurements, and maturity score) based on Spearman correlation, where pairs of features with *ρ* > 0.85 were flagged, and manual review was performed to select a final, reduced set of 32 features (Figure 4A, Supp. table S5.1, Supp. Figure S5.1, Methods M5). The same procedure was applied to genes, removing *StCO, StCDF2, StFKF1*, and *StBEL5*, resulting in a subset of 41 genes whose expression is not highly correlated. The set of non-highly correlated features derived from vegetation metrics and environmental data, together with maturity scores, were used as explanatory variables, and tuber yield and quality traits as target variables in a regression framework. To rigorously evaluate whether ML models could accurately predict target outcomes for novel varieties entirely excluded from model tuning, training, and feature selection, we applied the same 90-10% splitting of the balanced dataset into training (40 varieties, 550 plots) and test sets (four varieties, 56 plots), as used in the correlation analysis above (Methods M4). Classical machine learning methods were used, including linear models ^26–28^, tree-based ensembles ^29,30^, and kernel-based methods ^31,32^, identifying kernel ridge regression as the best-performing model across the tested methods (Supp. Figure S5.2).

For predicting yield per hectare, the kernel ridge regression model achieved a coefficient of determination (*R²*) of 0.78 on the test set, with a Spearman correlation of 0.89 between predicted and observed values (*p*-value < 10=^16^, Figure 4B), indicating strong predictive performance. Test set *R²* performance for the rest of the tuber yield and quality targets, namely tubers per plant, overall impression and underwater weight, was 0.17, 0.16 and 0.11, respectively (Figure 4C, Supp. Table S5.2). In leave-one-out experiments per field trial (location and year) or by variety, the mean absolute percentage error for YHA was 36.0 or 16.7, respectively (Supp. figure S5.3). This revealed that the available dataset lacked sufficient geographic diversity to enable reliable generalization of the models to entirely new locations. In contrast, the models generalized well across varieties, with varieties such as Asterix, Flamenco and Desiree achieving mean absolute percentage errors below 6.2.

To next assess how early in the season reliable yield predictions could be made, we divided the training data into four temporal subsets: (i) features available up to 40 days post planting, (ii) up to 60 days, (iii) up to 90 days, and (iv) the full dataset up to 120 days post planting (Methods M6). A kernel ridge regression model was trained separately on each subset and evaluated on the test set (Figure 4D, Supp. Table S5.2). For YHA, the model trained on features available up to 60 dpp achieved a coefficient of determination (*R²*) of 0.77 on the test set (*ρ* = 0.87, *p*-value < 10=^16^) (Figure 4C, Supp. Figure S5.4), which is only a ∼1.3% lower performance compared to the full model with features up to 120 dpp, indicating that yield can be accurately predicted as early as the first two months of the growing season.

Additionally, we evaluated the predictive performance of different combinations of input feature groups for YHA on a subset of varieties with measured gene expression data (Figure 4D, Supp. Table S5.3, Methods M6). This showed that drone-derived vegetation metrics, capturing real-time ‘resource and response’ dynamics during growth, and environmental variables reflecting general location- and weather-related patterns, were the most informative feature groups when comparing models that were trained using a single feature group, achieving *R^2^* ≥ 0.69 (Supp. Table S5.3). Unsurprisingly, combining these two data types yielded the best-performing models (Figure 4B), whereas inclusion of gene expression data did not further improve their prediction accuracy. These results not only demonstrated that highly accurate predictive models can be constructed from drone-derived vegetation metrics and environmental features, but that potato yield can be robustly predicted already from data captured in the first two months of plant growth.

### 2.6 Data-derived ML models facilitate the extraction of predictive equations for potato yield

Finally, we focused on identifying a minimal set of most informative features and on deriving a compact, easily interpretable linear equation capturing the key drivers for yield prediction. To this end, we simplified the ML models for interpretability rather than maximizing overall predictive performance. The Kernel Ridge regression models might have achieved strong predictive performance using data up to 60 days post planting (Figure 4C), but they operate as “black boxes” in a highly complex, transformed feature space, making it difficult to isolate the exact contribution of each individual input feature. By contrast, a sparse Linear Regression model provides direct, quantifiable insight into the relationships between features and the target, as the magnitude and sign of each coefficient indicate the strength and direction of the association.

Thereby, to identify the five most informative features for linear regression modeling, we performed a combinatorial search over the set of selected features (Figure 4A, Methods M6), derived from vegetation metrics and environmental data available up to 60 days post planting, together with maturity scores. Every possible five-feature combination was evaluated using cross-validation on the training data, and the linear regression model achieving the maximal median *R²* was then selected to define the final linear regression equation (Figure 5A). The test set *R²* of the linear regression equation for YHA was 0.72 (Figure 5B), which represents only a modest (∼7.7%) reduction in performance compared to the full kernel ridge model with features up to 120 dpp (Figure 4B). The Spearman correlation between predicted and observed values was 0.82 (*p*-value = 1.3×10=^14^, Figure 5B). The strongest positive influences on YHA were attributed to the environmental features: the number of days humidity was below 60% during the 40–60 dpp period (+15.20), and the maximum humidity observed between 0 and 40 dpp (+10.95) (Figure 5C). The drone-derived vegetation metrics also showed positive associations with yield, with both NDVI at 40 dpp (+5.51) and vegetation cover at 60 dpp (+6.88) contributing as positive predictors, which demonstrated that better early-season plant vigor and greater mid-season vegetation coverage are indicators of higher final yield. Conversely, the cumulative precipitation between 0 and 40 dpp had a negative coefficient (−17.31), indicating that excessive rainfall early in the growing season is associated with decreased yield. Verification of correlations among the features included in the equation revealed no strong pairwise correlations, except for ndvi_40 and vegetation_cover_60 (*ρ* = 0.71, *p*-value < 10 ^16^, Fig. 4B, Supp. Table S6.1), consistent with the selected variables capturing largely complementary predictive information.

**Figure 5.**
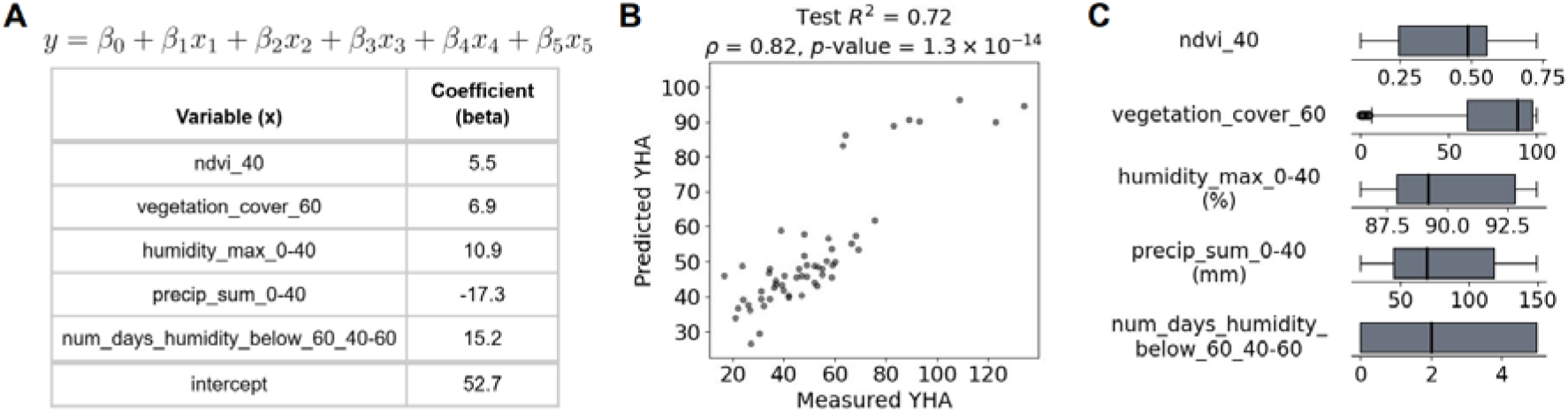
Data-derived ML models facilitate the extraction of predictive equations for potato yield. (A) The linear regression equation for yield per hectare with variables and coefficients provided in tabular format. The variables comprised NDVI at 40 days post planting (dpp), vegetation cover at 60 dpp, maximum humidity between 0 and 40 dpp (%), cumulative precipitation between 0 and 40 dpp (mm), and the number of days with humidity below 60% during the 40–60 dpp period. (B) Performance of a linear regression model trained on the five most informative features in predicting tuber yield per hectare. The scatter plot shows observed versus predicted yield per hectare (YHA) values for the test set. The coefficient of determination (*R²*) of 0.72 and Spearman correlation coefficient of 0.82 (*p*-value = 1.3×10□^14^) between predicted and actual yield per hectare indicate strong predictive performance. (C) Boxplots showing the distribution of the five features used in the linear regression model for yield per hectare on the training dataset.

## 3. Discussion

In the present study, we performed an in-depth analysis of field trial data of a variety of potato cultivars grown under varying growth conditions across 5 European locations in 3 consecutive years (Figure 1A). The different collected data types, namely drone-captured vegetation metrics, environmental variables, gene expression data, and post-harvest tuber yield and quality variables, represent heterogeneous data that individually and jointly provide essential indicators of potato plant development and performance. We demonstrated that the distinct environmental conditions at each field trial location are closely reflected across all collected data types (Figure 2). We also showcase that the relation between yield magnitude and stability can be used to identify resilient, high-yielding varieties across diverse locations and growing conditions (Figure 3). With regard to the quality-related tuber traits, underwater weight, as an indicator of starch content, is a variety-specific trait linked to the intended market use. A lower RSD is desirable across all traits, as it reflects greater stability under varying growth conditions (Figure 3A-D). For yield, varieties positioned toward the upper-left corner of the plot thus exhibit both high median performance and low variability, making them particularly strong candidates for resilience (Figure 3A).

A large proportion of the measured variables showed significant correlations across the data types (*p*-value < 0.05, Figure 4). Notably, drone-captured vegetation metrics, environmental variables and gene expression data were associated with tuber yield and quality. These findings suggest that post-harvest performance can be predicted from plant development and environmental data collected during the growing season ^1,5^. Beyond the pairwise monotonic associations captured by Spearman correlation, unable to reveal more complex dependencies, machine learning (ML) models can leverage non-linear patterns and higher-order feature interactions across heterogeneous data sources ^33^. Therefore, by training ML models on the field trial data, we demonstrated highly accurate predictive performance (*R^2^* ∼ 0.8, Figure 4B) of tuber yield from the vegetation and environmental data. Importantly, we show that predictive performance within ∼1% of the maximum can be achieved using only explanatory data from the first two months of plant growth (Figure 4C). In potato, this generally corresponds to a growth stage up to and including tuber initiation ^34^. Indeed, successful early-prediction results have also been described previously ^35,36^, with vegetation-specific spectral data at the tuber initiation stage found to be more correlated with potato marketable yield than from the later growing season in which plants enter a maturation stage (>3 months post-planting) ^37^. With other crops, early-season crop yield prediction has also been demonstrated across different scales and agricultural contexts, with measurements taken 3-4 months before harvest for annual cereals ^38,39^ and up to eight months before harvest for perennial crops ^40^.

We provide indications that gene expression data also exhibits a strong potential for predicting potato yield (see Figure 4A,D, Supp. table S4.6), despite in our case showing a lower predictive power than vegetation metrics and environmental features. These features were measured multiple times throughout the growing season, whereas gene expression was measured at a single time point. Notably, the gene expression data corroborate the gene-regulatory modules for tuberisation, which position StGI1 and StSP6A together with its proximal regulators, including StCDF1, StFKF1 and StTPL1, as central determinants of tuber yield ^41^. Together with the predictive importance of early canopy development, these findings underscore that early photosynthetic surface establishment, coupled with tuberisation capacity, are key contributors to yield stability under variable conditions ^42^.

By further studying the vegetation and environmental features, which in our dataset were most complete and informative for potato yield prediction, we uncovered that a simple 5-parameter equation could already predict over ∼72% of the variation of tuber yield (Figure 5A), which is within ∼8% of the achievable maximum predictive performance. In our specific case, this entails measuring humidity and precipitation for the first two months of growth, and performing either drone or manual sensor-based measurements of visible light, yielding the NDVI index, and vegetation cover, at 40 and 60 days post planting, respectively (Eq. 1). The inclusion of these variables is in line with previous findings. Both environmental variables, such as precipitation ^43^, as well as vegetation metrics, such as the NDVI and vegetation cover ^1,35,44^, have been found to be highly predictive of potato tuber yield. Nevertheless, as demonstrated by previous studies, accurate prediction is also likely possible with the use of different feature sets ^1,19^, which is known as the ‘Rashomon Effect’ in ML - a phenomenon when models achieve similar performance through multiple, distinct feature sets ^45,46^. Therefore, our approach demonstrates how potato growers and farmers can simply measure a handful of vegetation and environmental variables to achieve early and highly accurate yield predictions. Moreover, using similar or expanded datasets for different crop varieties, it is possible to develop similar or improved predictive models and equations, facilitating on-site, precision-based, digital agriculture ^47,48^.

The field trial data included 5 locations in Europe (Figure 1A), with large variability across environmental conditions (Figure 2A,B). Here we demonstrated how the varying environmental properties between field trials were captured by the measured explanatory data (Figure 2), and resulted in highly accurate predictive models and minimal equations (Figures 4 and 5). This supports the validity and applicability of our approach, also suggesting that the field trial was already of sufficient size for precision agriculture, especially considering the amount of different cultivars tested (Supp. table S1.2: 44). Robust generalizability (i.e. ability to make accurate predictions on entirely new data) of the predictive models across new varieties was demonstrated both by the high model performance on the independent test set, comprising four varieties entirely withheld during model development (Figure 4B), as well as in leave-one-variety-out cross-validation experiments (Supplementary Figure S5.3). On the other hand, generalizability decreased when withholding entire field trials. This suggests that while the models were highly effective at predicting yield within known environments, predicting performance in a completely new field trial remains a greater challenge, which is in accordance with published results ^7,49^. Nevertheless, by performing field trials across a larger set of locations and a wider range of native conditions, more comprehensive datasets can be obtained that improve the effectiveness of data analysis and modelling in relation to the biological variability and technical noise ^50^. This could be further optimized by incorporating additional variables, such as soil properties and other agricultural management practices, which can undoubtedly lead to more accurate models and generalizable insights. For instance, such data could aid in the analysis of post-harvest tuber yield and quality, identifying varieties that exhibit high performance and low environmental variance and correspond to resilient and stable-yielding varieties across the different locations and growing conditions (e.g. building upon the analysis in Figure 3). Moreover, models specific to a group of varieties or even a single variety could be built, yielding further insights into the different mechanisms by which distinct varieties respond to fluctuating growing conditions, and increasing the precision and specificity of the approach, further facilitating precision agriculture ^48^.

The approach developed and presented here benefits crop breeders and farmers in a number of ways. Firstly, it represents a straightforward solution to analyse field trial datasets, build variety-generic predictive models, and obtain informative feature sets and equations (scripts available online, see Methods M7). This results in highly useful information showing which variables are key to follow for accurate early prediction of post-harvest performance. Importantly, if specific vegetation or environmental variables cannot be measured by the crop growers or farmers and substitutions are required, those can likely be replaced with highly correlated variable counterparts from the entire feature set yielding similarly accurate equations. This opens up the possibility to customize and fine-tune the crop yield equations according to one’s measurement capabilities. The approach also presents a solution for the integration of heterogeneous data across different levels, such as plot (drone, yield and gene expression data), field (environmental data) and variety level (maturity scores), as well as for extraction of features from data acquired at different temporal frequencies. Secondly, we show that expensive drone flights after day 60 may not be required as they do not improve yield prediction. Such early predictors of crop performance also have substantial utility for breeders, facilitating the rapid generation of high-value phenotypic inputs for genomic prediction models, thereby enhancing selection accuracy and shortening breeding cycle times. Thirdly, to the best of our knowledge, our study is the first to integrate the analysis of gene expression data in the context of not only potato, but general crop yield prediction and modeling. The development of rapid and reliable methods to directly or indirectly quantify a small subset of key gene expression markers (e.g., via field-deployable molecular or proxy assays) would enable more precise and timely crop management interventions in the future, as increasingly demonstrated by emerging translational plant phenomics and molecular diagnostics approaches ^51^. Finally, the successful models and results that can be obtained with the present capabilities hint at even more detailed and accurate future possibilities, achievable with larger and more diverse field trial datasets.

## 4. Methods

### 4.1 Field trials and data collection

#### Selection of varieties

The principal criteria for selection of potato genotypes for inclusion in the ADAPT trial were as follows: (i) representation of all major market sectors, including fresh/table, processing, and starch cultivars; (ii) inclusion of varieties with documented or anticipated differences in resilience/sensitivity to environmental perturbations; (iii) incorporation of standard control cultivars routinely used in our laboratories for molecular and physiological analyses; and (iv) availability of fully phased, chromosome-scale genome assemblies to facilitate downstream molecular analyses ^52^. An initial panel comprising >60 candidate varieties was compiled. This list was subsequently reduced to 44 genotypes (used in the present study) based primarily on the availability and phytosanitary status of certified seed tuber material. In addition, diploid lines and other genotypes considered likely to introduce ploidy-related outlier effects were excluded to maintain comparability within the predominantly tetraploid germplasm set. All genotypes were autotetraploid (2n=4x=48). Commercial certified seed tuber material was acquired and distributed by Meijer Potato BV, Netherlands.

#### Field trials

Potato field trials were conducted between spring 2021 and autumn 2023 in four European countries across 5 different locations: Netherlands - Zeeland (Meijer), Austria - Fuchsenbigl and Großnondorf (AGES), Spain - Valencia, and Serbia - Vojvodina (HZPC). Approximately half of the trials were conducted under irrigation. In Zeeland, Valencia and Vojvodina, 43-44 varieties were cultivated in duplicate plots (i.e. two distinct sections per variety), yielding up to 88 plots per field trial, while in the locations in Austria, 12-13 varieties were cultivated in quadruple plots, yielding up to 52 plots per field trial. This yielded a total of 16 field trials and 1,165 plots (Supp. Table S1). The trials were designed as randomized blocks with 2 replicated plots per variety, with the exception of Austria, where 4 replicates were used. One single plot comprised either 8 (Valencia and Zeeland), 12 (Vojvodina) or 50 plants (Austria; 2 rows of 25 plants). Trials were performed under conventional farming conditions with flood irrigation (Valencia), sprinkler irrigation (Vojvodina and Austria) and drip irrigation (Zeeland) during the growing season (see Supp. table SM1 for specific details across the different field trials).

#### Plant growth, development and environmental measurements

During the growing season, plant growth and development and environmental parameters were measured with drones, gene expression measurements and sensors, respectively. Drone flights scanning the trial field were executed at 4 distinct flight dates. The exceptions were some of the field trials in Valencia with 3 drone flights and the Austrian field trials, which were limited to a single drone flight (Supp. Tables S1.1, S1.4). From the drone-captured imagery in all field trials, vegetation metrics was derived, including percent of vegetation cover, normalised difference vegetation index (*NDVI*), weighted difference vegetation index (*WDVI*), and chlorophyll index based on *NIR* and *RED* bands (*CIRED*). The indices were computed as *NDVI = (NIR-RED)/(NIR+RED)*, *CIRED = (NIR/RED)-1)*, and WDVI = NIR−a*RED, where *a* is the soil factor. All the indices were calculated as average values per plot. The service providers were Aurea Imaging (Zeeland, Valencia, Vojvodina) and Blickwinkel Agrarconsulting (Austria).

Environmental data were captured by in-field sensors measuring soil temperature (°C) and moisture (%, volumetric water content). The sensors were placed at 10 cm depth, except for Austrian fields where they were placed at 5 cm depth. Sensors used in Zeeland were from AgroExact (NL), one SoilExact sensor and one CropExact sensor, in Valencia and Vojvodina Dacom Terrasen Pro sensors (CropX, NL), and in Austria LSE01-LoRaWAN Soil Moisture & EC Sensor (Dragino, CN). This was complemented with additional data on air temperature, solar radiation, daylength and precipitation from nearby weather stations, obtained from the Visual Crossing database (Weather Data Services, https://www.visualcrossing.com) on 19. 12. 2023 (see Supp. Table SM2 for geographic coordinates).

#### Maturity scores

At a time point when all but the latest maturing varieties are showing senescence; scores are given to plots ranging from 10 to 90. Here, 90 denotes a very early variety showing senescence at the specific time point in the growing season and 10 a variety which is showing no signs of senescence. Median values per variety (from all available data) of vegetation cover on 90 dpp and NDVI on 90 and 120 dpp were used to impute missing maturity scores with linear regression (Supp. table S1.3).

#### Tuber yield and quality measurements

Post-harvest plant performance was assessed via four tuber yield and quality variables. Yield per hectare (YHA, in tons per hectare) and number of tubers per plant (TP) were measured for every plot in all field trials. Underwater weight was measured by taking a 5050 gram sample, measured in air, and measuring the sample under water. Overall impression (OI) is a batch index on a percentage scale, where higher is better, calculated from a combination of tuber characteristics based on expert quality assessment. OI and underwater weight (UW, in grams, indicative of starch content) were measured at the plot level in all field trials except those conducted in Austria, where a single value representing the aggregate of all four plots was recorded per variety.

### 4.2 Data processing

#### Drone data processing

Due to the differences in the frequency and distribution of drone flights across the growing season between the field trials, we included data from drone flight dates closest to 40, 60, 90 and 120 days post planting (dpp) from each field trial (Supp. table S1.4, note that the 0 dpp reference point marks the date of planting). 40 dpp was chosen as the initial time point because the first flight in the majority of trials occurred closer to 40 dpp than to 30 dpp. To ensure a complete feature matrix prior to machine learning model training, imputation was performed as follows: (i) missing data for the drone flight at 120 dpp for the Valencia 2021 non-irrigated trial were imputed with respective variety-specific values from the drone flight at 90 dpp, (ii) missing data for the drone flight at 40 dpp for Valencia 2022 irrigated trial were imputed with respective variety-specific values of the Valencia 2021 irrigated trial, and (iii) missing values for the variety Orchestra in the Valencia 2022 irrigated trial were imputed by the respective median value calculated across all varieties present in that trial.

#### Environmental data processing

Soil temperature and moisture measurements were taken from the planting date or first available measurement until 120 dpp. For missing data in the case of Vojvodina, these were obtained from the Visual Crossing database. The sensor-measured soil moisture values from the Valencia field trials were linearly re-scaled to the same range as other locations, utilizing data from the Visual Crossing database (Supp. Figures SM1, SM2) to estimate the rescaling parameters. Daily and hourly meteorological data from the Visual Crossing database were also taken from the planting date until 120 dpp. To generate predictive environmental features, we segmented the full growing season data into four time windows (0–40, 40–60, 60–90, and 90–120 dpp where the first day is excluded, the last in included) to match the measurement periods of the vegetation metrics. Various summary statistics (e.g. mean, median, maximum, minimum), cumulative and rolling statistics, and counts of days exceeding or falling below specific threshold values from the planting date to the corresponding drone flight date) as well as growing degree days (GDD) and revised GDD ^19,53^ with base temperature 4.5 °C were then calculated for each environmental variable within each window. When feasible, data was also divided into daytime (defined as the period from sunrise to sunset) and nighttime (the interval from the previous day’s sunset to the current day’s sunrise) to create distinct features for the daytime and nighttime period. Note that features for a specific period were not computed for environmental variables from in-field sensors that had no recorded values for that period. This resulted in a total of 191 environmental features (Supp. table S1.5).

### 4.3 Gene expression measurements

#### Field sampling description

Samples from 16 varieties, selected based on diverse phenotypes observed in the first season, were used for gene expression analysis (Supp. table S1.2). In the Valencia trial 2 fully grown leaves halfway exposed to light of four individual plants per plot (distributed over the entire plot, avoiding border plants) were collected and snap frozen in liquid nitrogen. In all other trials four independent leaf discs from the second fully developed leaf (∼50 mg tissue total) were collected using a puncher (diameter 0.8 mm, surface ≈0.5 cm²) and submerged in RNAlater solution (Thermo Fisher Scientific). To ensure comparability across all experiments, sampling was done between 10:00 and 14:00, the most stable in terms of circadian clock changes, once during the growing season - at the stage of tuber initialisation. Two plots were sampled per each cultivar. Gene expression of 50 genes, including 47 targets and 3 housekeeping genes (Supp. table S1.6), was measured for the 16 varieties in a selection of five field trials (128 plots in total, Figure 1B, Supp. table S1.1, Supp. table S1.2).

#### Target gene selection

Target genes (Figure 1F, Supp. table S1.6) for monitoring the activation of different physiological processes were selected based on one of the two criteria: a) known drought, heat and waterlogging stress marker genes selected by experts based on published data ^21^ and b) genes obtained by reanalysis of selected heat and drought, and stress transcriptomics datasets (Supp. table S1.6). Publicly available transcriptomics RNA-seq datasets with the following SRA Study accession numbers were used for the target gene selection: SRP103919, SRP229087, SRP069961, SRP150699 for drought stress, and SRP005965 (SRR122112 and SRR122115) and SRP618560 (SRR35315774- SRR35315782) ^54–57^ for heat stress. For reanalysis of RNA-seq datasets, reads from individual studies were trimmed and quality-filtered using TrimGalore v 0.6.4_dev ^58^ (non- default parameters used: --quality 20, --trim-n) and mapped to potato pan-transcriptome ^59^ using STAR v2.7.5c ^60^ (non-default mapping parameters used: --outFilterMultimapNmax 10, --outFilterMismatchNoverReadLmax 0.02, --quantTranscriptomeBan Singleend). Differential expression analysis was performed in R v3.6.3 using a procedure based on TMM normalisation from the EdgeR package v3.26.8 ^61^ the *voom* function from the limma package v3.40.6 ^62,63^. Differences between the control group and abiotic stress condition were assessed also using a non=paired two=sided Wilcoxon rank=sum test on RNA=seq read counts, with tests only performed when fewer than half of the values were missing in either group. Target genes were selected based on the consistency and specificity of their transcripts’ stress responsiveness across various cultivars.

#### Nanostring measurements

Samples collected as full leaves were homogenised using mortar and pestle in liquid nitrogen. For the Valencia 2021 trial, samples from two separate plots of the same variety were pooled. Samples collected as leaf discs were homogenized in a lysis buffer using a FastPrep-24 homogenizer (MP Biomedicals) for each plot independently. RNA was isolated with MagMAX™ Plant RNA Isolation Kit (Applied Biosystems) using KingFisher Apex (Thermo Fisher Scientific). Total RNA quantity and quality were assessed with a NanoDrop spectrophotometer (Thermo Fisher Scientific) or a Qubit fluorimeter (Thermo Fisher Scientific). 200 ng of RNA was used for gene quantification using nCounter CodeSet chemistry on the nCounter Sprint Profiler System and (both Nanostring Technologies). nCounter CodeSet chemistry utilizes two ∼50 base probes, the color-coded reporter probe and the capture probe, for each mRNA target of interest ^64^. As multiple potato cultivars were analyzed within the study, probes were designed to target the most conserved 100 bp regions of the target transcript while remaining specific to individual transcript/gene to prevent cross-hybridization. In this study, a NanoString multiplex custom CodeSet was designed and synthesized for the above mentioned 50 genes (Figure 1F, Supp. table S1.6,). RNA was first subjected to a hybridization step in a ProFlex PCR System (Applied Biosystems). RNA samples were hybridized at 65 °C for 18 h in a 15 μL volume consisting of 5 μL RNA sample (200 ng), 3 μL of Reporter, 5 μL of hybridization buffer, and 2 μL Capture Probes (all Nanostring Technologies). After hybridization between target mRNA and reporter–capture probe pairs, excess probes were removed, and the probe–target complexes were aligned and immobilized in the nCounter cartridge. The cartridge was then placed in the NanoString nCounter Sprint Profiler System for image acquisition and data processing.

#### Data processing

Raw gene expression counts were subjected to quality control and normalization using nSolver Analysis Software v4.0 (NanoString Technologies) with default settings. Data analysis included background correction followed by normalization to positive controls and housekeeping genes, using the geometric mean to compute normalization factors, in accordance with the manufacturer’s recommendations. Positive controls consisted of six synthetic DNA targets of different concentrations (NanoString Technologies) spiked into the hybridization reaction to assess hybridization efficiency and assay linearity. To account for variability in RNA input and sample quality, the housekeeping genes *Cox*, *Ef-1*, and *Cyclophilin* were selected, already previously being used as reference genes for normalization of potato nCounter gene expression data ^65,66^. Normalized counts from nSolver were used for further analysis. Genes with expression values lower than 50 in all samples were filtered out (genes *CDF3, TFL1*). Then, log_2_ transformation was applied to the expression values. Consequently, the measured gene expression value for each variety was assigned to both corresponding plots.

### 4.4 Datasets

#### Data balancing

To mitigate location bias and redundancy of trials conducted simultaneously in Zeeland (as observed from PCA and tSNE analysis of drone collected vegetation metrics and environmental features, see Figure 2B,C), the most overrepresented location (Figure 1B), a filtered subset of the initial entire dataset with features derived from vegetation metrics and environmental data from seven field trials (606 plots) was selected (Supp. table S1.1). In this balanced dataset, data from only one field trial per year from Zeeland was included, and data from field trials conducted at Fuchsenbigl and Großnondorf were excluded, as they captured only 13 of the total 44 varieties and vegetation metrics were available only for a single drone flight date. This balanced dataset better reflected the diversity of the original data, reduced the risk of overfitting and enhanced the robustness and generalizability of predictive models. A gene expression data subset consisted of data from five out of the seven field trials in the balanced dataset that had additional gene expression data available for 16 varieties (in total 128 plots, Figure 1B, Supp. table S1.1, Supp. table S1.2).

#### Train-test split

We split the balanced dataset into training and test sets based on varieties. Specifically, 10% of the varieties (four varieties: Farida, Musica, Valdivia, Charlotte, 56 plots) were held out as the test set, and the remaining 40 varieties (550 plots) were used for training. Features with zero values in over half of the training set’s field trials were excluded due to being uninformative. This yielded a training dataset of 196 features that was used to conduct the correlation analysis and train the predictive machine learning models. The same partitioning strategy was applied to the gene expression dataset to ensure consistency in variety allocation. Two of the four varieties from the balanced test set were allocated to the gene expression test set (as data was not available for all four), resulting in a training set of 112 plots (12 varieties) and a test set of 16 plots (2 varieties: Farida, Musica).

### 4.5 Statistical data analysis

#### Soil temperature and moisture analysis

To assess differences in soil temperature and soil moisture between irrigated and non-irrigated field trials conducted simultaneously at a specific location (namely Fuchsenbigl and Zeeland, note that in Valencia irrigated and non-irrigated trials were conducted at two distinct times of the year, Supp. table S1.1), we compared daily mean values of soil temperature and soil moisture. Daily means were used instead of hourly measurements for two main reasons: (i) hourly data are highly autocorrelated, with consecutive observations not being statistically independent, which could artificially inflate sample size and significance levels, and (ii) measurement frequencies varied across field sites, and aggregating to daily means standardized temporal resolution, ensuring comparability across locations.

#### Dimensionality reduction

Principal Component Analysis (PCA) was conducted on two distinct datasets: (i) on the entire dataset with all environmental features and (ii) on the subset with gene expression data. For PCA of gene expression values, also loading vectors were derived from the first two principal components. t-distributed Stochastic Neighbor Embedding (t-SNE) with a perplexity set to 30 was performed using vegetation metrics derived features on the entire dataset, with exclusion of the Fuchsenbigl and Großnondorf trials due to having vegetation metrics data only for a single drone flight.

#### Analysis of yield and variety rankings

We computed the weighted median and weighted relative standard deviation (RSD) for each variety across the four tuber yield and quality variables (Figure 1G: YHA, TP, OI and UW). Note that weighting was applied due to the variability of the number of plots per variety among the field trials (Supp. table S1.1). To enable comparison across all four tuber yield and quality variables, we ranked the varieties separately for each trait, with independent rankings for the median values and for the RSD (Supp. figure S3.2). Specifically, varieties were ranked such that the one with the highest median value for a trait received a rank of 1, while the variety with the lowest RSD received a rank of 1. The ranks were then summed per variety, where a lower sum indicates a potentially more stable-performing variety. Additionally, the rank range (difference between maximum and minimum rank) and the standard deviation of ranks per variety were calculated.

#### Analysis of gene expression distributions

To determine if the differences distinguishing high and low performance can be explained by specific signaling/transcriptional module activation, we compared the three highest (Musica, Sound, Acoustic) and three lowest-performing (Agria, Altus, Bartina) varieties with available gene expression data from our rank list. Differences in the distributions of log_2_ expression values between the best and worst performing varieties per gene across all field trials combined were assessed with the Wilcoxon rank-sum test. A differential expression profile was generated by computing the difference between the median log_2_-transformed expression values per gene of the best and worst performing varieties within each field trial.

#### Correlation analysis

For correlation analysis, the Spearman correlation coefficient (*ρ*) is reported and statistical significance was tested using Welch’s t-test. All correlation analyses were based solely on the balanced training dataset to prevent information from the test data influencing further feature selection and model development, thereby avoiding data leakage. Correlation analysis was performed to investigate relationships across three distinct sets of variables: (i) targets vs targets, i.e. tuber yield and quality variables at both the plot level and the variety level (using the median values calculated per variety), (ii) environmental and vegetation features vs. target variables, and (iii) features vs features. As high correlations among features can lead to multicollinearity, negatively impacting model performance, feature pairs with *ρ* > 0.85 were flagged for manual review and selection. First, the selection was done among pairs of highly correlated features of the same category (i.e. environmental features or vegetation metrics) within the same time period. Then, selection was performed among features of the same category across different measurement periods. Finally, features were selected among all of the remaining pairs of highly correlated features by additional expert analysis. This resulted in a set of 32 features derived from environmental data, vegetation metrics and maturity score (Supplementary table S5.1), where the highest pairwise correlation among the features was lower than 0.85. For gene expression data, removing *CO, CDF2, FKF1*, and *BEL5* resulted in a subset of 41 non-highly correlated (i.e. with pairwise *ρ* < 0.85) genes.

### 4.6 Machine learning

#### Data pre-processing

Preprocessing of the feature matrix, involving specific transformations for distinct feature groups, was integrated directly into the modeling workflow. Crucially, to prevent data leakage, the necessary parameters for these transformations (e.g., mean, standard deviation, or scaling boundaries) were always estimated exclusively on the current training set or cross-validation fold and subsequently applied to the corresponding test set or validation fold. For vegetation metrics features, +1 was first added to all values to ensure all values were strictly positive, then Box-Cox transformation was applied to achieve a more Gaussian-like distribution, followed by standardization. Environmental features were standardized. Maturity score was scaled to the range [-1, 1], assuming a minimum possible value of 25 and a maximum possible value of 100. The target variables were standardized, except for the final linear regression model, for which the original scale was retained to allow direct interpretation of regression coefficients.

#### Model class selection and hyperparameter tuning

Regression models were developed to predict tuber yield and quality variables at the plot level, with model performance assessed using the coefficient of determination (R²). Hyperparameter tuning and model selection were conducted on the training dataset using nested cross-validation, with 5-fold splits at the variety level in both the inner and outer loops. Models were trained using the set of 32 non-highly correlated features derived from vegetation and environmental data. A diverse range of regression model classes was initially evaluated for YHA, including linear models (linear, Ridge, Lasso, and Elastic Net regression) ^26–28^, kernel-based methods (support vector regression and kernel ridge regression) ^31,32^, and, tree-based ensembles (random forest and XGBoost) ^29,30^. In the inner cross-validation loops, each model’s hyperparameters were optimized via grid search (Supp. Table SM3). The set of hyperparameters yielding the highest median R² across the majority of inner folds was then selected for evaluation in the outer loops. Kernel Ridge Regression achieved the highest median R² for predicting YHA and was therefore selected as the final model class for all targets (Supp. figure S5.2). The final kernel ridge regression model for each target utilized the best hyperparameters determined during the inner cross-validation phase. For all other targets (e.g., TP, OI, UW), the hyperparameters for the final kernel ridge model were determined on the inner cross-validation loops for the respective target, as for YHA (Supp. Table SM4). The specific hyperparameters chosen for each target model are provided in Supplementary Table SM3.

#### Model performance analysis

The final predictions for all tuber yield and quality targets were generated using kernel ridge regression models. Each kernel ridge regression model, configured with its target-specific optimal hyperparameters (Supplementary Table SM3), was trained on the complete training dataset using the 32 non-highly correlated features collected throughout the full growing season (up to 120 dpp). Performance validation was conducted on the test set. To next assess how early in the season reliable predictions could be made, the set of 32 selected features was partitioned into four cumulative temporal subsets based on the maximum dpp of the included data points: (i) features available up to 40 days post planting, (ii) up to 60 days, (iii) up to 90 days, and (iv) up to 120 days post planting (i.e. all 32 features). For each target variable, kernel ridge regression models were trained separately on each temporal subset and evaluated on the test set. The predictive capacity for YHA was systematically evaluated by testing all possible combinations of the four primary feature groups: maturity score, vegetation metrics, environmental features, and the expression data of the 41 non-highly correlated genes (Supp. table S5.1, Supp. table S5.3). Kernel ridge regression models were fitted using these exhaustive feature combinations. Recognizing the constraint of available gene expression data, both the training and evaluation steps were restricted to the corresponding subset of samples that possessed all gene expression measurements.

#### Error analysis

To evaluate the generalizability of our predictive kernel ridge regression models with the 32 selected features to new field trials and new varieties, we implemented two distinct cross-validation schemes for predicting YHA. For each iteration of the leave-one-field-trial-out cross-validation scheme, data from one field trial were held out as the validation fold and the remaining as training fold. For each iteration of the leave-one-variety-out cross-validation, data from one variety were held out as the validation fold and the remaining as training fold. For every sample in the validation folds, the Absolute Percentage Error (APE) was calculated to quantify the deviation between the actual and predicted YHA. From the APEs from each cross-validation scheme separately, mean APEs were calculated for all samples and per field trial/variety.

#### Feature selection and linear equations

Previous experiments in which model performance was assessed using subsets of features available up to a certain dpp showed that for YHA, the model trained on features available up to 60 dpp achieved only slightly worse performance as the model trained with features collected throughout the full growing season (i.e. with features up to 120 dpp). Thus, to identify the optimal subset of five features from the 16 features available up to 60 dpp that (a) most strongly determine the potato yield and (b) for predictive modeling of YHA, we employed a feature selection method based on combinatorial search. All possible combinations of 5 features from this set of 16 were generated and for each five feature subset, a linear regression model was trained and evaluated on the training dataset using a five fold cross-validation scheme based on variety. The feature subset that maximized the median *R^2^* score was selected as the final five feature set. Then, the linear regression model with the final five feature subset was fitted to the whole training dataset, the model performance was evaluated on the test set and the coefficients of the model were extracted to derive the linear equation for calculating YHA.

### 4.7 Software and code

#### Software and code

Python v3.8.16 (www.python.org) and R v4.0.4 (www.r-project.org) were used for computations. For general data analysis and visualisation, the Python packages Pandas v1.5.2, Numpy v1.23.5 ^67^, Matplotlib v3.6.2 ^68^, Seaborn v0.12.2 ^69^, Upsetplot v0.9.0, Rasterio v1.3.7, Fiona v1.9.4, Geopandas v0.13.2, pyproj v3.5.0, and adjusttext v1.2.0 were used. For statistical hypothesis testing, Scipy v1.9.3 ^70^ and Statsmodels v0.14.0 ^71^ were used with default settings. All statistical tests were two-tailed. Xgboost v2.1.1 ^30^ and Scikit-learn v1.3.2 ^72^ were used for machine learning, the latter was also utilized for PCA and t-SNE. Code to reproduce the results and analysis, including data preprocessing, dataset generation, statistical data analysis and machine learning is available at https://github.com/NIB-SI/Adapt_field_ML.

#### Data

The field trial dataset created in the present study is available at https://zenodo.org/records/xxx. All results are shared within the manuscript and supplementary material, with additional data and models available at https://github.com/NIB-SI/Adapt_field_ML.

## Acknowledgements

We thank Rik Nuijten for critical comments, Carissa Bleker and Valentina Levak for technical support. The study was supported by the Slovenian Research and Innovation Agency (ARIS) grants no. J2-3060 (J.Z.), P4-0165 (K.G.) and IO-0004, the Public Scholarship, Development, Disability and Maintenance Fund of the Republic of Slovenia grant no. 11013-9/2021-2 (J.Z.), and the European Union Horizon 2020 Research Innovation Action project ADAPT, G.A. 862858 (https://adapt.univie.ac.at/).

## Author contributions

GH, AZ, RG, II, SP, CB, MT, BD, AR, JZ and KG conceptualized the study; MK, MP, GH, JS, PJL, JB, RG, II, SP, CB, MT, BD, AR and KG designed the experimental analysis; MZ, MK, KS, BW, MAS, II, SP, CB, BD performed the experimental analysis; AV, MZ, MK, AZ, II, BD, AR, JZ and KG designed the computational analysis; AV, MZ, ET, MK, MP and JZ performed the computational analysis; AV, II, BD, AR, JZ and KG interpreted the results; AV, JZ and KG wrote the draft; All authors contributed to the final manuscript.

