## Supplementary information for "Potato yield can be predicted using drone-captured and environmental measurements early in the growing season"

[8] (current) 't Laandje (The Little Lane), Haarsterweg 20, 9363 VD Marum, The Netherlands

[9] (current) Enza Zaden, Haling 1-E, 1602 DB Enkhuizen, The Netherlands

[10] Solynta, Dreijenlaan 2, 6703 HA Wageningen, The Netherlands

[11] Wageningen University & Research (WUR), Droevendaalsesteeg 4, 6708 PB Wageningen, The Netherlands

[12] Austrian Agency for Health and Food Safety (AGES), Spargelfeldstraße 191, 1220 Vienna, Austria

#### Table of contents

Supplementary figures … p.2 - p.14

Supplementary tables … p.15 - p.79

### Supplementary figures

#### Section 2

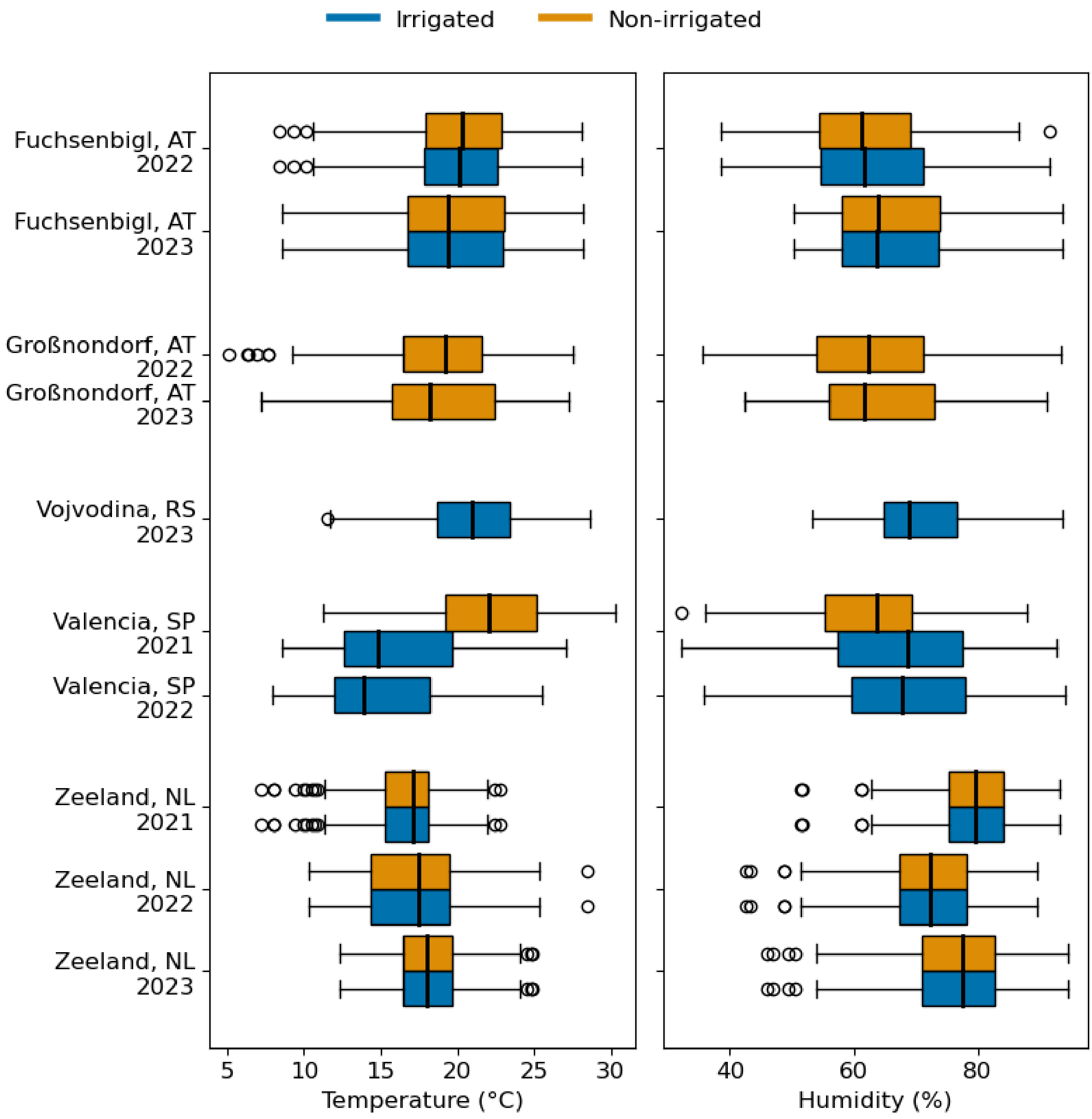


Supplementary Figure S2.1: Air temperature (°C, left panel) and humidity (%, right panel) across field trials. Blue boxes represent irrigated plots, while orange boxes represent non-irrigated plots.

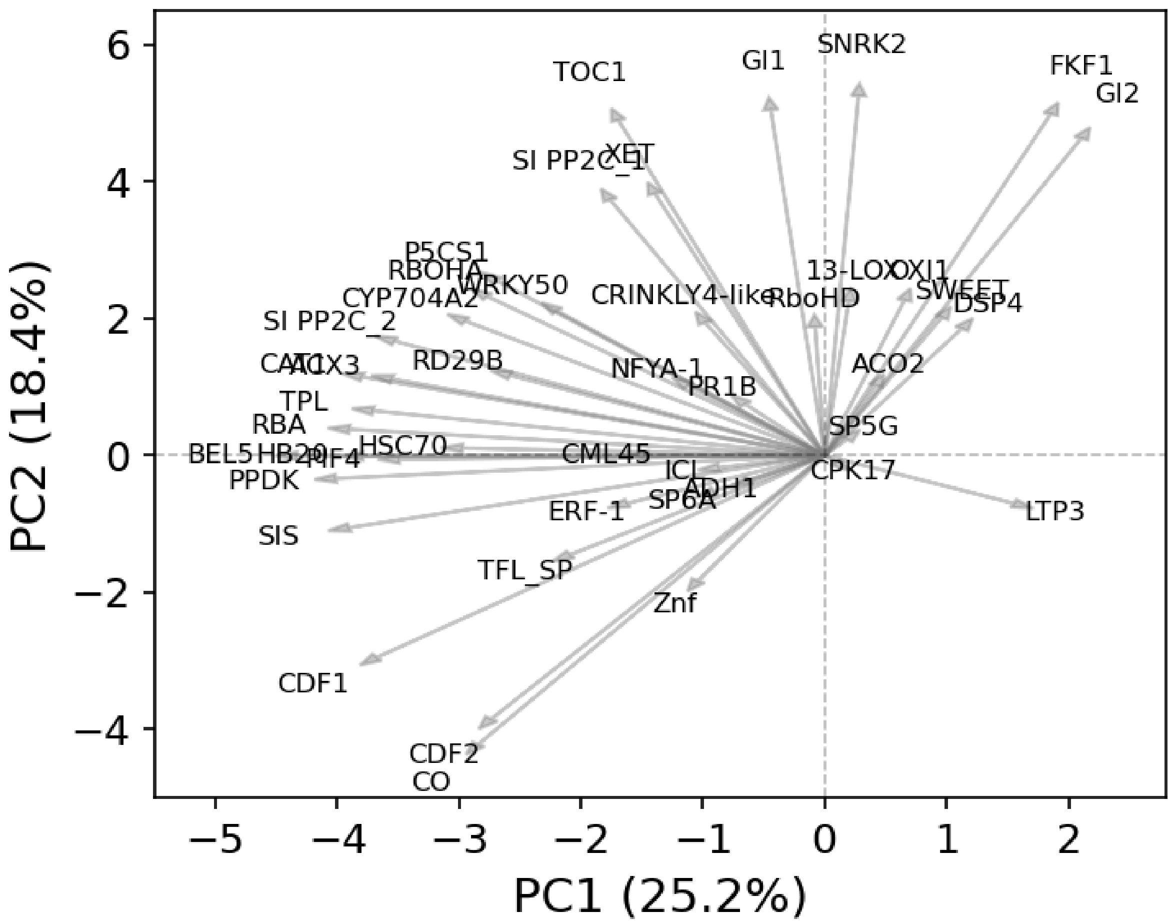


Supplementary Figure S2.2: Principal component analysis (PCA) biplot showing gene loadings on the first two principal components. Each arrow represents a gene, and its direction and length indicate the contribution of that gene to the corresponding principal components (PC1 and PC2). The plot highlights how genes vary in their influence on the main axes of variation across samples.

#### Section 3

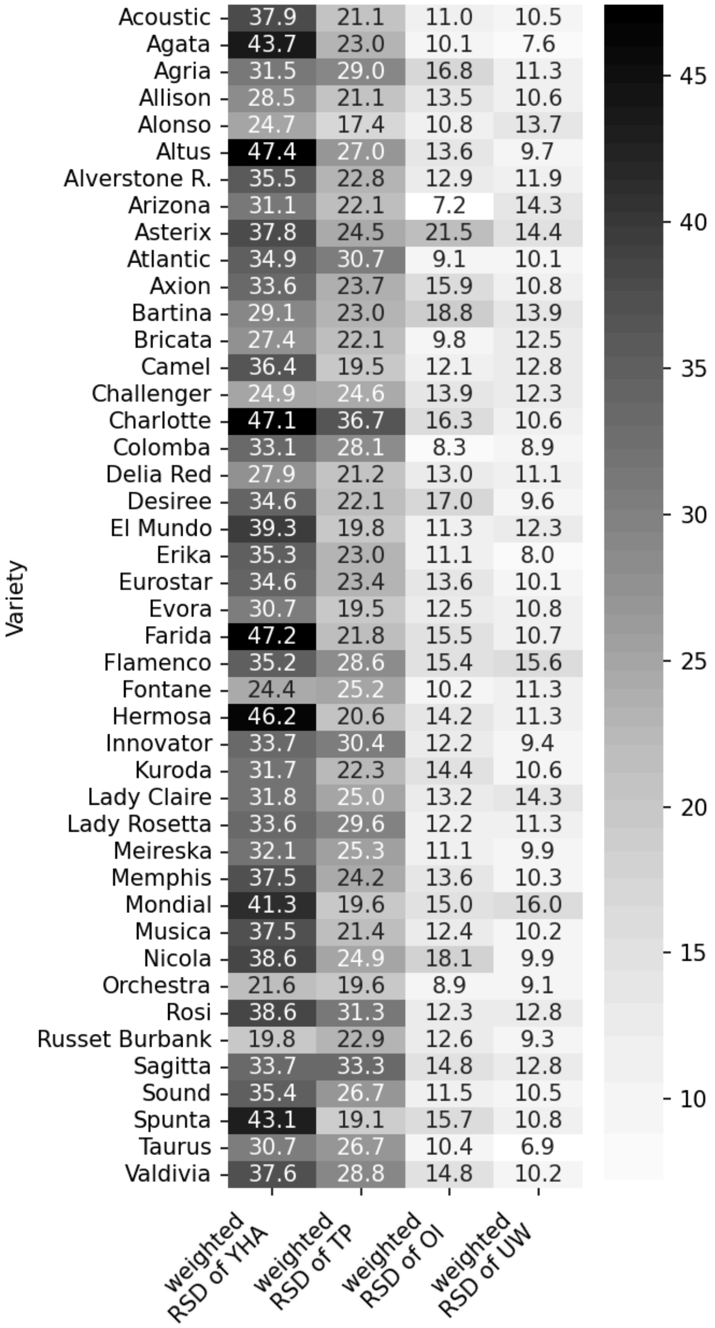


Supplementary Figure S3.1: Heatmap of relative standard deviations (RSDs) of tuber yield and quality variables per field trial: yield per hectare (YHA), number of tubers per plant (TP), overall impression (OI), and underwater weight (UW).

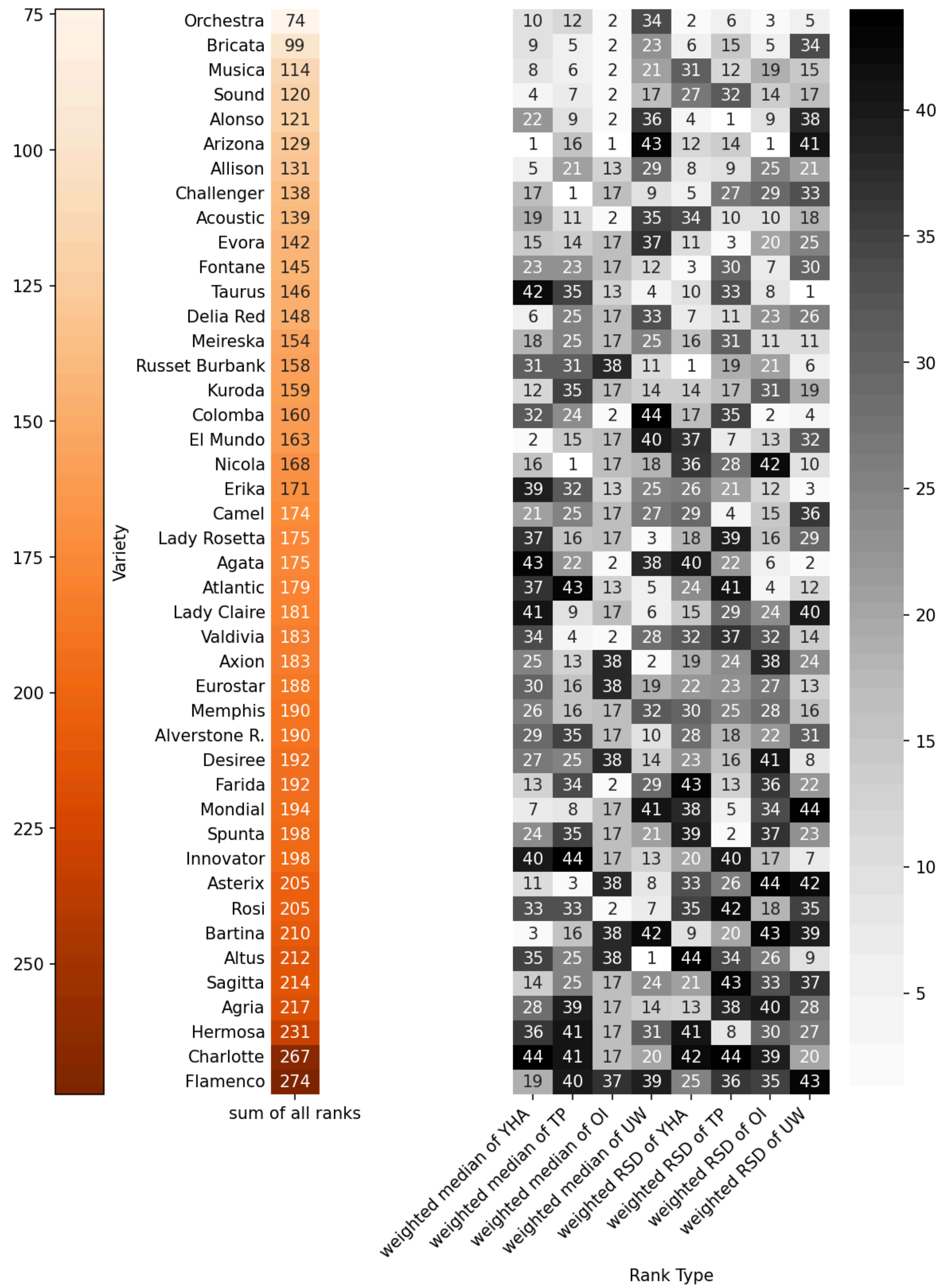


Supplementary Figure S3.2: Heatmap of ranks of medians and relative standard deviations (RSDs) of tuber yield and quality variables per field trial: yield per hectare (YHA), number of tubers per plant (TP), overall impression (OI), and underwater weight (UW). Varieties were ranked such that the one with the highest median value for a trait received a rank of 1, while in the case of the RSD of a trait, the variety with the lowest RSD received a rank of 1.

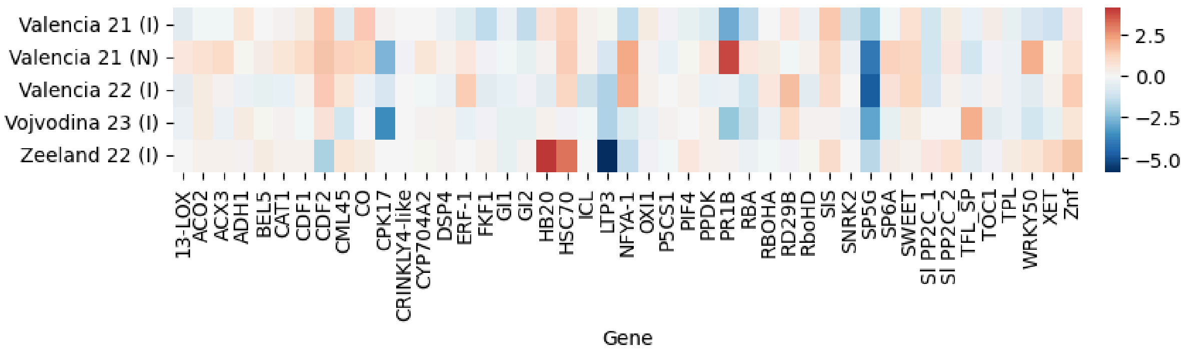


Supplementary Figure S3.3: Heatmap of difference between the median $log_2$-transformed expression values of the best and worst performing varieties within each field trial, with (I) denoting irrigated and (N) denoting non-irrigated conditions. Red indicates higher expression in best performing varieties and blue indicates higher expression in worse performing varieties.

#### Section 4

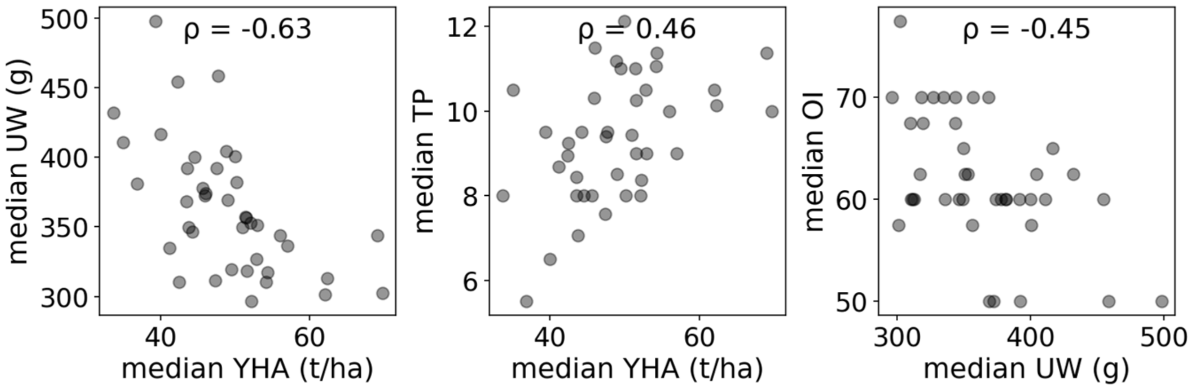


Supplementary Figure S4.1: Scatter plots (N=40) of median tuber yield and quality variables (aggregated by variety) with marked Spearman correlation coefficients (ρ). The top plot shows median yield per hectare (YHA) vs. median underwater weight (UW, $\rho = -0.63$, p-value = $1.5\times10^{-5}$), the middle plot shows median YHA vs. the median number of tubers per plant (TP, $\rho = 0.46$, p-value = $2.9\times10^{-3}$), and the bottom plot shows median UW vs. median overall impression (OI, $\rho = 0.45$, p-value = $3.7\times10^{-3}$).

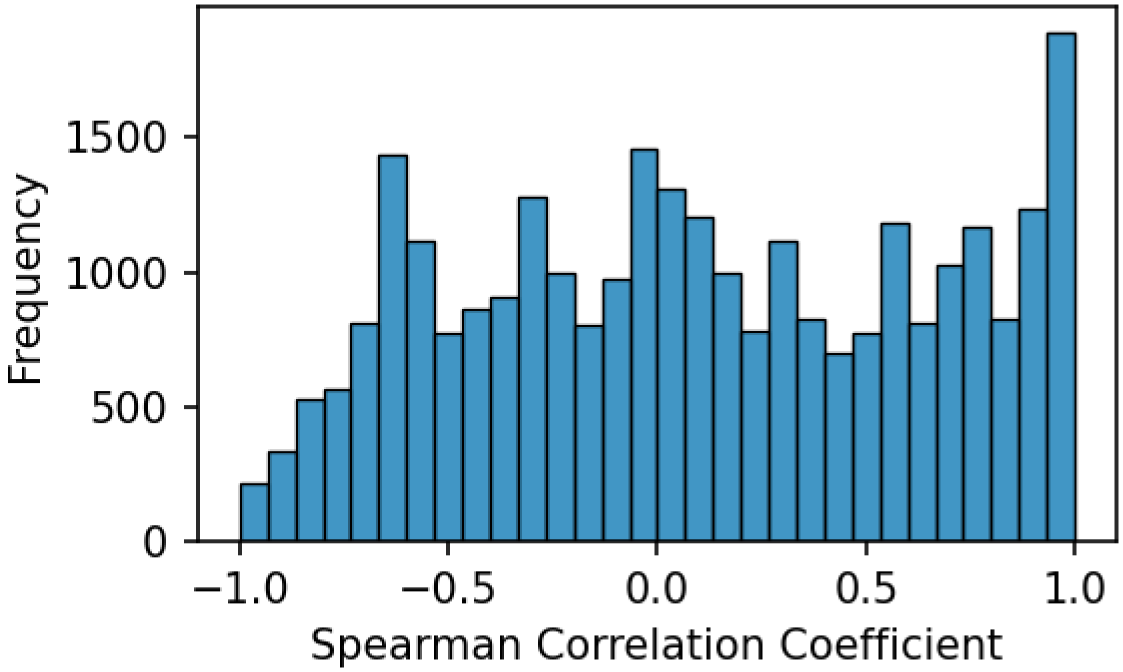


Supplementary Figure S4.2: Histogram of Spearman correlation coefficients between all feature pairs of the training dataset, excluding targets-yield variables.

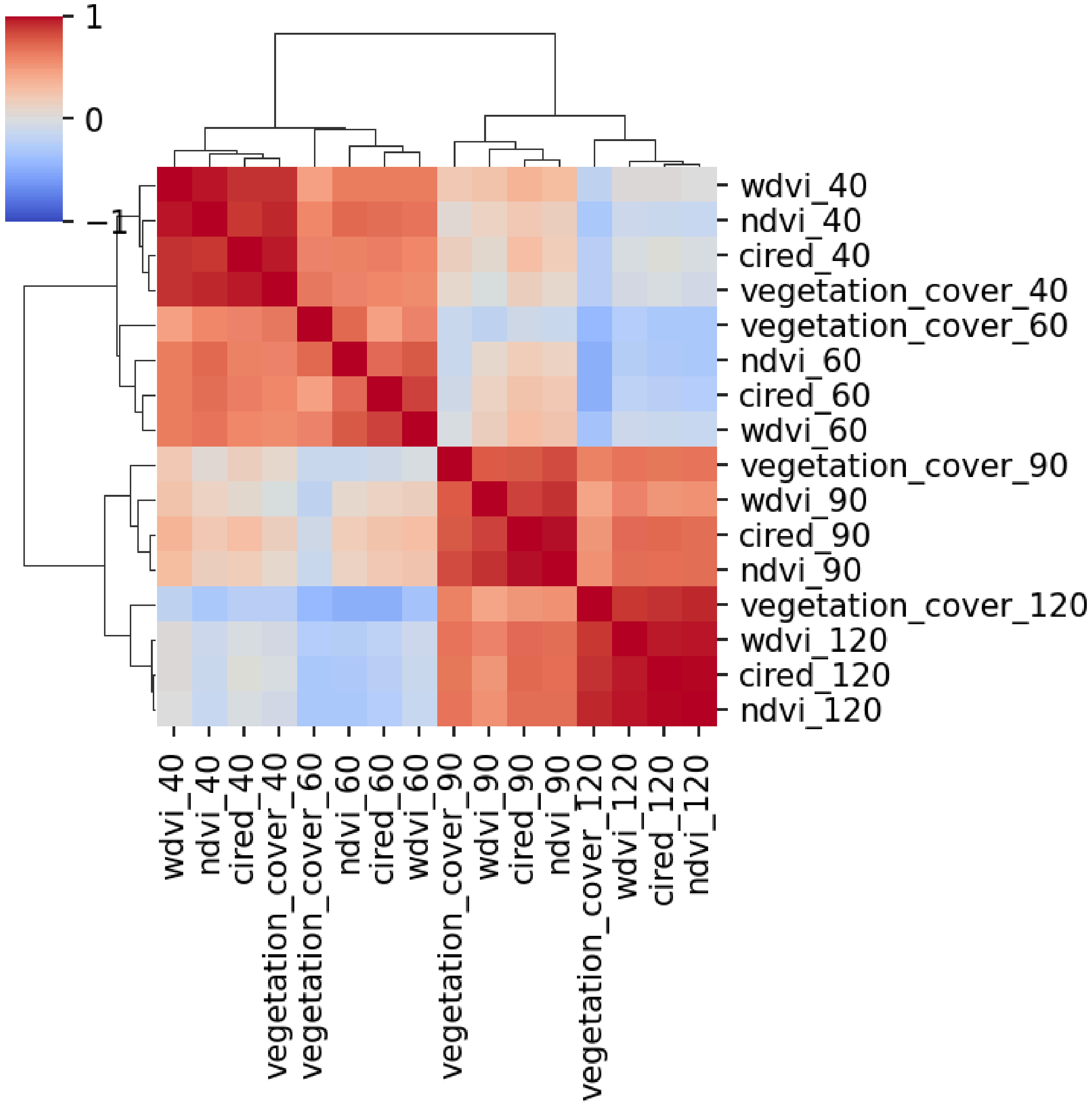


Supplementary Figure S4.3: Heatmap of Spearman correlation coefficients between features derived from drone-measured vegetation metrics on the training dataset.

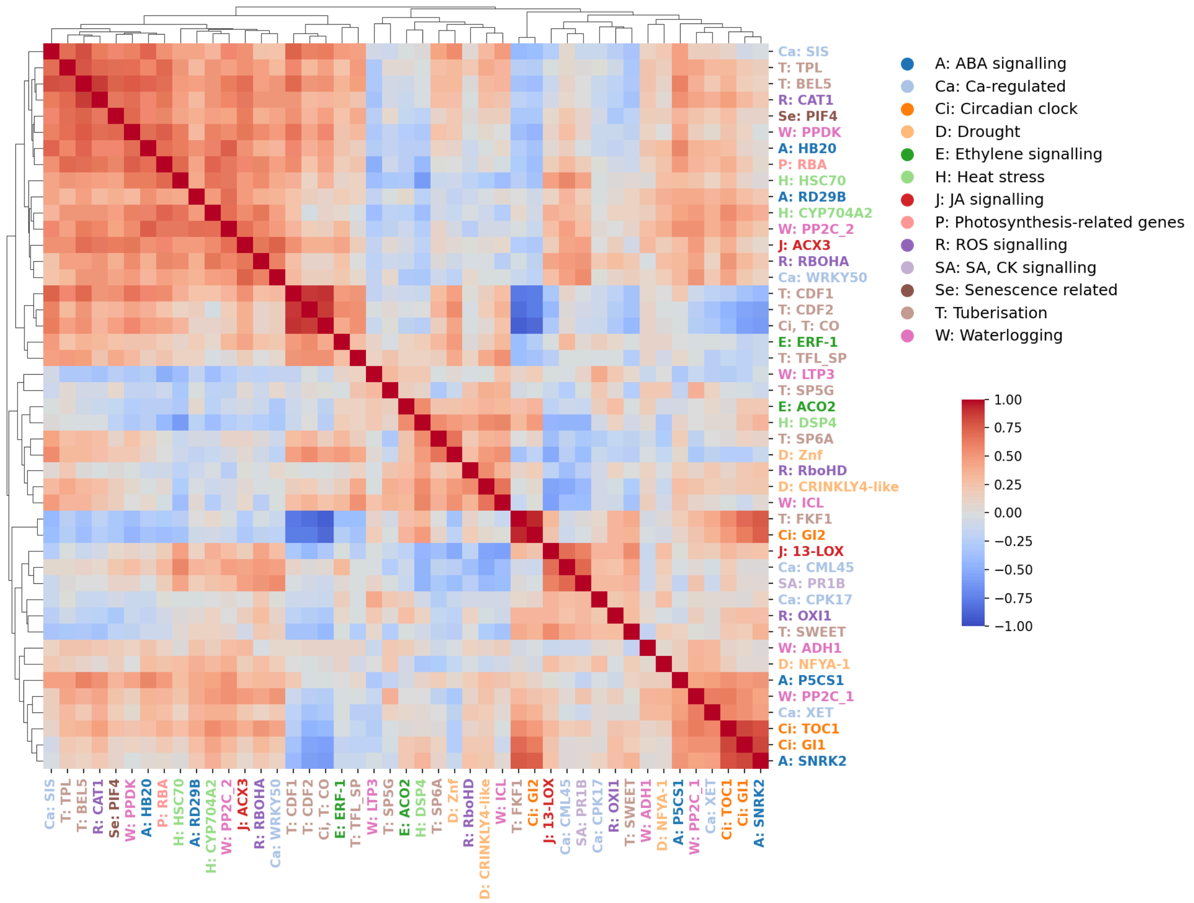


Supplementary Figure S4.4: Heatmap of Spearman correlation coefficients between gene expression values on the training dataset.

### Section 5

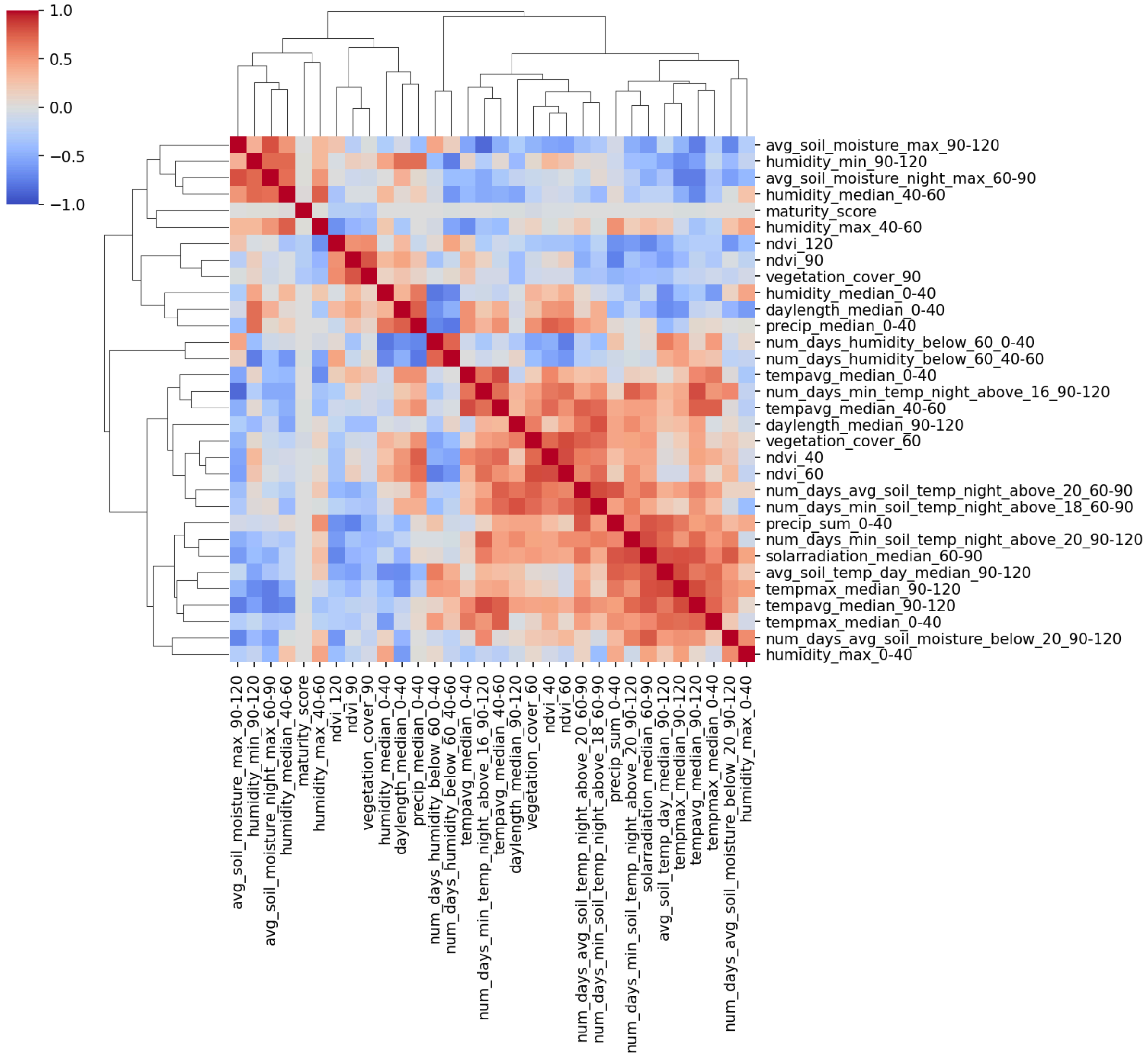


Supplementary Figure S5.1: Heatmap of correlation (Spearman) between selected features on training set.

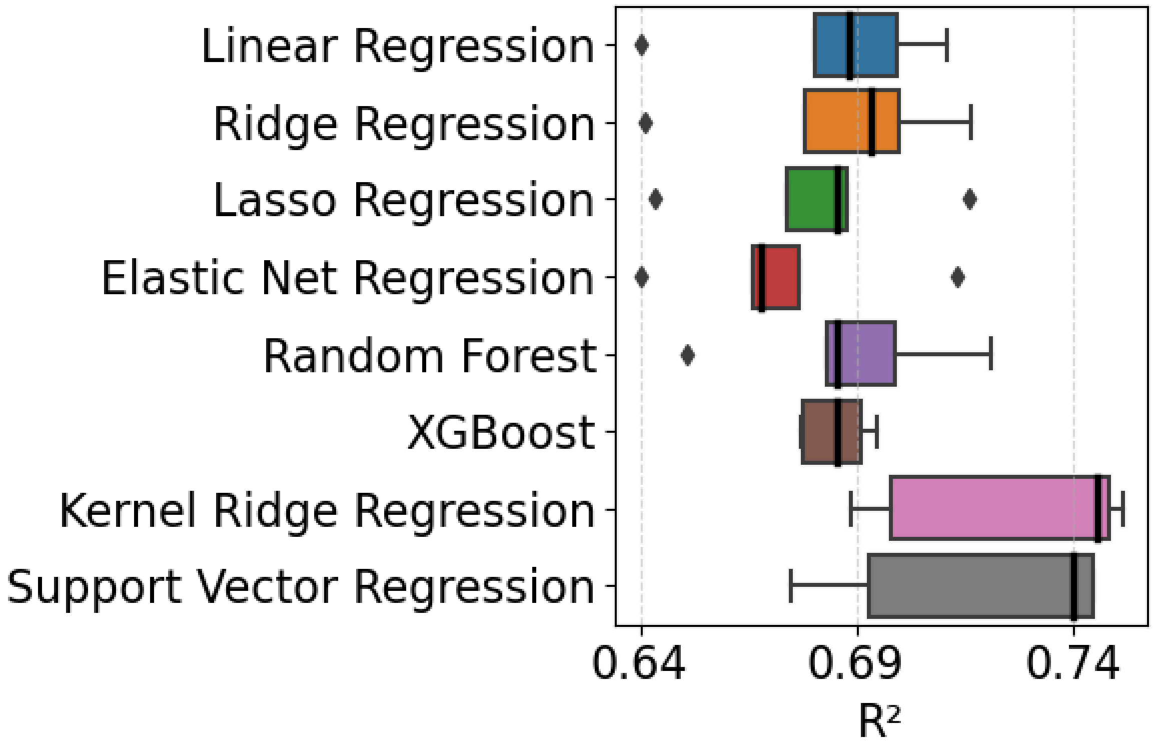


Supplementary Figure S5.2: Performance of different models (evaluated on the outer loops of nested cross-validation for YHA with hyperparameters selected as best in the inner loops of nested cross-validation (on training set).

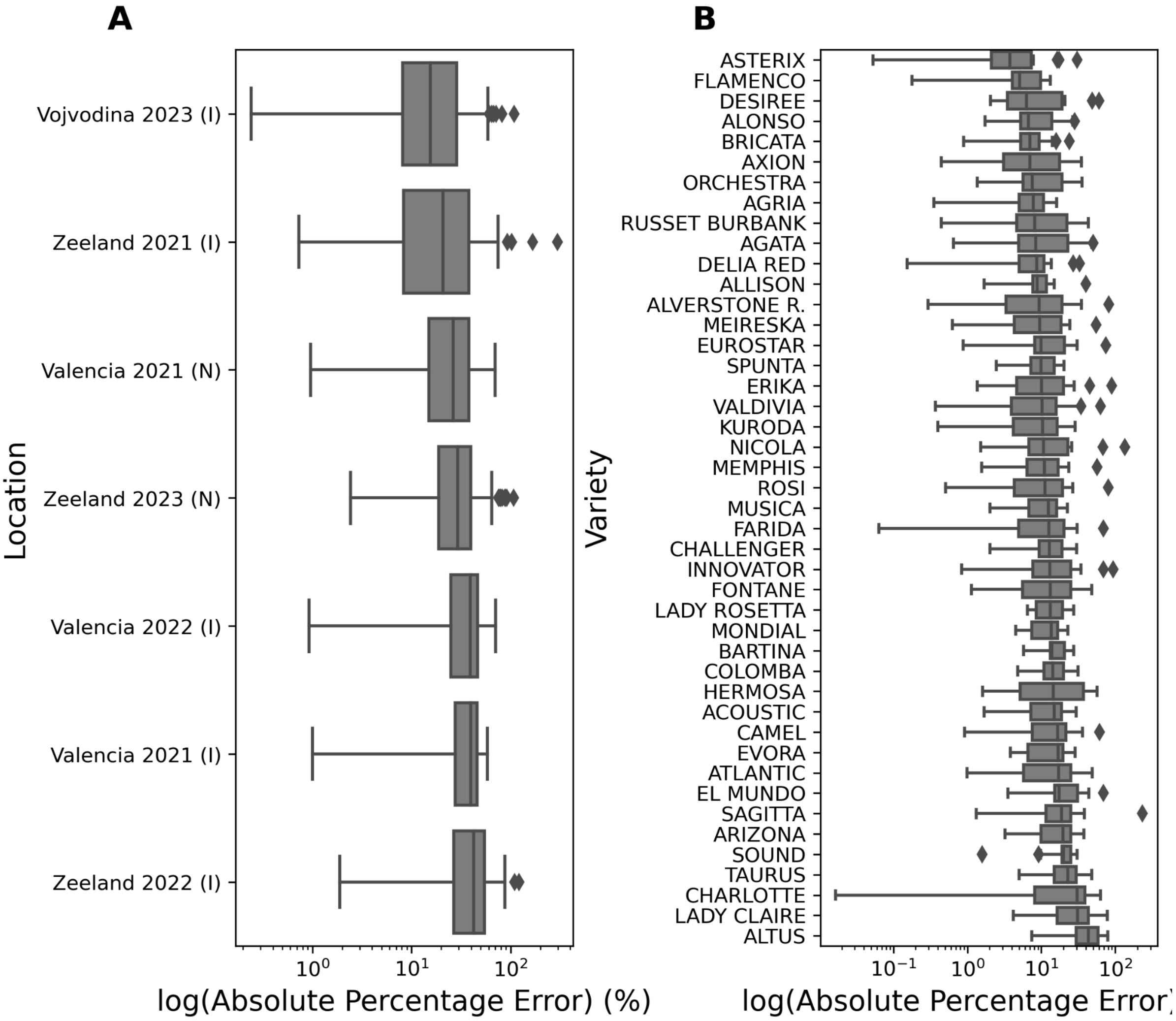


Supplementary Figure S5.3: Boxplots showing the distribution of absolute percentage errors (APE) of the validation folds for kernel ridge regression models evaluated in a (A) leave-one-field-trial-out or (B) leave-one-variety-out cross-validation setting. For field trials, (I) denotes irrigated and (N) denotes non-irrigated conditions.

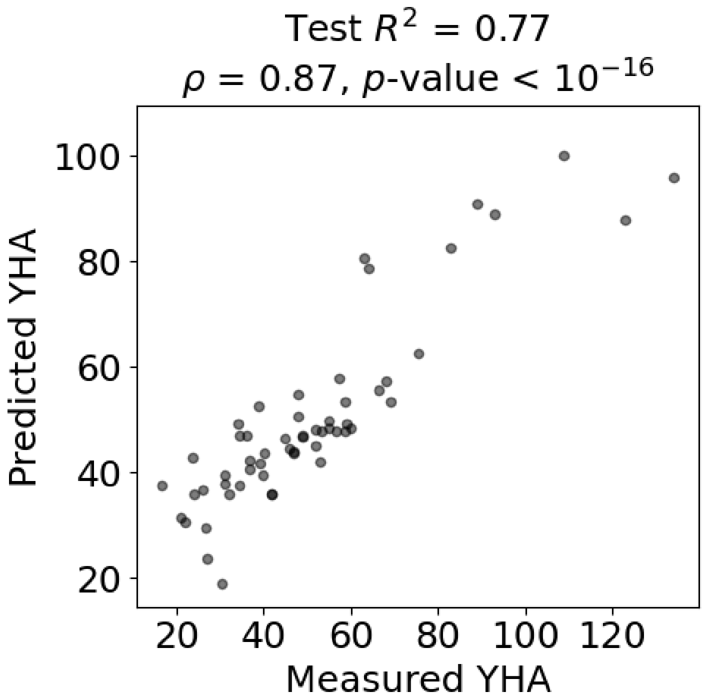


Supplementary Figure S5.4: Performance of kernel ridge regression model in predicting tuber yield per hectare (YHA) from features available up to day 60 post planting. The scatter plot shows observed versus predicted yield values for the test set. The coefficient of determination ($R^2$) of 0.77 and Spearman correlation coefficient 0.87 (p-value < $10^{-}16$) between predicted and actual yields indicate strong predictive performance.

### Methods

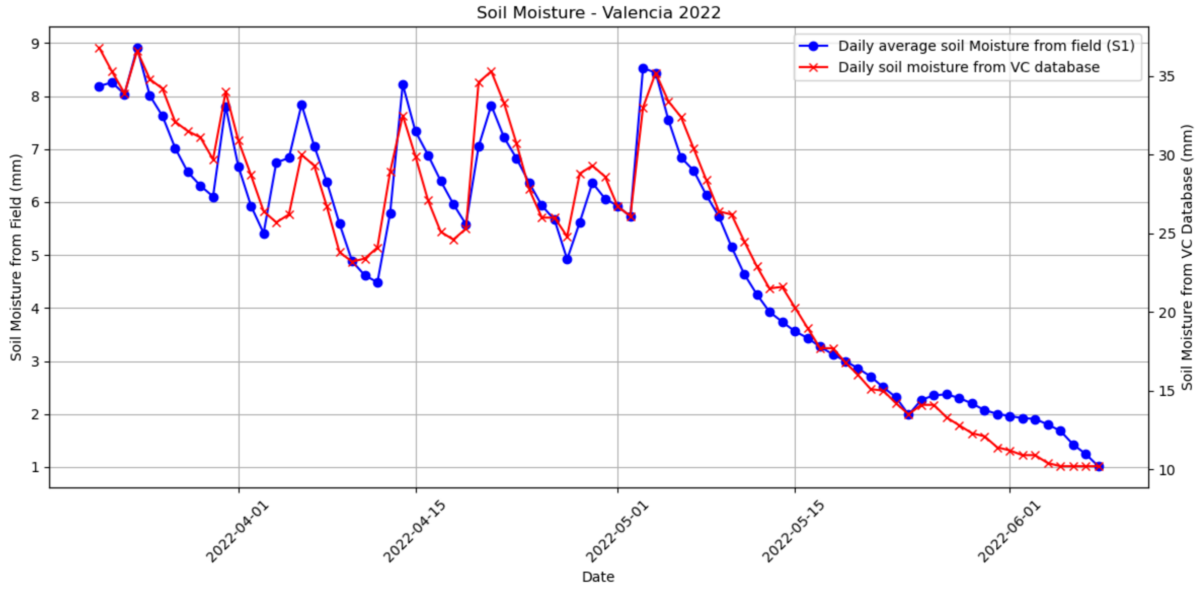


Supplementary Figure SM1: Daily average soil moisture measured by the in-field sensor on the irrigated field trial in Valencia in 2022 (blue) and from the Visual Crossing Weather Data Services database (red).

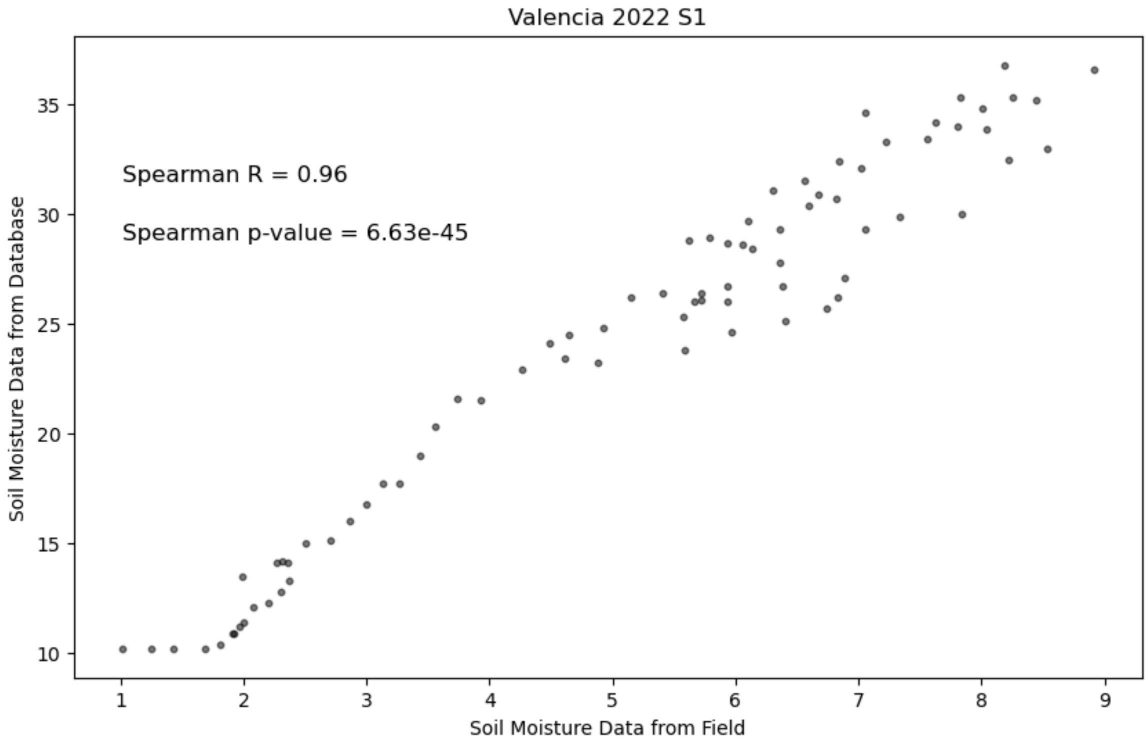


Supplementary Figure SM2: Correlation between daily average soil moisture measured by the in-field sensor on the irrigated field trial in Valencia in 2022 (blue) and from the Visual Crossing Weather Data Services database (red) measured on the same date.

### Supplementary tables

#### Section 1

Supplementary table S1.1: Information about potato field trials.

| Location | Year | Planting date | Haulm killing date | Harvest date | Irrigated | Partner | Included in balanced dataset | Included in gene expression dataset | Number of varieties | Number of plots | Number of drone flyby dates |
|---|---|---|---|---|---|---|---|---|---|---|---|
| Valencia, Spain | 2021 | 3.02.2021 | 15.06.2021 | 15.06.2021 | yes | HZPC | yes | yes | 43 | 86 | 4 |
| Valencia, Spain | 2021 | 14.04.2021 | 27.07.2021 | 27.07.2021 | no | HZPC | yes | yes | 43 | 86 | 3 |
| Valencia, Spain | 2022 | 28.01.2022 | 9.06.2022 | 9.06.2022 | yes | HZPC | yes | yes | 44 | 86 | 3 |
| Vojvodina, Serbia | 2023 | 28.04.2023 | 25.08.2023 | 13.09.2023 | yes | HZPC | yes | yes | 44 | 88 | 4 |
| Zeeland, the Netherlands | 2021 | 3.05.2021 | 20.09.2021 | 27.09.2021 | yes | Meijer | yes | no | 44 | 88 | 4 |
| Zeeland, the Netherlands | 2021 | 3.05.2021 | 20.09.2021 | 27.09.2021 | no | Meijer | no | no | 44 | 88 | 4 |

| | | | | | | | | | | | |
|---|---|---|---|---|---|---|---|---|---|---|---|
| Zeeland, the Netherlands | 2022 | 2.05.2022 | 26.08.2022 | 11.10.2022 | yes | Meijer | yes | yes | 44 | 84 | 4 |
| Zeeland, the Netherlands | 2022 | 2.05.2022 | 26.08.2022 | 11.10.2022 | no | Meijer | no | no | 44 | 84 | 4 |
| Zeeland, the Netherlands | 2023 | 22.05.2023 | 12.09.2023 | 10.10.2023 | yes | Meijer | no | no | 44 | 88 | 4 |
| Zeeland, the Netherlands | 2023 | 22.05.2023 | 12.09.2023 | 10.10.2023 | no | Meijer | yes | no | 44 | 88 | 4 |
| Fuchsenbigl, Austria | 2022 | 21.04.2022 | 13.09.2022 | 13.09.2022 | yes | AGES | no | no | 12 | 48 | 1 |
| Fuchsenbigl, Austria | 2022 | 21.04.2022 | 6.09.2022 | 6.09.2022 | no | AGES | no | no | 12 | 48 | 1 |
| Fuchsenbigl, Austria | 2023 | 26.04.2023 | 7.09.2023 | 7.09.2023 | yes | AGES | no | no | 13 | 52 | 1 |
| Fuchsenbigl, Austria | 2023 | 26.04.2023 | 4.09.2023 | 4.09.2023 | no | AGES | no | no | 13 | 52 | 1 |
| Großnondorf, Austria | 2022 | 14.04.2022 | 6.09.2022 | 6.09.2022 | no | AGES | no | no | 12 | 48 | 1 |
| Großnondorf, Austria | 2023 | 26.04.2023 | 6.09.2023 | 6.09.2023 | no | AGES | no | no | 13 | 52 | 1 |

Supplementary table S1.2: Overview of potato varieties included in the field trials. Varieties marked with asterisk (*) have associated gene expression data available from the respective trial.

| **Variety** | **Zeeland 2021 (irrigated)** | **Zeeland 2021 (non-irrigated)** | **Zeeland 2022 (irrigated)** | **Zeeland 2022 (non-irrigated)** | **Zeeland 2023 (irrigated)** | **Zeeland 2023 (non-irrigated)** | **Valencia 2021 (irrigated)** | **Valencia 2021 (non-irrigated)** | **Valencia 2022 (irrigated)** | **Vojvodina 2023 (irrigated)** | **Fuchsenbigl 2022 (irrigated)** | **Fuchsenbigl 2022 (non-irrigated)** | **Großnondorf 2022** | **Fuchsenbigl 2023 (irrigated)** | **Fuchsenbigl 2023 (non-irrigated)** | **Großnondorf 2023** |
|---|---|---|---|---|---|---|---|---|---|---|---|---|---|---|---|---|
| Acoustic | yes* | yes | yes | yes | yes | yes | yes* | yes* | yes* | yes* | yes | yes | yes | yes | yes | yes |
| Agata | yes* | yes | yes | yes | yes | yes | yes* | yes* | yes* | yes* | yes | yes | yes | yes | yes | yes |
| Agria | yes* | yes | yes | yes | yes | yes | yes* | yes* | yes* | yes* | yes | yes | yes | yes | yes | yes |
| Allison | yes | yes | yes | yes | yes | yes | yes | yes | yes | yes | no | no | no | no | no | no |
| Alonso | yes | yes | yes | yes | yes | yes | yes | yes | yes | yes | no | no | no | no | no | no |
| Altus | yes | yes | yes | yes | yes | yes | yes* | yes* | yes* | yes* | no | no | no | no | no | no |
| Alverstone r. | yes | yes | yes | yes | yes | yes | yes | yes | yes | yes | no | no | no | no | no | no |
| Arizona | yes | yes | yes | yes | yes | yes | yes | yes | yes | yes | no | no | no | no | no | no |
| Asterix | yes | yes | yes | yes | yes | yes | yes | yes | yes | yes | no | no | no | no | no | no |

| | | | | | | | | | | | | | | | | |
|---|---|---|---|---|---|---|---|---|---|---|---|---|---|---|---|---|
| Atlanti c | yes* | yes | yes | yes | yes | yes | yes* | yes* | yes* | yes* | no | no | no | no | no | no |
| Axion | yes | yes | yes | yes | yes | yes | yes | yes | yes | yes | no | no | no | no | no | no |
| Bartin a | yes* | yes | yes | yes | yes | yes | yes* | yes* | yes* | yes* | no | no | no | no | no | no |
| Bricat a | yes | yes | yes | yes | yes | yes | yes | yes | yes | yes | no | no | no | no | no | no |
| Camel | yes | yes | yes | yes | yes | yes | yes | yes | yes | yes | no | no | no | no | no | no |
| Challe nger | yes | yes | yes | yes | yes | yes | yes | yes | yes | yes | no | no | no | no | no | no |
| Charlo tte | yes | yes | yes | yes | yes | yes | yes | yes | yes | yes | no | no | no | no | no | no |
| Colom ba | yes* | yes | yes | yes | yes | yes | yes* | yes* | yes* | yes* | yes | yes | yes | yes | yes | yes |
| Delia red | yes | yes | yes | yes | yes | yes | yes | yes | yes | yes | no | no | no | no | no | no |
| Desire e | yes* | yes | yes | yes | yes | yes | yes* | yes* | yes* | yes* | yes | yes | yes | yes | yes | yes |
| El mundo | yes | yes | yes | yes | yes | yes | yes | yes | yes | yes | no | no | no | no | no | no |
| Erika | yes | yes | yes | yes | yes | yes | yes | yes | yes | yes | yes | yes | yes | yes | yes | yes |

| | | | | | | | | | | | | | | | | |
|---|---|---|---|---|---|---|---|---|---|---|---|---|---|---|---|---|
| Eurostar | yes | yes | yes | yes | yes | yes | yes | yes | yes | yes | no | no | no | no | no | no |
| Evora | yes | yes | yes | yes | yes | yes | yes | yes | yes | yes | no | no | no | no | no | no |
| Farida | yes* | yes | yes | yes | yes | yes | yes* | yes* | yes* | yes* | no | no | no | no | no | no |
| Flamenco | yes | yes | yes | yes | yes | yes | yes | yes | yes | yes | no | no | no | no | no | no |
| Fontane | yes | yes | yes | yes | yes | yes | yes | yes | yes | yes | no | no | no | no | no | no |
| Hermosa | yes | yes | yes | yes | yes | yes | yes | yes | yes | yes | no | no | no | no | no | no |
| Innovator | yes | yes | yes | yes | yes | yes | yes | yes | yes | yes | no | no | no | no | no | no |
| Kuroda | yes | yes | yes | yes | yes | yes | yes | yes | yes | yes | no | no | no | no | no | no |
| Lady claire | yes | yes | yes | yes | yes | yes | yes | yes | yes | yes | no | no | no | no | no | no |
| Lady rosetta | yes* | yes | yes | yes | yes | yes | yes* | yes* | yes* | yes* | yes | yes | yes | yes | yes | yes |
| Meireska | yes | yes | yes | yes | yes | yes | yes | yes | yes | yes | yes | yes | yes | yes | yes | yes |
| Memphis | yes | yes | yes | yes | yes | yes | yes | yes | yes | yes | no | no | no | no | no | no |

| | | | | | | | | | | | | | | | | |
|---|---|---|---|---|---|---|---|---|---|---|---|---|---|---|---|---|
| Mondial | yes | yes | yes | yes | yes | yes | yes | yes | yes | yes | no | no | no | no | no | no |
| Musica | yes* | yes | yes | yes | yes | yes | yes* | yes* | yes* | yes* | yes | yes | yes | yes | yes | yes |
| Nicola | yes* | yes | yes | yes | yes | yes | yes* | yes* | yes* | yes* | no | no | no | no | no | no |
| Orchestra | yes | yes | yes | yes | yes | yes | no | no | yes | yes | no | no | no | no | no | no |
| Rosi | yes* | yes | yes | yes | yes | yes | yes* | yes* | yes* | yes* | yes | yes | yes | yes | yes | yes |
| Russet burbank | yes* | yes | yes | yes | yes | yes | yes* | yes* | yes* | yes* | no | no | no | no | no | no |
| Sagitta | yes | yes | yes | yes | yes | yes | yes | yes | yes | yes | no | no | no | no | no | no |
| Sound | yes* | yes | yes | yes | yes | yes | yes* | yes* | yes* | yes* | no | no | no | yes | yes | yes |
| Spunta | yes* | yes | yes | yes | yes | yes | yes* | yes* | yes* | yes* | yes | yes | yes | yes | yes | yes |
| Taurus | yes | yes | yes | yes | yes | yes | yes | yes | yes | yes | no | no | no | no | no | no |
| Valdivia | yes | yes | yes | yes | yes | yes | yes | yes | yes | yes | yes | yes | yes | yes | yes | yes |

Supplementary table S1.3: Info about varieties included in field trials. Market: 1 - table variety, 2 - french fry variety, 3 - crisp variety, 7 - starch variety. Maturity scores reflect earliness of varieties: 30 - full on flowering healthy green crop at the time of scoring (= very late variety), 40 - healthy green crop (= medium late), 80 - crop almost completely dead stems not completely dry (= early), 90 - crop completely dead (= very early). The maturity scores marked with asterisk (*) were imputed.

| Variety | Breeder | Mother | Father | Breeding target | Market | Maturity score |
|---|---|---|---|---|---|---|
| Acoustic | Meijer Seedpotatoes Heselmans G. | ORCHESTRA | DOB1997-507-01 | 101 - CONSUMPTIE/LUX | 1 | 60 |
| Agata | AGRICO Research | BOHM 52- 72 | SIRCO | 17 - EIGEN INVOER | 1 | 82 |
| Agria | BOHM LUNEBURG | QUARTA | SEMLO | 864 - MP home fry | 2 | 56 |
| Allison | HZPC Research BV | HEO 98-1620 | AGATA | 862 - Early lux | 1 | 56 |
| Alonso | SAATBAUGENOS.WENEN | | | 842 - retail | 1 | 54* |
| Altus | Averis Seeds | KARNA 87-2306 | KARTEL | 432 - ZETMEEL | 7 | 43 |
| Alverstone r. | HZPC Research BV | CRE 98- 200 | INNOVATOR | 418 - FRITES-eigen invoer | 2 | 60 |
| Arizona | AGRICO Research | UK 150-19D22 | MASCOTTE | 946 - TRAD MANUAL Y | 1 | 68 |
| Asterix | HZPC Research BV | CARDINAL | VE 70- 9 | 418 - FRITES-eigen invoer | 1 | 54 |
| Atlantic | MAINE AGR. EXP. STA. | WAUSEON | LENAPE | 722 - crisp | 3 | 64 |
| Axion | AVERIS Kamping T. | KARNA 87-2306 | STABILO | 432 - ZETMEEL | 7 | 44 |

| | | | | | | |
|---|---|---|---|---|---|---|
| Bartina | van der Werff sr. Y.P. | | ZPC 62- 75 | 945 - TRAD MANUAL R | 1 | 54 |
| Bricata | KWS POTATO B.V. KWS | OPAL | VR 808 | 18 - FRITES | 2 | 64 |
| Camel | KWS POTATO B.V. KWS | ROSARA | RZ- 97-6139 | 861 - Maincrop lux | 1 | 67 |
| Challen ger | HZPC Research BV | AZIZA | VICTORIA | 418 - FRITES-eigen invoer | 2 | 56 |
| Charlott e | ST.THEGONNEC/CLAUSE | HANSA | DANAE | 862 - Early lux | 1 | 77 |
| Colomb a | HZPC Research BV | CARRERA | AGATA | 861 - Maincrop lux | 1 | 81 |
| Delia red | HZPC Research BV | HIL 95- 303 | DYNAMICA | 418 - FRITES-eigen invoer | 2 | 63 |
| Desiree | HZPC Research BV | URGENTA | DEPESCHE | 870 - Maincrop red skin | 1 | 57 |
| El mundo | KWS POTATO B.V. KWS | HEO 95- 84 | VALOR | 164 - ONBEKEND | 1 | 59 |
| Erika | SAATBAUGENOS.WENE N | MARABEL | AR 88- 156 | 432 - ZETMEEL | 1 | 73 |
| Eurosta r | Van Rijn Van Rijn | VICTORIA | INNOVATOR | 164 - ONBEKEND | 2 | 60 |
| Evora | HZPC Research BV | LEE 92- 196 | VALOR | 861 - Maincrop lux | 1 | 67 |
| Farida | van der Werff sr. Y.P. | RZ- 91-2313 | VDW 87- 36 | 946 - TRAD MANUAL Y | 1 | 63 |

| | | | | | | |
|---|---|---|---|---|---|---|
| Flamenco | HZPC Research BV | RED SCARLETT | RED CLOUD | 946 - TRAD MANUAL Y | 1 | 63 |
| Fontane | AGRICO Research | AGRIA | AR 76 34 3 | 418 - FRITES-eigen invoer | 2 | 59 |
| Hermosa | Kingma H.J. | CARRERA | MONDIAL | 946 - TRAD MANUAL Y | 1 | 72 |
| Innovator | HZPC Research BV | SHEPODY | RZ- 84-2580 | 418 - FRITES-eigen invoer | 2 | 68 |
| Kuroda | KUIK G. | AR 76 199 3 | KONST 80-1407 | 861 - Maincrop lux | 1 | 54 |
| Lady claire | Meijer Seedpotatoes Heselmans G. | AGRIA | KW78- 34- 470 | 722 - crisp | 3 | 73 |
| Lady rosetta | Meijer Seedpotatoes Heselmans G. | CARDINAL | VTN2 62- 33- 3 | 722 - crisp | 3 | 63 |
| Meireska | SAATBAUGENOS.WENEN | | | 164 - ONBEKEND | 1 | 61* |
| Memphis | Mulder H. | MUH 92- 13 | MUH 91- 13 | 946 - TRAD MANUAL Y | 1 | 66 |
| Mondial | Kweekbedrijf D. Biemond BV | SPUNTA | VE 66-295 | 946 - TRAD MANUAL Y | 1 | 50 |
| Musica | Meijer Seedpotatoes Heselmans G. | CMK 93-042-005 | LADY CHRISTL | 861 - Maincrop lux | 1 | 63 |

| | | | | | | |
|---|---|---|---|---|---|---|
| Nicola | SAATZUCHT SOLT-BERG. | CLIVIA | 6430 1011 | 861 - Maincrop lux | 1 | 61 |
| Orchestra | Meijer Seedpotatoes Heselmans G. | MARADONNA | CUPIDO | 862 - Early lux | 1 | 70 |
| Rosi | HZPC Research BV | SYMFONIA | MOZART | 861 - Maincrop lux | 1 | 56 |
| Russet burbank | onbekend | EARLY ROSE | | 418 - FRITES-eigen invoer | 2 | 56 |
| Sagitta | HZPC Research BV | GALLIA | RZ- 86-2918 | 418 - FRITES-eigen invoer | 2 | 63 |
| Sound | Meijer Seedpotatoes Heselmans G. | | | 842 - retail | 1 | 59* |
| Spunta | HZPC Research BV | BEA | USDA 96 56 | 946 - TRAD MANUAL Y | 1 | 64 |
| Taurus | HZPC Research BV | PANDA | RZ- 87- 44 | 722 - crisp | 3 | 63 |
| Valdivia | SAATBAUGENOS.WENEN | VIENNA | NO 00-205 | 842 - retail | 1 | 58 |

Supplementary table S1.4: Drone fly-by dates for each field trial

| **Filed trial** | **Drone fly by date** | **Corresponding days post planting** | **Assigned days post planting** |
|---|---|---|---|
| Fuchsenbigl 2022 (irrigated) | 15.06.2022 | 55 | 60 |
| Fuchsenbigl 2022 (non-irrigated) | 15.06.2022 | 55 | 60 |
| Fuchsenbigl 2023 (irrigated) | 27.06.2023 | 62 | 60 |
| Fuchsenbigl 2023 (non-irrigated) | 27.06.2023 | 62 | 60 |
| Großnondorf 2023 (non-irrigated) | 27.06.2023 | 62 | 60 |
| Großnondorf 2022 (non-irrigated) | 15.06.2022 | 62 | 60 |
| Valencia 2021 (irrigated) | 15.03.2021 | 40 | 40 |
| Valencia 2021 (irrigated) | 31.03.2021 | 56 | 60 |
| Valencia 2021 (irrigated) | 27.04.2021 | 83 | 90 |
| Valencia 2021 (irrigated) | 21.05.2021 | 107 | 120 |
| Valencia 2021 (non-irrigated) | 21.05.2021 | 37 | 40 |
| Valencia 2021 (non-irrigated) | 11.06.2021 | 58 | 60 |
| Valencia 2021 (non-irrigated) | 15.07.2021 | 92 | 90 |

| | | | |
|---|---|---|---|
| Valencia 2021 (non-irrigated) | Imputed by copying values from Valencia 2021 (non-irrigated) on 15.07.2021 | | 120 |
| Valencia 2022 (irrigated) | Imputed by copying values from Valencia 2021 (irrigated) on 15.03.2021 | | 40 |
| Valencia 2022 (irrigated) | 28.03.2022 | 59 | 60 |
| Valencia 2022 (irrigated) | 2.05.2022 | 94 | 90 |
| Valencia 2022 (irrigated) | 1.06.2022 | 124 | 120 |
| Vojvodina 2023 (irrigated) | 5.06.2023 | 38 | 40 |
| Vojvodina 2023 (irrigated) | 26.06.2023 | 59 | 60 |
| Vojvodina 2023 (irrigated) | 26.07.2023 | 89 | 90 |
| Vojvodina 2023 (irrigated) | 25.08.2023 | 119 | 120 |
| Zeeland 2021 (irrigated) | 14.06.2021 | 42 | 40 |
| Zeeland 2021 (irrigated) | 29.06.2021 | 57 | 60 |
| Zeeland 2021 (irrigated) | 3.08.2021 | 92 | 90 |
| Zeeland 2021 (irrigated) | 19.08.2021 | 108 | 120 |
| Zeeland 2021 (non-irrigated) | 14.06.2021 | 42 | 40 |
| Zeeland 2021 (non-irrigated) | 29.06.2021 | 57 | 60 |

| | | | |
|---|---|---|---|
| Zeeland 2021 (non-irrigated) | 3.08.2021 | 92 | 90 |
| Zeeland 2021 (non-irrigated) | 19.08.2021 | 108 | 120 |
| Zeeland 2022 (irrigated) | 8.06.2022 | 37 | 40 |
| Zeeland 2022 (irrigated) | 29.06.2022 | 58 | 60 |
| Zeeland 2022 (irrigated) | 27.07.2022 | 86 | 90 |
| Zeeland 2022 (irrigated) | 17.08.2022 | 107 | 120 |
| Zeeland 2022 (non-irrigated) | 8.06.2022 | 37 | 40 |
| Zeeland 2022 (non-irrigated) | 29.06.2022 | 58 | 60 |
| Zeeland 2022 (non-irrigated) | 27.07.2022 | 86 | 90 |
| Zeeland 2022 (non-irrigated) | 17.08.2022 | 107 | 120 |
| Zeeland 2023 (irrigated) | 3.07.2023 | 42 | 40 |
| Zeeland 2023 (irrigated) | 18.07.2023 | 57 | 60 |
| Zeeland 2023 (irrigated) | 14.08.2023 | 84 | 90 |
| Zeeland 2023 (irrigated) | 7.09.2023 | 108 | 120 |
| Zeeland 2023 (non-irrigated) | 3.07.2023 | 42 | 40 |
| Zeeland 2023 (non-irrigated) | 18.07.2023 | 57 | 60 |
| Zeeland 2023 (non-irrigated) | 14.08.2023 | 84 | 90 |

| | | | |
|---|---|---|---|
| Zeeland 2023 (non-irrigated) | 7.09.2023 | 108 | 120 |

Supplementary table S1.5: Environmental features.

| Feature | Period | Feature description | Category |
|---|---|---|---|
| 2d_humidity_below_55_0-40 | 0-40 | Number of 2-days periods with humidity below 55 % | Humidity |
| 2d_humidity_below_55_60-90 | 60-90 | | Humidity |
| 2d_tempmax_above_30_60-90 | 60-90 | Number of 2-days periods with maximum daily air temperature above 30°C | Air temperature |
| 2d_tempmax_above_30_90-120 | 90-120 | | Air temperature |
| 3d_humidity_below_55_0-40 | 0-40 | Number of 3-days periods with humidity below 55 % | Humidity |
| 3d_tempmax_above_30_90-120 | 90-120 | Number of 3-days periods with maximum daily air temperature above 30°C | Air temperature |
| 4d_tempmax_above_30_90-120 | 90-120 | Number of 4-days periods with maximum daily air temperature above 30°C | Air temperature |
| avg_soil_moisture_day_max_60-90 | 60-90 | Maximum of the daily mean daytime soil moisture | Soil moisture |
| avg_soil_moisture_day_max_90-120 | 90-120 | | Soil moisture |
| avg_soil_moisture_day_median_60-90 | 60-90 | Median of the daily mean daytime soil moisture | Soil moisture |
| avg_soil_moisture_day_median_90-120 | 90-120 | | Soil moisture |
| avg_soil_moisture_day_min_60-90 | 60-90 | Minimum of the daily mean daytime soil moisture | Soil moisture |
| avg_soil_moisture_day_min_90-120 | 90-120 | | Soil moisture |
| avg_soil_moisture_max_60-90 | 60-90 | Maximum of the daily mean soil moisture | Soil moisture |
| avg_soil_moisture_max_90-120 | 90-120 | | Soil moisture |
| avg_soil_moisture_median_60-90 | 60-90 | Median of the daily mean soil moisture | Soil moisture |
| avg_soil_moisture_median_90-120 | 90-120 | | Soil moisture |
| avg_soil_moisture_min_60-90 | 60-90 | Minimum of the daily mean soil moisture | Soil moisture |
| avg_soil_moisture_min_90-120 | 90-120 | | Soil moisture |
| avg_soil_moisture_night_max_60-90 | 60-90 | Maximum of the daily mean nighttime soil moisture | Soil moisture |
| avg_soil_moisture_night_max_90-120 | 90-120 | | Soil moisture |
| avg_soil_moisture_night_median_60-90 | 60-90 | Median of the daily mean nighttime soil moisture | Soil moisture |
| avg_soil_moisture_night_median_90-120 | 90-120 | | Soil moisture |
| avg_soil_moisture_night_min_60-90 | 60-90 | Minimum of the daily mean nighttime soil moisture | Soil moisture |
| avg_soil_moisture_night_min_90-120 | 90-120 | | Soil moisture |
| avg_soil_temp_day_median_60-90 | 60-90 | Median of the daily mean daytime soil temperature | Soil temperature |
| avg_soil_temp_day_median_90-120 | 90-120 | | Soil temperature |
| avg_soil_temp_day_sum_60-90 | 60-90 | Cumulative sum of the daily mean daytime soil temperatures | Soil temperature |
| avg_soil_temp_day_sum_90-120 | 90-120 | | Soil temperature |
| avg_soil_temp_median_60-90 | 60-90 | Median of the daily mean soil temperature | Soil temperature |

| | | | |
|---|---|---|---|
| avg_soil_temp_median_90-120 | 90-120 | | Soil temperature |
| avg_soil_temp_night_median_60-90 | 60-90 | Median of the daily mean nighttime soil temperature | Soil temperature |
| avg_soil_temp_night_median_90-120 | 90-120 | | Soil temperature |
| avg_soil_temp_night_sum_60-90 | 60-90 | Cumulative sum of the daily mean nighttime soil temperature | Soil temperature |
| avg_soil_temp_night_sum_90-120 | 90-120 | | Soil temperature |
| avg_soil_temp_sum_60-90 | 60-90 | Cumulative sum of the daily mean soil temperature | Soil temperature |
| avg_soil_temp_sum_90-120 | 90-120 | | Soil temperature |
| avg_temp_day_median_0-40 | 0-40 | Median of the daily mean daytime air temperature | Air temperature |
| avg_temp_day_median_40-60 | 40-60 | | Air temperature |
| avg_temp_day_median_60-90 | 60-90 | | Air temperature |
| avg_temp_day_median_90-120 | 90-120 | | Air temperature |
| avg_temp_day_sum_0-40 | 0-40 | Cumulative sum of the daily mean daytime air temperature | Air temperature |
| avg_temp_day_sum_40-60 | 40-60 | | Air temperature |
| avg_temp_day_sum_60-90 | 60-90 | | Air temperature |
| avg_temp_day_sum_90-120 | 90-120 | | Air temperature |
| avg_temp_night_median_0-40 | 0-40 | Median of the daily mean nighttime air temperature | Air temperature |
| avg_temp_night_median_40-60 | 40-60 | | Air temperature |
| avg_temp_night_median_60-90 | 60-90 | | Air temperature |
| avg_temp_night_median_90-120 | 90-120 | | Air temperature |
| avg_temp_night_sum_0-40 | 0-40 | Cumulative sum of the daily mean nighttime air temperature | Air temperature |
| avg_temp_night_sum_40-60 | 40-60 | | Air temperature |
| avg_temp_night_sum_60-90 | 60-90 | | Air temperature |
| avg_temp_night_sum_90-120 | 90-120 | | Air temperature |
| cloudcover_median_0-40 | 0-40 | Median of cloudcover | Cloud cover |
| cloudcover_median_40-60 | 40-60 | | Cloud cover |
| cloudcover_median_60-90 | 60-90 | | Cloud cover |
| cloudcover_median_90-120 | 90-120 | | Cloud cover |
| daylength_median_0-40 | 0-40 | Median of daylenght | Daylength |
| daylength_median_40-60 | 40-60 | | Daylength |
| daylength_median_60-90 | 60-90 | | Daylength |
| daylength_median_90-120 | 90-120 | | Daylength |
| daylength_sum_0-40 | 0-40 | Cumulative sum of daylenght | Daylength |
| daylength_sum_40-60 | 40-60 | | Daylength |
| daylength_sum_60-90 | 60-90 | | Daylength |
| daylength_sum_90-120 | 90-120 | | Daylength |
| GDD_sum_0-40 | 0-40 | Cumulative sum of growing degree days | Air temperature |
| GDD_sum_40-60 | 40-60 | | Air temperature |

| | | | |
|---|---|---|---|
| GDD_sum_60-90 | 60-90 | | Air temperature |
| GDD_sum_90-120 | 90-120 | | Air temperature |
| humidity_max_0-40 | 0-40 | | Humidity |
| humidity_max_40-60 | 40-60 | | Humidity |
| humidity_max_60-90 | 60-90 | | Humidity |
| humidity_max_90-120 | 90-120 | Maximum of daily air humidity | Humidity |
| humidity_median_0-40 | 0-40 | | Humidity |
| humidity_median_40-60 | 40-60 | | Humidity |
| humidity_median_60-90 | 60-90 | | Humidity |
| humidity_median_90-120 | 90-120 | Median of daily air humidity | Humidity |
| humidity_min_0-40 | 0-40 | | Humidity |
| humidity_min_40-60 | 40-60 | | Humidity |
| humidity_min_60-90 | 60-90 | | Humidity |
| humidity_min_90-120 | 90-120 | Minimum of daily air humidity | Humidity |
| min_soil_moisture_max_60-90 | 60-90 | Maximum of the daily minimum soil moisture | Soil moisture |
| min_soil_moisture_max_90-120 | 90-120 | | Soil moisture |
| min_soil_moisture_median_60-90 | 60-90 | Median of the daily minimum soil moisture | Soil moisture |
| min_soil_moisture_median_90-120 | 90-120 | | Soil moisture |
| min_soil_moisture_min_60-90 | 60-90 | Minimum of the daily minimum soil moisture | Soil moisture |
| min_soil_moisture_min_90-120 | 90-120 | | Soil moisture |
| min_soil_moisture_night_max_60-90 | 60-90 | Maximum of the daily minimum nighttime soil moisture | Soil moisture |
| min_soil_moisture_night_max_90-120 | 90-120 | | Soil moisture |
| min_soil_moisture_night_median_60-90 | 60-90 | Median of the daily minimum nighttime soil moisture | Soil moisture |
| min_soil_moisture_night_median_90-120 | 90-120 | | Soil moisture |
| min_soil_moisture_night_min_60-90 | 60-90 | Minimum of the daily minimum nighttime soil moisture | Soil moisture |
| min_soil_moisture_night_min_90-120 | 90-120 | | Soil moisture |
| min_soil_temp_night_median_60-90 | 60-90 | Median of the daily minimum nighttime soil temperature | Soil temperature |
| min_soil_temp_night_median_90-120 | 90-120 | | Soil temperature |
| min_soil_temp_night_sum_60-90 | 60-90 | Cumulative sum of the daily minimum nighttime soil temperature | Soil temperature |
| min_soil_temp_night_sum_90-120 | 90-120 | | Soil temperature |
| min_temp_night_median_0-40 | 0-40 | | Air temperature |
| min_temp_night_median_40-60 | 40-60 | | Air temperature |
| min_temp_night_median_60-90 | 60-90 | | Air temperature |
| min_temp_night_median_90-120 | 90-120 | Median of the daily minimum nighttime air temperature | Air temperature |

| | | | |
|---|---|---|---|
| min_temp_night_sum_0-40 | 0-40 | Cumulative sum of the daily minimum nighttime air temperature | Air temperature |
| min_temp_night_sum_40-60 | 40-60 | | Air temperature |
| min_temp_night_sum_60-90 | 60-90 | | Air temperature |
| min_temp_night_sum_90-120 | 90-120 | | Air temperature |
| num_days_avg_soil_moisture_below_20_90-120 | 90-120 | Number of days with daily mean soil moisture below 20 % | Soil moisture |
| num_days_avg_soil_temp_night_above_20_60-90 | 60-90 | Number of days with daily mean nighttime soil moisture above 20 % | Soil temperature |
| num_days_avg_soil_temp_night_above_20_90-120 | 90-120 | | Soil temperature |
| num_days_avg_soil_temp_night_above_22_90-120 | 90-120 | | Soil temperature |
| num_days_avg_temp_day_above_25_60-90 | 60-90 | Number of days with mean daily daytime air temperature above 25°C | Air temperature |
| num_days_avg_temp_day_above_25_90-120 | 90-120 | | Air temperature |
| num_days_avg_temp_night_above_20_40-60 | 40-60 | Number of days with mean daily nighttime air temperature above 20°C | Air temperature |
| num_days_avg_temp_night_above_20_60-90 | 60-90 | | Air temperature |
| num_days_avg_temp_night_above_20_90-120 | 90-120 | | Air temperature |
| num_days_avg_temp_night_above_22_40-60 | 40-60 | Number of days with mean daily nighttime air temperature above 22°C | Air temperature |
| num_days_avg_temp_night_above_22_60-90 | 60-90 | | Air temperature |
| num_days_avg_temp_night_above_22_90-120 | 90-120 | | Air temperature |
| num_days_humidity_below_60_0-40 | 0-40 | Number of days with air humidity below 60 % | Humidity |
| num_days_humidity_below_60_40-60 | 40-60 | | Humidity |
| num_days_humidity_below_60_60-90 | 60-90 | | Humidity |
| num_days_humidity_below_60_90-120 | 90-120 | | Humidity |
| num_days_min_soil_temp_above_20_60-90 | 60-90 | Number of days with minimum daily soil temperature above 20°C | Soil temperature |
| num_days_min_soil_temp_above_20_90-120 | 90-120 | | Soil temperature |
| num_days_min_soil_temp_night_above_18_60-90 | 60-90 | Number of days with minimum daily nighttime soil temperature above 18°C | Soil temperature |
| num_days_min_soil_temp_night_above_18_90-120 | 90-120 | | Soil temperature |

| | | | |
|---|---|---|---|
| num_days_min_soil_temp_night_above_20_60-90 | 60-90 | Number of days with minimum daily nighttime soil temperature above 20°C | Soil temperature |
| num_days_min_soil_temp_night_above_20_90-120 | 90-120 | | Soil temperature |
| num_days_min_temp_night_above_16_40-60 | 40-60 | Number of days with minimum daily nighttime air temperature above 16°C | Air temperature |
| num_days_min_temp_night_above_16_60-90 | 60-90 | | Air temperature |
| num_days_min_temp_night_above_16_90-120 | 90-120 | | Air temperature |
| num_days_min_temp_night_above_18_60-90 | 60-90 | Number of days with minimum daily nighttime air temperature above 18°C | Air temperature |
| num_days_min_temp_night_above_18_90-120 | 90-120 | | Air temperature |
| num_days_tempavg_above_22_40-60 | 40-60 | Number of days with mean daily air temperature above 22°C | Air temperature |
| num_days_tempavg_above_22_60-90 | 60-90 | | Air temperature |
| num_days_tempavg_above_22_90-120 | 90-120 | | Air temperature |
| num_days_tempavg_above_25_60-90 | 60-90 | Number of days with mean daily air temperature above 25°C | Air temperature |
| num_days_tempavg_above_25_90-120 | 90-120 | | Air temperature |
| num_days_tempmax_above_30_60-90 | 60-90 | Number of days with đ above 30°C | Air temperature |
| num_days_tempmax_above_30_90-120 | 90-120 | | Air temperature |
| precip_median_0-40 | 0-40 | Median of precipitation | Precipitation |
| precip_median_40-60 | 40-60 | | Precipitation |
| precip_median_60-90 | 60-90 | | Precipitation |
| precip_median_90-120 | 90-120 | | Precipitation |
| precip_sum_0-40 | 0-40 | Cumulative sum of precipitation | Precipitation |
| precip_sum_40-60 | 40-60 | | Precipitation |
| precip_sum_60-90 | 60-90 | | Precipitation |
| precip_sum_90-120 | 90-120 | | Precipitation |
| rGDD_sum_0-40 | 0-40 | Cumulative sum of revised growing degree days | Air temperature |
| rGDD_sum_40-60 | 40-60 | | Air temperature |
| rGDD_sum_60-90 | 60-90 | | Air temperature |
| rGDD_sum_90-120 | 90-120 | | Air temperature |
| solarenergy_median_0-40 | 0-40 | Median of solar energy | Solar energy |
| solarenergy_median_40-60 | 40-60 | | Solar energy |
| solarenergy_median_60-90 | 60-90 | | Solar energy |
| solarenergy_median_90-120 | 90-120 | | Solar energy |
| solarenergy_sum_0-40 | 0-40 | Cumulative sum of solar energy | Solar energy |
| solarenergy_sum_40-60 | 40-60 | | Solar energy |

| | | | |
|---|---|---|---|
| solarenergy_sum_60-90 | 60-90 | | Solar energy |
| solarenergy_sum_90-120 | 90-120 | | Solar energy |
| solarradiation_median_0-40 | 0-40 | Median of solar radiation | Solar radiation |
| solarradiation_median_40-60 | 40-60 | | Solar radiation |
| solarradiation_median_60-90 | 60-90 | | Solar radiation |
| solarradiation_median_90-120 | 90-120 | | Solar radiation |
| solarradiation_sum_0-40 | 0-40 | Cumulative sum of solar radiation | Solar radiation |
| solarradiation_sum_40-60 | 40-60 | | Solar radiation |
| solarradiation_sum_60-90 | 60-90 | | Solar radiation |
| solarradiation_sum_90-120 | 90-120 | | Solar radiation |
| tempavg_median_0-40 | 0-40 | Median of mean daily air temperature | Air temperature |
| tempavg_median_40-60 | 40-60 | | Air temperature |
| tempavg_median_60-90 | 60-90 | | Air temperature |
| tempavg_median_90-120 | 90-120 | | Air temperature |
| tempavg_sum_0-40 | 0-40 | Cumulative sum of mean daily air temperature | Air temperature |
| tempavg_sum_40-60 | 40-60 | | Air temperature |
| tempavg_sum_60-90 | 60-90 | | Air temperature |
| tempavg_sum_90-120 | 90-120 | | Air temperature |
| tempmax_median_0-40 | 0-40 | Median of maximum daily air temperature | Air temperature |
| tempmax_median_40-60 | 40-60 | | Air temperature |
| tempmax_median_60-90 | 60-90 | | Air temperature |
| tempmax_median_90-120 | 90-120 | | Air temperature |
| tempmax_sum_0-40 | 0-40 | Cumulative sum of maximum daily air temperature | Air temperature |
| tempmax_sum_40-60 | 40-60 | | Air temperature |
| tempmax_sum_60-90 | 60-90 | | Air temperature |
| tempmax_sum_90-120 | 90-120 | | Air temperature |
| tempmin_median_0-40 | 0-40 | Median of minimum daily air temperature | Air temperature |
| tempmin_median_40-60 | 40-60 | | Air temperature |
| tempmin_median_60-90 | 60-90 | | Air temperature |
| tempmin_median_90-120 | 90-120 | | Air temperature |
| tempmin_sum_0-40 | 0-40 | Cumulative sum of minimum daily air temperature | Air temperature |
| tempmin_sum_40-60 | 40-60 | | Air temperature |
| tempmin_sum_60-90 | 60-90 | | Air temperature |
| tempmin_sum_90-120 | 90-120 | | Air temperature |

Supplementary table S1.6: Genes for gene expression analysis. Genes with short names in bold were selected based on the reanalysis of available gene expression data. Target genes were systematically filtered from the machine learning feature set based on the following criteria: genes marked with asterisk (*) were removed due to low expression values across all measured samples, while genes marked with hash ([#]) were removed due to high pairwise Spearman correlation ($\rho > 0.85$) with other retained genes.

| Gene name | Gene description | Gene involved in process | DMv6.1 geneID | Target sequence for probe design | Class |
|---|---|---|---|---|---|
| 13-LOX | 13-lipoxygenase | JA signalling | Soltu.DM.03G037120 | TCGTTCTGGTCGTGTTCCTACAGACACAGATATAA<br>GTGCGGAGAGTCGTGTGGAGAAGCCAAATCCAAC<br>GTATGTTCCGAGAGATGAACAATTTGAGGAG | Target |
| ACO2 | 1-aminocyclopropane-1-carboxylate oxidase 2 | Ethylene signalling | Soltu.DM.07G016780 | ATGGACACTGTGGAGAAATTGACAAAGGGACATTA<br>CAAGAAGTGCATGGAACAGAGATTTAAGGAACTGG<br>TGGCAAGTAAGGGACTTGAAGCTGTTCAAG | Target |
| ACX3 | Medium-chain acyl-CoA oxidase 3 | JA signalling | Soltu.DM.10G003860 | TCAAGTGCATTCAAAGCTACTACAAGTTGGCATAAC<br>ATGCGAACTCTTCAGGAATGTCGTGAAGCCTGTGG<br>AGGCCAAGGGTTGAAGACCGAAAATAGGA | Target |
| ADH1 | Alcohol dehydrogenase 1 | Waterlogging stress | Soltu.DM.06G016070 | CCTTGCTCCTCTTGACAAAGTATGTGTCCTTAGTTG<br>TGGAATCTCGACAGGCCTTGGAGCAACTTTGAATG<br>TTGCTAAACCAACAAAAGGCTCAAGTGTG | Target |
| BEL5[#] | BEL1-like homeodomain transcription factor 5 | Tuberisation | Soltu.DM.04G035890 | TGTCGGTGTGCCTCTTCCGGCAGTAAGTTTGCAC<br>GATCAGATCAATCATCATGGACTTTTACAGCGTATG<br>TGGAACAACCAAGATCAATCTCAGCAGGTG | Target |
| CAT1 | Catalase 1 | ROS signalling | Soltu.DM.12G004810 | ATGAACGTGGAAGCCCCGAGTCAATCAGGGACAT<br>TCGCGGTTTTGCTGTCAAGTTTTACACCAGAGAGG<br>GTAACTTTGATCTTGTTGGAAACAATGTCCC | Target |
| CDF1 | DOF zink finger transcription factor 1 | Tuberisation | Soltu.DM.05G005140 | CAGCCATTTCCTCCTCAAGTACCGTACTTTCAGGG<br>CGCTCCGTGGCCTTATTCTGGCTTTCCAGTATCATT<br>CTATCCAGCAACACCGTACTGGGGCTGCA | Target |
| CDF2[#] | DOF zink finger transcription factor 2 | Tuberisation | Soltu.DM.02G009620 | AAAGAGTTCTATATGGGCGACATTGGGAATAAAACA<br>TGATACCGTTGATTCAGTTGGTGGAAGTCCTTTCA<br>GTGCTTTTCAGCCGAAGAATGATGACAAC | Target |
| CDF3* | DOF zink finger transcription factor 3 | Tuberisation | Soltu.DM.02G031630 | CTGTAGAAATGCAGTACCTCCACCGGGTTACTCCC<br>TTCCTGGCATTCCTATGCCGTTCTTCCCTGCAACC<br>ACTTATTGGGGTTGTACAATACCAGGTTCT | Target |

| | | | | | |
|---|---|---|---|---|---|
| **CML45** | Calcium-binding EF-hand family protein 45 | Ca-regulated | Soltu.DM.02G031620 | GTTGATGGAAGCTTAAACAGGGAAGAGGTGGAAATGGTTATGGCTAATTTAGGTATTTTTGCCAATCCTCAAGGTGTGAAGATTCAGGGGAGGTTGGATT | Target |
| CO[#] | Constants | Circadian clock, Tuberisation | Soltu.DM.02G030260 | TAAGAACAACATTAACAATAACAACAACAATCAAAATAACAACTATGGGATGTTGTTTGGTGGGGAAGTAGTGGATGAATACTTGGATCTTGCAGAGTAT | Target |
| **CPK17** | Calcium-dependent protein kinase 17 | Ca-regulated | Soltu.DM.12G001160 | ATGCATCACCAAACCATTCTTCAAAGCCATCTAAGGCAGCCCCAATAGGGCCAGTATTAGGCAGGCCAATGGAGGATATAAAGGCAACATACACCCTAGG | Target |
| **CRINKLY4-like** | Receptor-like serine/threonine-protein kinase | Drought stress | Soltu.DM.01G026590 | GGAGGTGGCTGAATGTTTGAGTGGGTTGAGTAAGTTGGTACCTCTACATTCCTGGAATGGTTTTACCAATCCTTGTTTGATGGTTGAGACCGTGGGCCGA | Target |
| **CYP704A2** | Cytochrome P450, family 704, subfamily A2 | Heat stress | Soltu.DM.01G033030 | TTAGTTTTAATACTAAGGATCTATGCTGGCAAGTCAATCAGAAACCCCAAATCTGCACCAGTGGTAGGAACTGTGTTTCACCAACTCTTCTACTTCAAAA | Target |
| **DSP4** | Dual-specificity protein-like phosphatase 4 | Heat stress | Soltu.DM.11G004900 | AAGCTATAAGCTTAGTGAAGCTTTCGATTTACTCATGAGCAAACGCTCCTGCTTTCCGAAACTTGATGCCATCAAAAGTGCCACAGCAGACGTTCTTACA | Target |
| ERF-1 | Ethylene responsive transcription factor 1 | Ethylene signalling | Soltu.DM.03G014550 | GCTTTTAGGATGCGCGGTACTAAGGCTCTATTGAATTTCCCGCATAGAATCGGTTTAAATGAGCCGGAGCCGGTTAGAGTGACGGTTAAGAGACGATTAC | Target |
| FKF1[#] | Flavin-binding kelch domain F box protein 1 | Tuberisation | Soltu.DM.01G000490 | AGTGCATGTGCTGCAGGGAACAGGCTAGTGTTATTTGGAGGGGAAGGTGTTAATATGCAGCCAATGGATGACACATTTGTTCTCAATCTTGATGCTGCTA | Target |
| GI1 | Gigantea1 | Circadian clock | Soltu.DM.03G010520 | GCCATCAGGGCCTCTGTGTAAAGACTCCTATGATTGTAAAAGTTCACTTGTGTGTGAGAAAGCTTCAGACTCAAGTTCACACTCTAGTGAGATTGCTGGG | Target |
| GI2 | Gigantea2 | Circadian clock | Soltu.DM.04G027760 | TGATATCCATAGTAAAGTTGTTGCATCAATTGTTGACAAGGCCGAACCATTGGAAGCACACCTAATACCTGTTCCTGTTAAGAAAAGAAGCTCGTGCTTG | Target |
| HB20 | Homeobox leucine zipper transcription factor 20 | ABA signalling | Soltu.DM.01G035490 | TGCAAAAACTGAATGATGTGATGCAGAAAGAAAGGGGGAAGTACTGTTGCAAGCAAGAAATCGAGTTAGACAACAGATGTGCTGCAATTAAGAAAAATGA | Target |
| HSC70 | Heat shock cognate protein 70 | Heat stress | Soltu.DM.04G007430 | CTTGAGTGGTGAAGGGAATGAAAAAGTTCAGGACCTTTTGCTGTTGGATGTTACACCTCTTTCCCTTGGTCTGGAGACTGCTGGAGGTGTTATGACCACC | Target |

| | | | | | |
|---|---|---|---|---|---|
| ICL | Isocitrate lyase | Waterlogging stress | Soltu.DM.07G017900 | AAGCTGTGTAAGCTTTTCGTGGAGCGCGGTGCTGCTGGTGTCCACATTGAGGATCAGTCCTCTGTTACTAAGAAATGTGGTCATATGGCTGGTAAAGTGC | Target |
| LTP3 | Lipid transfer protein 3 | Waterlogging stress | Soltu.DM.08G016090 | GTATCGCCAAAACGACGGCGGACCGCCAAGGCGTTTGCTCGTGCTTGAAAATGGCTGCCTCAAGTGTTAGTGGTATCGATTTTAAGAATGCTGCTGCCCT | Target |
| **NFYA-1** | Nuclear transcription factor Y subunit A-1 | Drought stress | Soltu.DM.10G028300 | GAACTAATGGCACCCTCCTCAATTGCATTTAAATCATACCCCTATTCAGAAGTACCACAATATTCTGGTGGCAATGTTACTATTGCTTGTGGTGAACCTA | Target |
| OXI1 | Serine/threonine-protein kinase oxidative signal-inducible1 | ROS signalling | Soltu.DM.09G016790 | ATTGAGAAACTTGATAAACAAGTTACTCGAAAAGGATCCGAAGCAGAGGATCTCTGTGGATGAAATCAAAGGTCACGATTTCTTCAGATCCGTAGACTGG | Target |
| P5CS1 | Delta1-pyrroline-5-carboxylate synthase 1 | ABA signalling | Soltu.DM.06G007050 | GTATTCCTGAAGCAAATTCCTTCCATCATGAATATGGTGCACTGGCTTGCACCGTGGAAATTGTTGAAGATGTGAATACTGCTATAGAGCACATACATCG | Target |
| PIF4 | Phytochrome interacting transcription factor 4 | Senescence | Soltu.DM.07G014300 | ATCTGGAACTGCCCGAAGGAGCCGTGCTGCAGAAGTGCATAATCTCTCTGAAAGGAGACGGAGGGACAGAATCAATGAGAAAATGAAGGCCTTGCAAGAG | Target |
| PP2C_1 | Protein phosphatase 2C | Waterlogging stress | Soltu.DM.06G031720 | GTGATAACGCTGTGGAAAGTTCTAGCACCGACGATGATGAAGAACAACTGAGGTTAGTAGAGACGAAAGGGTCACCGCCAACTATTATTTTGACTCCATT | Target |
| PP2C_2 | Protein phosphatase 2CA | Waterlogging stress | Soltu.DM.03G012480 | AAAGAAGGTAGCAGAGAATAAAGAAACCGAAACGAACGCTTTGGATTTGACGGAATCTGCTTCTATATCGTTGAATATTGAACGCCAGGGAGTATCCGAC | Target |
| PPDK | Pyruvate orthophosphate dikinase | Waterlogging stress | Soltu.DM.01G023530 | GCATTAATCAGATAACAGGACTTAAAGGAACTGCAGTAAACATTCAATGCATGGTGTTTGGGAACATGGGTAACACTTCAGGAACAGGTGTTCTCTTCAC | Target |
| PR1B | Basic pathogenesis-related protein 1B; CAPE precursor | SA signalling | Soltu.DM.09G007020 | TGATTGTAACTTGATTCATTCCGGTGCAGGGGAGAACCTTGCCAAGGGAACTGGTGATTTCACGGGGAGGGCAGCTGTGCAATTGTGGGTGGGGGAGAAG | Target |
| RBA | Rubisco activase | Photosynthesis | Soltu.DM.10G023180 | GGGGAAAGGGTATGGTAGACCCTCTTTTCCAAGCTCCTGTGGGTACTGGTACTCACCATCCTGTCTTGTCATCATACGAATACATCAGCCAGGGTCTTCG | Target |

| | | | | | |
|---|---|---|---|---|---|
| RBOHA | Respiratory burst NADPH oxidase A | ROS signalling | Soltu.DM.08G028440 | GTGGAGGGTGTGACAGGAATAATAATGGTAATCCTCATGGCCATTGCTTTCACTCTTGCAACGCGATGGTTTAGGCGGAGCCTCATTAAGTTTCCCAAAC | Target |
| RD29B | ow-temperature-induced 65 kDa protein; CAP160 protein | ABA signalling | Soltu.DM.03G017570 | CAGCTCAGAATGCTCTCAAAGGCTTCAAATTTATCAGTAAGACTGATGGTGGCTCTGGATGGGACACCGTCCAACAGCGCTTTGACGAGCTTACTGCCAA | Target |
| RboHD | Respiratory burst NADPH oxidase D | ROS signalling | Soltu.DM.06G024560 | TGCTGCTAAGAACATTGTCGCCTCCAAACTCGGGTATGGTGGAACAGAGGAGGAAACTCAGGCAACAGCAGGCGATAAAGATGCAACAAAAACAACCTCA | Target |
| **SIS** | Conserved gene of unknown function | Ca-regulated | Soltu.DM.09G001460 | ATCAGAACGAAAAAGTACAGCCATGCCATCTTAGCTCATCGATATACTACGGGGGCCAAGATGTCTATTCATTTCCCCAAAATAATAATCAGAGCTCTAC | Target |
| SNRK2 | Sucrose nonfermenting 1(SNF1)-related protein kinase 2 | ABA signalling | Soltu.DM.02G029320 | TCCGGCACAGAGGATCACAATGCCTGAAATCAGAAATCATGTATGGTTTTTAAAGAATCTTCCAGCGGACTTAATAGACGATAAGATGATTAGTGACCAG | Target |
| SP5G | Self-pruning 5G; Flowering locus T / florigen (FT) regulatory protein | Tuberisation | Soltu.DM.05G024030 | AGTTGTACACTACGAGAGCCCACGACCCTCGATGGGAATCCATCGTTATATTTTCGTGTTGTTCCAGCAATTGGGCCGCGAGGCCATCAATGCGCCGGAC | Target |
| SP6A | Self-pruning 6A; Flowering locus T / florigen (FT) regulatory protein | Tuberisation | Soltu.DM.05G026370 | ACATTGGCTGGTCACAGATATCCCAGCAACTACAAATACAAGCTTTGGAAATGAAGTCGTATGCTACGAGAATCCAACACCTACGATGGGAATTCATCGA | Target |
| SWEET | Sugar efflux transporter | Tuberisation | Soltu.DM.06G027970 | ATTTACAAGAAGAAATCAACAGAAGGCTATCAATCAATTCCATATGTGGTTGCTCTATGCAGTGCAATGATTTTGATATACTATGTATATCTTAAAGGAC | Target |
| TFL1* | Terminal flower 1 regulatory protein | Tuberisation | Soltu.DM.03G017110 | TAAACCTAGGGTTGAAGTTCATGGTGGTGATCTCAGATCCTTCTTCACACTGGTCATGATAGATCCAGATGTTCCTGGTCCTAGTGATCCATATCTCAGG | Target |
| TFL_SP | Terminal flower 1; Self-pruning regulatory protein; centroradialis | Tuberisation | Soltu.DM.06G029780 | GTATTTTTGTTGTATAAACAAAGAGGAAGGCAAACAGTGAAAGCACCAACAACAAGGGATCAATTCAATACAAGGAGTTTTTCGGCGGAAAATGGATTGG | Target |
| TOC1 | Circadian clock evening oscillator component | Circadian clock | Soltu.DM.06G025760 | CCAACCTCTATCCATGGAAGTGTGCCTCAGCAATTGGGGATTGAAGAAAAGACTTCATTGGGAGGACATGAACACATTGATACTAACATACAAGTCACTG | Target |

| | | | | | |
|---|---|---|---|---|---|
| TPL | Topless; auxin signalling transcriptional co-repressor | Tuberisation | Soltu.DM.03G031570 | TCTATTGATTATCCATCTGGGGAATCTGATCATGCT GCCAAAAGAACTAGGTCACTGGGAATATCTGATGA GGTTAATCTTCCTGTGAATGTGCTACCTA | Target |
| **WRKY50** | WRKY transcription factor | Ca-regulated | Soltu.DM.04G023540 | AGTCATAACATAAAATGTATGAAAGGTGTAAAGAAG GTGGATGCAAAGTCTAGGGTTGCATTTAGATTTAGA TCAGAGTTGGAGGTGTTGGATGATGGAT | Target |
| **XET** | Xyloglucan endotransglycosylase | Ca-regulated | Soltu.DM.11G019660 | ACCGATTGGTCCGGGGCGCCATTCACAGCTTCCT ATACTTCATTCCACATCGATGGCTGTGAGGCTGTC ACCCCACAAGAAGTGCAAGTTTGTAACACTA | Target |
| **Znf** | RING domain zinc finger transcription factor | Drought stress | Soltu.DM.01G023980 | TGGAGGGTGGTGGTAATTTTAGCCTTGTGTCTTTA GTGAAATGCTTCAATGGGTGCAAAGATCAAGCTCA AGTCTCTAGTTTTGGAACAAGGATTTGGAA | Target |
| COX | Cytochrome c oxidase subunit 1 (mtDNA) | Housekeeping | / | CAAGTCTCTTGCTCCTATTAAGCTCAGCCTTAGTAG AAGTGGGTAGCGGCACTGGGTGGACGGTCTATCC GCCCTTAAGTGGTATTACCAGCCATTCTGG | Housekeeping |
| EF1 | GTP binding Elongation factor Tu family protein | Housekeeping | Soltu.DM.11G022490 | AAGGTAGGATACAACCCTGACAAAATCCCCTTTGT CCCCATCTCTGGTTTTGAAGGAGACAACATGATTG AGCGATCAACCAACCTTGACTGGTACAAGG | Housekeeping |
| cyclophilin | Peptidyl-prolyl cis-trans isomerase family protein | Housekeeping | Soltu.DM.01G050750 | CGTGGTGATGGAGCTCTTCGCCGATACCACTCCC AAAACCGCTGAGAACTTCCGAGCTCTCTGTACCG GTGAAAAAGGTGTTGGAAAGATGGGTAAGCCT | Housekeeping |

### Section 2

Supplementary table S2.1: Results of Wilcoxon rank-sum test for mean daily soil temperature and mean daily soil moisture between irrigated and non-irrigated field trial conducted simultaneously at the same location.

| Field trials | p-value | |
|---|---|---|
| | Mean daily soil temperature | Mean daily soil moisture |
| Fuchsenbigl 2022 | $8.9\times10^{-14}$ | $4.2\times10^{-3}$ |
| Fuchsenbigl 2023 | $3.1\times10^{-14}$ | $< 10^{-16}$ |
| Zeeland 2021 | $1.7\times10^{-3}$ | 0.488 |
| Zeeland 2022 | 0.941 | 0.179 |
| Zeeland 2023 | 0.967 | 0.354 |

Supplementary table S2.2: Results of Wilcoxon rank-sum test for tuber yield and quality variables yield per hectare (YHA), number of tubers per plant (TP), overall impression (OI), and underwater weight (UW) between irrigated and non-irrigated field trials conducted simultaneously at the same location.

| Field trials | p-value | | | |
|---|---|---|---|---|
| | YHA | TP | OI | UW |
| Fuchsenbigl 2022 | $< 10^{-16}$ | $5.9\times10^{-14}$ | $1.5\times10^{-8}$ | $5.2\times10^{-4}$ |
| Fuchsenbigl 2023 | $1.9\times10^{-13}$ | $1.2\times10^{-4}$ | 0.726 | $7.5\times10^{-6}$ |
| Zeeland 2021 | 0.199 | 0.145 | 0.64 | 0.492 |
| Zeeland 2022 | $< 10^{-16}$ | $7.8\times10^{-4}$ | 0.06 | 0.09 |
| Zeeland 2023 | $2.7\times10^{-4}$ | 0.016 | 0.829 | 0.739 |

#### Section 3

Supplementary table S3.1: Minimum and maximum ranks, rank range (maximum minus minimum rank), and standard deviation (SD) of ranks assigned to varieties based on the medians and relative standard deviations of tuber yield and quality traits: yield per hectare, number of tubers per plant, overall impression, and underwater weight.

| Variety | Rank min | Rank max | Rank range | Rank SD |
|---|---|---|---|---|
| Orchestra | 2 | 34 | 32 | 10.6 |
| Bricata | 2 | 34 | 32 | 11.1 |
| Musica | 2 | 31 | 29 | 9.3 |
| Sound | 2 | 32 | 30 | 10.7 |
| Alonso | 1 | 38 | 37 | 15.0 |
| Arizona | 1 | 43 | 42 | 17.1 |
| Allison | 5 | 29 | 24 | 8.8 |
| Challenger | 1 | 33 | 32 | 11.7 |
| Acoustic | 2 | 35 | 33 | 11.8 |
| Evora | 3 | 37 | 34 | 10.1 |
| Fontane | 3 | 30 | 27 | 10.1 |
| Taurus | 1 | 42 | 41 | 15.9 |
| Delia Red | 6 | 33 | 27 | 9.8 |
| Meireska | 11 | 31 | 20 | 7.1 |
| Russet Burbank | 1 | 38 | 37 | 13.1 |
| Kuroda | 12 | 35 | 23 | 8.5 |
| Colomba | 2 | 44 | 42 | 16.4 |
| El Mundo | 2 | 40 | 38 | 14.2 |
| Nicola | 1 | 42 | 41 | 13.6 |
| Erika | 3 | 39 | 36 | 11.7 |
| Camel | 4 | 36 | 32 | 9.8 |
| Agata | 2 | 43 | 41 | 17.2 |

| | | | | |
|---|---|---|---|---|
| Lady Rosetta | 3 | 39 | 36 | 12.2 |
| Atlantic | 4 | 43 | 39 | 16.1 |
| Lady Claire | 6 | 41 | 35 | 13.3 |
| Valdivia | 2 | 37 | 35 | 14.1 |
| Axion | 2 | 38 | 36 | 12.0 |
| Eurostar | 13 | 38 | 25 | 8.1 |
| Alverstone R. | 10 | 35 | 25 | 8.4 |
| Memphis | 16 | 32 | 16 | 6.5 |
| Farida | 2 | 43 | 41 | 13.9 |
| Desiree | 8 | 41 | 33 | 11.4 |
| Mondial | 5 | 44 | 39 | 16.6 |
| Spunta | 2 | 39 | 37 | 12.3 |
| Innovator | 7 | 44 | 37 | 14.3 |
| Rosi | 2 | 42 | 40 | 14.7 |
| Asterix | 3 | 44 | 41 | 16.3 |
| Bartina | 3 | 43 | 40 | 16.1 |
| Altus | 1 | 44 | 43 | 14.8 |
| Sagitta | 14 | 43 | 29 | 10.1 |
| Agria | 13 | 40 | 27 | 11.3 |
| Hermosa | 8 | 41 | 33 | 11.5 |
| Charlotte | 17 | 44 | 27 | 12.0 |
| Flamenco | 19 | 43 | 24 | 8.1 |

Supplementary table S3.2: Results of Wilcoxon rank-sum tests comparing the distributions of $log_2$-transformed gene expression values between top- and low-performing varieties across all field trials, along with the differences between their medians.

| Gene | p-value | Median difference |
|---|---|---|
| SP5G | 0.001 | -1.92 |
| Znf | 0.003 | 1.28 |
| LTP3 | 0.019 | -1.31 |
| CPK17 | 0.029 | -0.48 |
| GI1 | 0.036 | -0.18 |
| SIS | 0.095 | 0.99 |
| SP6A | 0.176 | 0.65 |
| CO | 0.198 | 0.93 |
| ACX3 | 0.229 | 0.16 |
| GI2 | 0.246 | -0.54 |
| ACO2 | 0.264 | 0.44 |
| HSC70 | 0.264 | 0.19 |
| 13-LOX | 0.273 | -0.54 |
| HB20 | 0.302 | 1.03 |
| CDF2 | 0.317 | 0.77 |
| SWEET | 0.322 | 0.91 |
| ADH1 | 0.35 | 0.06 |
| PP2C_1 | 0.366 | -0.23 |
| CML45 | 0.366 | -0.29 |
| RBA | 0.377 | -0.18 |
| CDF1 | 0.4 | 0.41 |
| TFL_SP | 0.425 | -0.54 |
| WRKY50 | 0.45 | -0.65 |
| PR1B | 0.502 | -0.28 |
| RD29B | 0.516 | -0.24 |
| OXI1 | 0.53 | -0.12 |
| FKF1 | 0.53 | -0.09 |
| PP2C_2 | 0.558 | 0.04 |
| CRINKLY4-like | 0.573 | -0.1 |
| CAT1 | 0.573 | 0.74 |
| XET | 0.623 | -0.49 |
| TOC1 | 0.647 | 0.24 |

| | | |
|---|---|---|
| RboHD | 0.663 | 0.03 |
| SNRK2 | 0.663 | 0.05 |
| TPL | 0.694 | -0.1 |
| ERF-1 | 0.694 | -0.01 |
| PIF4 | 0.758 | 0.06 |
| BEL5 | 0.79 | 0.33 |
| DSP4 | 0.823 | -0.06 |
| NFYA-1 | 0.84 | -0.01 |
| CYP704A2 | 0.907 | 0.01 |
| PPDK | 0.907 | -0.25 |
| P5CS1 | 0.924 | 0.33 |
| RBOHA | 0.958 | -0.09 |
| ICL | 0.992 | -0.05 |

### Section 4

Supplementary table S4.1: Spearman correlation coefficients and p-values for median tuber yield and quality variables (aggregated by variety).

| Feature 1 | Feature 2 | Spearman rho | p-value |
|---|---|---|---|
| median YHA | median UW | -0.63 | $1.4\times10^{-5}$ |
| median TP | median YHA | 0.46 | 0.003 |
| median OI | median UW | -0.45 | 0.004 |
| median YHA | median OI | 0.24 | 0.14 |
| median TP | median UW | -0.20 | 0.22 |
| median TP | median OI | 0.07 | 0.67 |

Supplementary table S4.2: Spearman correlation coefficients and p-values for yield per hectare (YHA) and features derived from drone-measured vegetation metrics.

| Feature | Spearman rho | p-value |
|---|---|---|
| wdvi_90 | 0.52 | 8.05E-39 |
| ndvi_90 | 0.5 | 2.82E-36 |
| vegetation_cover_90 | 0.48 | 1.29E-32 |
| vegetation_cover_120 | 0.45 | 1.13E-28 |
| wdvi_120 | 0.42 | 2.44E-25 |
| cired_90 | 0.42 | 6.07E-25 |
| ndvi_120 | 0.41 | 4.30E-24 |
| cired_120 | 0.39 | 3.23E-21 |
| vegetation_cover_40 | -0.19 | 1.05E-05 |
| cired_40 | -0.17 | 5.12E-05 |
| ndvi_60 | -0.15 | 2.93E-04 |
| ndvi_40 | -0.14 | 0.001 |
| cired_60 | -0.14 | 0.001 |
| vegetation_cover_60 | -0.12 | 0.004 |

| | | |
|---|---|---|
| wdvi_60 | -0.11 | 0.009 |
| wdvi_40 | -0.1 | 0.019 |

Supplementary table S4.3: Spearman correlation coefficients and p-values for tubers per plant (TP) and features derived from drone-measured vegetation metrics.

| **Feature** | **Spearman rho** | **p-value** |
|---|---|---|
| vegetation_cover_60 | 0.51 | 9.99E-38 |
| ndvi_60 | 0.49 | 5.43E-34 |
| wdvi_60 | 0.46 | 9.87E-30 |
| cired_60 | 0.45 | 1.33E-28 |
| vegetation_cover_40 | 0.35 | 1.54E-17 |
| ndvi_40 | 0.35 | 6.02E-17 |
| cired_40 | 0.32 | 1.06E-14 |
| wdvi_40 | 0.27 | 1.41E-10 |
| vegetation_cover_120 | -0.13 | 0.002 |
| ndvi_90 | 0.06 | 0.135 |
| ndvi_120 | -0.06 | 0.136 |
| cired_120 | -0.06 | 0.147 |
| wdvi_120 | -0.05 | 0.255 |
| cired_90 | 0.05 | 0.267 |
| vegetation_cover_90 | 0.02 | 0.645 |
| wdvi_90 | 0.01 | 0.730 |

Supplementary table S4.4: Spearman correlation coefficients and p-values for overall impression (OI) and features derived from drone-measured vegetation metrics.

| **Feature** | **Spearman rho** | **p-value** |
|---|---|---|
| wdvi_40 | -0.24 | 1.28E-08 |
| cired_40 | -0.23 | 5.29E-08 |
| cired_60 | -0.22 | 1.53E-07 |

| | | |
|---|---|---|
| wdvi_60 | -0.21 | 4.19E-07 |
| ndvi_40 | -0.21 | 5.37E-07 |
| ndvi_60 | -0.2 | 1.38E-06 |
| vegetation_cover_40 | -0.2 | 3.66E-06 |
| cired_90 | -0.17 | 4.64E-05 |
| ndvi_90 | -0.13 | 0.002 |
| vegetation_cover_60 | -0.13 | 0.003 |
| cired_120 | -0.11 | 0.008 |
| ndvi_120 | -0.11 | 0.010 |
| wdvi_120 | -0.1 | 0.016 |
| wdvi_90 | -0.08 | 0.063 |
| vegetation_cover_90 | -0.07 | 0.110 |
| vegetation_cover_120 | 0.02 | 0.683 |

Supplementary table S4.5: Spearman correlation coefficients and p-values for underwater weight (UW) and features derived from drone-measured vegetation metrics.

| **Feature** | **Spearman rho** | **p-value** |
|---|---|---|
| wdvi_60 | 0.23 | 5.01E-08 |
| cired_60 | 0.2 | 2.16E-06 |
| ndvi_60 | 0.2 | 2.41E-06 |
| vegetation_cover_60 | 0.13 | 0.003 |
| vegetation_cover_90 | -0.11 | 0.009 |
| cired_40 | 0.11 | 0.011 |
| vegetation_cover_40 | 0.09 | 0.030 |
| wdvi_90 | -0.09 | 0.042 |
| vegetation_cover_120 | -0.07 | 0.121 |
| cired_90 | 0.04 | 0.360 |
| ndvi_40 | 0.03 | 0.470 |

| | | |
|---|---|---|
| wdvi_120 | -0.03 | 0.536 |
| wdvi_40 | 0.02 | 0.698 |
| ndvi_90 | -0.01 | 0.763 |
| cired_120 | 0.01 | 0.798 |
| ndvi_120 | 0 | 0.993 |

Supplementary table S4.6: Spearman correlation coefficients and p-values for yield per hectare (YHA) and gene expression.

| **Gene** | **Spearman rho** | **p-value** |
|---|---|---|
| RBA | 0.59 | 6.43E-12 |
| CAT1 | 0.58 | 3.04E-11 |
| PPDK | 0.53 | 2.39E-09 |
| LTP3 | -0.49 | 4.78E-08 |
| CYP704A2 | 0.49 | 7.12E-08 |
| PIF4 | 0.42 | 3.71E-06 |
| BEL5 | 0.42 | 5.12E-06 |
| HB20 | 0.39 | 1.95E-05 |
| PP2C_2 | 0.36 | 1.21E-04 |
| ACX3 | 0.35 | 1.37E-04 |
| RBOHA | 0.34 | 2.17E-04 |
| HSC70 | 0.34 | 2.55E-04 |
| TOC1 | 0.33 | 3.69E-04 |
| DSP4 | -0.32 | 6.53E-04 |
| TPL | 0.31 | 8.32E-04 |
| XET | 0.30 | 0.001 |
| RD29B | 0.29 | 0.002 |
| GI2 | -0.27 | 0.005 |
| CDF1 | 0.26 | 0.007 |

| | | |
|---|---|---|
| P5CS1 | 0.24 | 0.010 |
| OXI1 | -0.24 | 0.011 |
| SWEET | -0.23 | 0.015 |
| SIS | 0.22 | 0.018 |
| CPK17 | -0.21 | 0.027 |
| WRKY50 | 0.20 | 0.034 |
| ACO2 | -0.20 | 0.036 |
| GI1 | 0.19 | 0.044 |
| RboHD | -0.19 | 0.050 |
| CO | 0.18 | 0.053 |
| SP5G | -0.18 | 0.064 |
| ICL | -0.15 | 0.123 |
| PP2C_1 | 0.14 | 0.134 |
| FKF1 | -0.14 | 0.154 |
| ADH1 | 0.14 | 0.153 |
| TFL_SP | -0.11 | 0.244 |
| Znf | -0.09 | 0.328 |
| PR1B | -0.07 | 0.440 |
| CDF2 | 0.06 | 0.519 |
| CML45 | 0.05 | 0.632 |
| CRINKLY4-like | -0.03 | 0.748 |
| NFYA-1 | 0.02 | 0.838 |
| SP6A | -0.02 | 0.837 |
| 13-LOX | -0.01 | 0.907 |
| SNRK2 | 0.00 | 0.984 |
| ERF-1 | 0.00 | 0.987 |

Supplementary table S4.7: Spearman correlation coefficients and p-values for tubers per plant and gene expression.

| **Gene** | **Spearman rho** | **p-value** |
|---|---|---|
| HSC70 | -0.35 | $1.6\times10^{-4}$ |
| CO | -0.27 | 0.005 |
| FKF1 | 0.25 | 0.008 |
| PR1B | -0.24 | 0.013 |
| ACO2 | 0.22 | 0.020 |
| CDF1 | -0.22 | 0.020 |
| GI2 | 0.22 | 0.022 |
| SIS | -0.22 | 0.022 |
| TFL_SP | -0.22 | 0.023 |
| DSP4 | 0.21 | 0.025 |
| XET | 0.21 | 0.027 |
| CDF2 | -0.18 | 0.065 |
| CML45 | -0.18 | 0.066 |
| TPL | -0.17 | 0.085 |
| WRKY50 | -0.16 | 0.097 |
| CYP704A2 | -0.16 | 0.098 |
| PPDK | -0.15 | 0.122 |
| SP5G | 0.13 | 0.179 |
| HB20 | -0.12 | 0.202 |
| ERF-1 | -0.12 | 0.211 |
| SNRK2 | 0.11 | 0.253 |
| OXI1 | -0.11 | 0.260 |
| ACX3 | -0.11 | 0.266 |
| TOC1 | 0.11 | 0.273 |

| | | |
|---|---|---|
| GI1 | 0.11 | 0.275 |
| SWEET | 0.10 | 0.310 |
| RBOHA | -0.10 | 0.317 |
| CRINKLY4-like | 0.10 | 0.318 |
| RBA | -0.09 | 0.332 |
| PP2C_2 | -0.09 | 0.346 |
| BEL5 | -0.09 | 0.358 |
| LTP3 | -0.08 | 0.420 |
| P5CS1 | 0.08 | 0.425 |
| Znf | -0.08 | 0.439 |
| PP2C_1 | 0.07 | 0.447 |
| 13-LOX | -0.07 | 0.475 |
| RboHD | -0.07 | 0.473 |
| ICL | 0.06 | 0.537 |
| ADH1 | 0.04 | 0.720 |
| RD29B | 0.02 | 0.869 |
| NFYA-1 | -0.02 | 0.865 |
| SP6A | -0.01 | 0.927 |
| CPK17 | -0.01 | 0.926 |
| PIF4 | 0.01 | 0.960 |
| CAT1 | 0.01 | 0.957 |

Supplementary table S4.8: Spearman correlation coefficients and p-values for overall impression and gene expression.

| **Gene** | **Spearman rho** | **p-value** |
|---|---|---|
| CAT1 | 0.37 | $7.0\times10^{-5}$ |
| PPDK | 0.31 | $8.2\times10^{-4}$ |
| Znf | 0.29 | 0.002 |

| | | |
|---|---|---|
| BEL5 | 0.29 | 0.002 |
| ERF-1 | 0.28 | 0.003 |
| RBA | 0.28 | 0.004 |
| PIF4 | 0.27 | 0.004 |
| LTP3 | -0.26 | 0.005 |
| SP5G | -0.24 | 0.011 |
| RBOHA | 0.22 | 0.018 |
| SIS | 0.22 | 0.023 |
| HB20 | 0.22 | 0.023 |
| SP6A | 0.21 | 0.024 |
| CO | 0.21 | 0.026 |
| PP2C_2 | 0.21 | 0.030 |
| CDF1 | 0.20 | 0.039 |
| TPL | 0.18 | 0.058 |
| RboHD | -0.18 | 0.066 |
| CML45 | 0.18 | 0.066 |
| CDF2 | 0.17 | 0.072 |
| ACX3 | 0.17 | 0.074 |
| CYP704A2 | 0.16 | 0.091 |
| ACO2 | -0.16 | 0.092 |
| HSC70 | 0.15 | 0.108 |
| FKF1 | -0.15 | 0.115 |
| ADH1 | -0.15 | 0.118 |
| OXI1 | -0.15 | 0.129 |
| GI2 | -0.13 | 0.173 |
| P5CS1 | 0.12 | 0.205 |
| WRKY50 | 0.11 | 0.237 |

| | | |
|---|---|---|
| CPK17 | -0.11 | 0.266 |
| PR1B | -0.10 | 0.317 |
| RD29B | 0.09 | 0.357 |
| PP2C_1 | -0.09 | 0.372 |
| SWEET | -0.07 | 0.453 |
| TOC1 | 0.07 | 0.458 |
| DSP4 | 0.06 | 0.531 |
| 13-LOX | 0.06 | 0.546 |
| SNRK2 | 0.05 | 0.624 |
| CRINKLY4-like | -0.04 | 0.694 |
| TFL_SP | 0.04 | 0.709 |
| NFYA-1 | -0.03 | 0.767 |
| GI1 | -0.01 | 0.881 |
| ICL | 0.01 | 0.952 |
| XET | 0.00 | 0.964 |

Supplementary table S4.9: Spearman correlation coefficients and p-values for underwater weight and gene expression.

| **Gene** | **Spearman rho** | **p-value** |
|---|---|---|
| PP2C_1 | 0.32 | $6.7\times10^{-4}$ |
| SP5G | 0.31 | 0.001 |
| TFL_SP | 0.30 | 0.002 |
| ICL | 0.27 | 0.004 |
| TPL | 0.24 | 0.014 |
| CRINKLY4-like | 0.21 | 0.031 |
| DSP4 | 0.18 | 0.060 |
| PPDK | -0.17 | 0.075 |
| CML45 | -0.14 | 0.135 |

| | | |
|---|---|---|
| HSC70 | -0.14 | 0.148 |
| PR1B | 0.14 | 0.149 |
| CPK17 | 0.14 | 0.157 |
| SNRK2 | 0.13 | 0.171 |
| BEL5 | 0.12 | 0.218 |
| WRKY50 | 0.11 | 0.252 |
| LTP3 | 0.11 | 0.264 |
| Znf | -0.11 | 0.274 |
| OXI1 | 0.10 | 0.323 |
| PP2C_2 | -0.10 | 0.327 |
| ERF-1 | -0.09 | 0.342 |
| ACO2 | 0.08 | 0.403 |
| HB20 | -0.08 | 0.437 |
| FKF1 | 0.07 | 0.448 |
| RBOHA | 0.07 | 0.464 |
| GI2 | 0.07 | 0.480 |
| PIF4 | -0.05 | 0.618 |
| CO | -0.05 | 0.625 |
| ADH1 | -0.05 | 0.631 |
| P5CS1 | 0.05 | 0.637 |
| CDF1 | 0.05 | 0.636 |
| GI1 | 0.04 | 0.659 |
| NFYA-1 | 0.04 | 0.689 |
| XET | 0.03 | 0.752 |
| TOC1 | 0.03 | 0.762 |
| RBA | -0.03 | 0.774 |
| RD29B | -0.02 | 0.842 |

| | | |
|---|---|---|
| RboHD | 0.02 | 0.867 |
| CYP704A2 | -0.01 | 0.889 |
| CDF2 | 0.01 | 0.912 |
| SIS | 0.01 | 0.927 |
| CAT1 | -0.01 | 0.925 |
| ACX3 | 0.01 | 0.924 |
| SP6A | 0.01 | 0.934 |
| SWEET | 0.01 | 0.953 |
| 13-LOX | 0.00 | 0.978 |

Supplementary table S4.10: Spearman correlation coefficients and p-values between features derived from drone-measured vegetation metrics.

| **Feature1** | **Feature2** | **Spearman rho** | **p-value** |
|---|---|---|---|
| cired_120 | ndvi_120 | 0.99 | $< 10^{-16}$ |
| cired_60 | ndvi_60 | 0.98 | $< 10^{-16}$ |
| cired_90 | ndvi_90 | 0.97 | $< 10^{-16}$ |
| ndvi_40 | wdvi_40 | 0.96 | $< 10^{-16}$ |
| ndvi_120 | wdvi_120 | 0.96 | $< 10^{-16}$ |
| cired_40 | vegetation_cover_40 | 0.95 | $< 10^{-16}$ |
| ndvi_60 | wdvi_60 | 0.95 | $< 10^{-16}$ |
| cired_60 | wdvi_60 | 0.95 | $< 10^{-16}$ |
| cired_120 | wdvi_120 | 0.95 | $< 10^{-16}$ |
| ndvi_40 | vegetation_cover_40 | 0.94 | $< 10^{-16}$ |
| cired_40 | wdvi_40 | 0.93 | $< 10^{-16}$ |
| ndvi_120 | vegetation_cover_120 | 0.92 | $< 10^{-16}$ |
| cired_40 | ndvi_40 | 0.91 | $< 10^{-16}$ |
| ndvi_90 | wdvi_90 | 0.91 | $< 10^{-16}$ |
| wdvi_40 | vegetation_cover_40 | 0.90 | $< 10^{-16}$ |
| cired_120 | vegetation_cover_120 | 0.89 | $< 10^{-16}$ |

| | | | |
|---|---|---|---|
| wdvi_120 | vegetation_cover_120 | 0.87 | $< 10^{-16}$ |
| cired_90 | wdvi_90 | 0.86 | $< 10^{-16}$ |
| ndvi_60 | vegetation_cover_60 | 0.82 | $< 10^{-16}$ |
| ndvi_40 | ndvi_60 | 0.82 | $< 10^{-16}$ |
| ndvi_40 | cired_60 | 0.80 | $< 10^{-16}$ |
| cired_40 | cired_60 | 0.78 | $< 10^{-16}$ |
| wdvi_40 | cired_60 | 0.78 | $< 10^{-16}$ |
| ndvi_90 | vegetation_cover_90 | 0.78 | $< 10^{-16}$ |
| ndvi_40 | wdvi_60 | 0.77 | $< 10^{-16}$ |
| cired_40 | ndvi_60 | 0.77 | $< 10^{-16}$ |
| wdvi_40 | ndvi_60 | 0.76 | $< 10^{-16}$ |
| wdvi_90 | vegetation_cover_90 | 0.76 | $< 10^{-16}$ |
| vegetation_cover_40 | vegetation_cover_60 | 0.75 | $< 10^{-16}$ |
| wdvi_60 | vegetation_cover_60 | 0.75 | $< 10^{-16}$ |
| cired_60 | vegetation_cover_60 | 0.75 | $< 10^{-16}$ |
| vegetation_cover_40 | ndvi_60 | 0.75 | $< 10^{-16}$ |
| wdvi_40 | wdvi_60 | 0.74 | $< 10^{-16}$ |
| cired_40 | wdvi_60 | 0.73 | $< 10^{-16}$ |
| vegetation_cover_40 | cired_60 | 0.72 | $< 10^{-16}$ |
| cired_90 | vegetation_cover_90 | 0.72 | $< 10^{-16}$ |
| ndvi_40 | vegetation_cover_60 | 0.71 | $< 10^{-16}$ |
| vegetation_cover_40 | wdvi_60 | 0.70 | $< 10^{-16}$ |
| cired_40 | vegetation_cover_60 | 0.70 | $< 10^{-16}$ |
| wdvi_40 | vegetation_cover_60 | 0.60 | $< 10^{-16}$ |
| ndvi_60 | vegetation_cover_120 | -0.57 | $< 10^{-16}$ |
| ndvi_90 | wdvi_120 | 0.56 | $< 10^{-16}$ |
| cired_90 | wdvi_120 | 0.56 | $< 10^{-16}$ |
| cired_90 | cired_120 | 0.56 | $< 10^{-16}$ |

| | | | |
|---|---|---|---|
| cired_60 | vegetation_cover_120 | -0.55 | $< 10^{-16}$ |
| vegetation_cover_90 | wdvi_120 | 0.55 | $< 10^{-16}$ |
| cired_90 | ndvi_120 | 0.54 | $< 10^{-16}$ |
| ndvi_40 | vegetation_cover_120 | -0.54 | $< 10^{-16}$ |
| vegetation_cover_90 | ndvi_120 | 0.54 | $< 10^{-16}$ |
| ndvi_90 | ndvi_120 | 0.54 | $< 10^{-16}$ |
| ndvi_90 | cired_120 | 0.53 | $< 10^{-16}$ |
| vegetation_cover_90 | cired_120 | 0.51 | $< 10^{-16}$ |
| cired_40 | vegetation_cover_120 | -0.50 | $< 10^{-16}$ |
| vegetation_cover_90 | vegetation_cover_120 | 0.48 | $< 10^{-16}$ |
| vegetation_cover_40 | vegetation_cover_120 | -0.48 | $< 10^{-16}$ |
| wdvi_40 | vegetation_cover_120 | -0.47 | $< 10^{-16}$ |
| wdvi_60 | vegetation_cover_120 | -0.46 | $< 10^{-16}$ |
| vegetation_cover_60 | vegetation_cover_120 | -0.46 | $< 10^{-16}$ |
| wdvi_90 | wdvi_120 | 0.45 | $< 10^{-16}$ |
| wdvi_90 | ndvi_120 | 0.37 | $< 10^{-16}$ |
| ndvi_90 | vegetation_cover_120 | 0.36 | $< 10^{-16}$ |
| wdvi_90 | cired_120 | 0.36 | $< 10^{-16}$ |
| ndvi_60 | ndvi_120 | -0.35 | $< 10^{-16}$ |
| ndvi_40 | ndvi_120 | -0.35 | $< 10^{-16}$ |
| ndvi_40 | cired_120 | -0.34 | $< 10^{-16}$ |
| ndvi_60 | cired_120 | -0.33 | 1.53E-15 |
| cired_90 | vegetation_cover_120 | 0.33 | 5.01E-15 |
| vegetation_cover_60 | cired_120 | -0.32 | 6.41E-15 |
| vegetation_cover_60 | ndvi_120 | -0.32 | 1.04E-14 |
| cired_60 | ndvi_120 | -0.30 | 9.42E-13 |
| cired_60 | cired_90 | 0.29 | 2.23E-12 |

| | | | |
|---|---|---|---|
| vegetation_cover_40 | ndvi_120 | -0.29 | 4.12E-12 |
| ndvi_60 | wdvi_120 | -0.28 | 2.38E-11 |
| vegetation_cover_40 | cired_120 | -0.28 | 2.70E-11 |
| ndvi_40 | wdvi_120 | -0.28 | 5.19E-11 |
| cired_60 | cired_120 | -0.27 | 1.31E-10 |
| wdvi_60 | cired_90 | 0.27 | 2.77E-10 |
| cired_40 | ndvi_120 | -0.26 | 6.78E-10 |
| vegetation_cover_60 | wdvi_120 | -0.26 | 1.18E-09 |
| vegetation_cover_40 | wdvi_120 | -0.25 | 4.46E-09 |
| wdvi_40 | ndvi_120 | -0.24 | 6.60E-09 |
| wdvi_90 | vegetation_cover_120 | 0.24 | 7.60E-09 |
| wdvi_40 | cired_90 | 0.24 | 8.28E-09 |
| cired_60 | wdvi_120 | -0.23 | 3.17E-08 |
| cired_40 | cired_120 | -0.23 | 3.35E-08 |
| wdvi_60 | ndvi_120 | -0.23 | 3.76E-08 |
| cired_60 | ndvi_90 | 0.23 | 4.22E-08 |
| wdvi_40 | cired_120 | -0.23 | 5.63E-08 |
| cired_40 | wdvi_120 | -0.22 | 2.97E-07 |
| wdvi_60 | cired_120 | -0.21 | 8.05E-07 |
| wdvi_60 | ndvi_90 | 0.21 | 1.19E-06 |
| ndvi_60 | cired_90 | 0.19 | 8.79E-06 |
| wdvi_40 | wdvi_120 | -0.18 | 2.35E-05 |
| wdvi_40 | ndvi_90 | 0.18 | 3.21E-05 |
| wdvi_60 | wdvi_120 | -0.18 | 3.24E-05 |
| cired_60 | wdvi_90 | 0.17 | 4.22E-05 |
| wdvi_40 | wdvi_90 | 0.16 | 1.45E-04 |
| cired_40 | cired_90 | 0.15 | 3.22E-04 |

| | | | |
|---|---|---|---|
| wdvi_60 | wdvi_90 | 0.15 | 3.69E-04 |
| ndvi_60 | ndvi_90 | 0.14 | 9.34E-04 |
| vegetation_cover_60 | wdvi_90 | -0.12 | 0.006 |
| vegetation_cover_40 | wdvi_90 | -0.11 | 0.009 |
| vegetation_cover_40 | vegetation_cover_90 | -0.11 | 0.013 |
| vegetation_cover_60 | vegetation_cover_90 | -0.10 | 0.015 |
| ndvi_60 | wdvi_90 | 0.10 | 0.021 |
| vegetation_cover_60 | ndvi_90 | -0.10 | 0.025 |
| ndvi_40 | cired_90 | 0.08 | 0.051 |
| vegetation_cover_40 | ndvi_90 | -0.08 | 0.054 |
| ndvi_60 | vegetation_cover_90 | -0.08 | 0.063 |
| vegetation_cover_60 | cired_90 | -0.08 | 0.072 |
| ndvi_40 | vegetation_cover_90 | -0.07 | 0.122 |
| cired_40 | ndvi_90 | 0.06 | 0.170 |
| wdvi_40 | vegetation_cover_90 | 0.05 | 0.245 |
| ndvi_40 | wdvi_90 | 0.04 | 0.401 |
| ndvi_40 | ndvi_90 | 0.04 | 0.418 |
| cired_60 | vegetation_cover_90 | -0.03 | 0.490 |
| cired_40 | vegetation_cover_90 | -0.02 | 0.597 |
| wdvi_60 | vegetation_cover_90 | -0.02 | 0.618 |
| cired_40 | wdvi_90 | 0.01 | 0.746 |
| vegetation_cover_40 | cired_90 | -0.01 | 0.863 |

Supplementary table S4.11: Spearman correlation coefficients and p-values between maturity scores and median (per variety) of features derived from drone-measured vegetation metrics.

| **Feature** | **Spearman rho** | **p-value** |
|---|---|---|
| ndvi_90_median | -0.84 | 1.29E-10 |

| | | |
|---|---|---|
| ndvi_120_median | -0.82 | 3.98E-10 |
| cired_120_median | -0.82 | 4.61E-10 |
| vegetation_cover_120_median | -0.82 | 5.29E-10 |
| wdvi_120_median | -0.81 | 1.14E-09 |
| cired_90_median | -0.77 | 2.03E-08 |
| wdvi_90_median | -0.69 | 2.70E-06 |
| vegetation_cover_90_median | -0.65 | 1.22E-05 |
| ndvi_60_median | -0.31 | 0.059 |
| wdvi_60_median | -0.26 | 0.114 |
| vegetation_cover_40_median | -0.21 | 0.215 |
| cired_60_median | -0.11 | 0.510 |
| cired_40_median | -0.10 | 0.541 |
| ndvi_40_median | -0.07 | 0.701 |
| vegetation_cover_60_median | -0.03 | 0.849 |
| wdvi_40_median | 0.00 | 0.984 |

Supplementary table S4.12: Spearman correlation coefficients and p-values between gene expression values. Note that only the pairs with absolute Spearman correlation coefficients ≥ 0.5 are shown.

| **Gene1** | **Gene2** | **Spearman rho** | **p-value** |
|---|---|---|---|
| FKF1 | GI2 | 0.93 | 4.83E-51 |
| CDF1 | CO | 0.9 | 1.06E-40 |
| CO | FKF1 | -0.9 | 1.63E-40 |
| CDF1 | CDF2 | 0.88 | 2.44E-37 |
| CDF2 | CO | 0.86 | 1.31E-34 |
| BEL5 | CAT1 | 0.86 | 2.17E-34 |
| CO | GI2 | -0.85 | 8.48E-33 |
| GI1 | SNRK2 | 0.84 | 8.07E-31 |

| | | | |
|---|---|---|---|
| CDF2 | FKF1 | -0.84 | 1.74E-30 |
| GI1 | TOC1 | 0.83 | 2.68E-30 |
| BEL5 | SIS | 0.8 | 2.27E-26 |
| SNRK2 | TOC1 | 0.79 | 2.01E-25 |
| BEL5 | TPL | 0.78 | 1.31E-24 |
| CDF1 | FKF1 | -0.78 | 3.17E-24 |
| CDF1 | GI2 | -0.77 | 5.94E-23 |
| CDF2 | GI2 | -0.77 | 7.02E-23 |
| FKF1 | SNRK2 | 0.77 | 7.26E-23 |
| BEL5 | PPDK | 0.76 | 3.42E-22 |
| PPDK | RBA | 0.75 | 6.93E-22 |
| HB20 | PP2C_2 | 0.75 | 8.89E-22 |
| GI2 | SNRK2 | 0.74 | 6.83E-21 |
| ACX3 | WRKY50 | 0.73 | 5.54E-20 |
| BEL5 | PIF4 | 0.73 | 7.46E-20 |
| CDF1 | SIS | 0.73 | 1.05E-19 |
| BEL5 | HB20 | 0.72 | 1.69E-19 |
| CYP704A2 | RBA | 0.72 | 2.60E-19 |
| HB20 | SIS | 0.72 | 3.81E-19 |
| PIF4 | PPDK | 0.72 | 5.98E-19 |
| CAT1 | TPL | 0.72 | 7.50E-19 |
| CML45 | PR1B | 0.71 | 9.50E-19 |
| BEL5 | CDF1 | 0.71 | 1.09E-18 |
| HSC70 | RBA | 0.71 | 3.42E-18 |
| RBA | TPL | 0.7 | 5.17E-18 |
| FKF1 | GI1 | 0.7 | 1.15E-17 |
| ACX3 | BEL5 | 0.69 | 4.18E-17 |

| | | | |
|---|---|---|---|
| RD29B | PP2C_2 | 0.69 | 4.56E-17 |
| 13-LOX | CML45 | 0.69 | 4.63E-17 |
| HB20 | PPDK | 0.68 | 9.15E-17 |
| BEL5 | RBA | 0.68 | 9.47E-17 |
| SIS | TPL | 0.68 | 1.30E-16 |
| PIF4 | TPL | 0.68 | 1.31E-16 |
| CAT1 | PPDK | 0.68 | 1.52E-16 |
| HB20 | RBA | 0.68 | 2.68E-16 |
| CYP704A2 | PP2C_2 | 0.68 | 3.09E-16 |
| PPDK | TPL | 0.67 | 3.43E-16 |
| ACX3 | PP2C_2 | 0.67 | 4.20E-16 |
| CRINKLY4-like | ICL | 0.67 | 9.77E-16 |
| CDF1 | PPDK | 0.67 | 1.26E-15 |
| SP6A | Znf | 0.66 | 2.91E-15 |
| CYP704A2 | HSC70 | 0.66 | 2.96E-15 |
| CAT1 | RBA | 0.65 | 5.24E-15 |
| ACX3 | PPDK | 0.65 | 5.74E-15 |
| ACX3 | RBOHA | 0.65 | 9.95E-15 |
| CYP704A2 | RBOHA | 0.65 | 1.03E-14 |
| HSC70 | PP2C_2 | 0.65 | 1.29E-14 |
| ACX3 | CAT1 | 0.65 | 1.54E-14 |
| RBA | PP2C_2 | 0.64 | 2.14E-14 |
| CAT1 | PIF4 | 0.64 | 3.89E-14 |
| GI1 | GI2 | 0.63 | 7.28E-14 |
| CAT1 | SIS | 0.63 | 8.38E-14 |
| PP2C_1 | TOC1 | 0.63 | 1.64E-13 |
| HSC70 | PPDK | 0.62 | 1.97E-13 |

| | | | |
|---|---|---|---|
| ACX3 | PIF4 | 0.62 | 2.12E-13 |
| HB20 | HSC70 | 0.62 | 2.16E-13 |
| BEL5 | PP2C_2 | 0.62 | 2.74E-13 |
| CO | SIS | 0.62 | 4.16E-13 |
| PP2C_1 | XET | 0.62 | 5.21E-13 |
| DSP4 | HSC70 | -0.61 | 1.01E-12 |
| CDF2 | SIS | 0.61 | 1.58E-12 |
| CO | SNRK2 | -0.6 | 1.89E-12 |
| RBOHA | WRKY50 | 0.6 | 2.09E-12 |
| 13-LOX | PR1B | 0.6 | 2.09E-12 |
| PPDK | SIS | 0.6 | 2.75E-12 |
| BEL5 | P5CS1 | 0.6 | 2.87E-12 |
| CYP704A2 | PPDK | 0.59 | 4.74E-12 |
| CYP704A2 | HB20 | 0.59 | 6.11E-12 |
| CDF2 | SNRK2 | -0.59 | 1.08E-11 |
| HB20 | P5CS1 | 0.59 | 1.15E-11 |
| RBA | RBOHA | 0.58 | 1.39E-11 |
| PIF4 | RBA | 0.58 | 1.43E-11 |
| CML45 | HSC70 | 0.58 | 2.17E-11 |
| DSP4 | ICL | 0.58 | 2.36E-11 |
| 13-LOX | ICL | -0.58 | 2.69E-11 |
| CO | GI1 | -0.57 | 3.44E-11 |
| RBOHA | PP2C_2 | 0.57 | 3.75E-11 |
| TOC1 | XET | 0.57 | 4.04E-11 |
| CAT1 | PP2C_2 | 0.57 | 4.39E-11 |
| HB20 | TPL | 0.57 | 5.16E-11 |
| CO | PPDK | 0.57 | 5.50E-11 |

| | | | |
|---|---|---|---|
| PIF4 | SIS | 0.57 | 7.00E-11 |
| CDF1 | TPL | 0.57 | 7.64E-11 |
| HB20 | PIF4 | 0.57 | 8.10E-11 |
| CAT1 | HB20 | 0.57 | 8.31E-11 |
| CDF1 | HB20 | 0.56 | 9.74E-11 |
| SIS | PP2C_2 | 0.56 | 1.01E-10 |
| ACX3 | RBA | 0.56 | 1.04E-10 |
| HSC70 | RBOHA | 0.56 | 1.12E-10 |
| FKF1 | TOC1 | 0.56 | 1.31E-10 |
| CDF2 | ERF-1 | 0.56 | 1.32E-10 |
| 13-LOX | SWEET | 0.56 | 1.66E-10 |
| ACO2 | DSP4 | 0.56 | 1.84E-10 |
| ACX3 | CYP704A2 | 0.56 | 1.90E-10 |
| CRINKLY4-like | RboHD | 0.55 | 2.35E-10 |
| CYP704A2 | TOC1 | 0.55 | 2.37E-10 |
| CRINKLY4-like | DSP4 | 0.55 | 2.39E-10 |
| 13-LOX | CRINKLY4-like | -0.55 | 2.87E-10 |
| PR1B | WRKY50 | 0.55 | 2.88E-10 |
| DSP4 | SP6A | 0.55 | 4.38E-10 |
| CDF2 | Znf | 0.54 | 5.70E-10 |
| SIS | Znf | 0.54 | 6.40E-10 |
| PPDK | RBOHA | 0.54 | 6.68E-10 |
| CDF1 | RBA | 0.54 | 7.05E-10 |
| PPDK | PP2C_2 | 0.54 | 7.92E-10 |
| ACX3 | TPL | 0.54 | 8.88E-10 |
| ACX3 | HB20 | 0.54 | 1.22E-09 |
| CDF2 | GI1 | -0.53 | 1.34E-09 |

| | | | |
|---|---|---|---|
| HB20 | RD29B | 0.53 | 1.49E-09 |
| ACX3 | CDF1 | 0.53 | 1.70E-09 |
| BEL5 | CO | 0.53 | 1.90E-09 |
| GI1 | XET | 0.53 | 1.93E-09 |
| PP2C_1 | PP2C_2 | 0.53 | 2.09E-09 |
| CDF2 | TFL_SP | 0.53 | 2.19E-09 |
| RBA | SIS | 0.53 | 2.21E-09 |
| P5CS1 | PP2C_2 | 0.52 | 3.13E-09 |
| 13-LOX | Znf | -0.52 | 3.28E-09 |
| ACX3 | SIS | 0.52 | 3.32E-09 |
| ICL | Znf | 0.52 | 3.62E-09 |
| CML45 | CRINKLY4-like | -0.52 | 3.84E-09 |
| CAT1 | P5CS1 | 0.52 | 3.92E-09 |
| ICL | TFL_SP | 0.52 | 4.14E-09 |
| GI1 | PP2C_1 | 0.52 | 5.19E-09 |
| CDF1 | PIF4 | 0.52 | 5.21E-09 |
| P5CS1 | TOC1 | 0.52 | 5.56E-09 |
| CO | TFL_SP | 0.52 | 5.89E-09 |
| CDF1 | TFL_SP | 0.51 | 7.47E-09 |
| CAT1 | CYP704A2 | 0.51 | 9.93E-09 |
| PP2C_1 | WRKY50 | 0.51 | 1.19E-08 |
| BEL5 | CDF2 | 0.51 | 1.21E-08 |
| BEL5 | CYP704A2 | 0.5 | 1.73E-08 |
| CAT1 | CDF1 | 0.5 | 1.76E-08 |
| CAT1 | RBOHA | 0.5 | 1.95E-08 |
| BEL5 | WRKY50 | 0.5 | 2.14E-08 |
| HSC70 | TPL | 0.5 | 2.21E-08 |

| | | | |
|---|---|---|---|
| CYP704A2 | TPL | 0.5 | 2.22E-08 |
| P5CS1 | SNRK2 | 0.5 | 2.75E-08 |

Supplementary table S4.13: Spearman correlation coefficients and p-values between gene expression values and features derived from drone-measured vegetation metrics. Note that only the pairs with absolute Spearman correlation coefficients ≥ 0.5 are shown.

| **Feature** | **Gene** | **Spearman rho** | **p-value** |
|---|---|---|---|
| wdvi_40 | BEL5 | -0.82 | 5.10E-28 |
| cired_40 | BEL5 | -0.79 | 3.39E-25 |
| ndvi_40 | BEL5 | -0.79 | 3.68E-25 |
| cired_60 | CDF1 | -0.78 | 2.47E-24 |
| ndvi_60 | CDF1 | -0.78 | 4.15E-24 |
| wdvi_40 | CDF1 | -0.77 | 3.68E-23 |
| cired_60 | CO | -0.76 | 2.24E-22 |
| ndvi_60 | CO | -0.76 | 6.20E-22 |
| vegetation_cover_40 | BEL5 | -0.74 | 1.25E-20 |
| cired_40 | CDF1 | -0.74 | 1.10E-20 |
| ndvi_40 | CDF1 | -0.74 | 3.14E-20 |
| vegetation_cover_120 | BEL5 | 0.72 | 9.37E-19 |
| wdvi_40 | TPL | -0.71 | 1.23E-18 |
| cired_40 | TPL | -0.71 | 1.48E-18 |
| ndvi_40 | TPL | -0.71 | 3.52E-18 |
| wdvi_40 | SIS | -0.71 | 4.83E-18 |
| wdvi_40 | PPDK | -0.71 | 4.63E-18 |
| vegetation_cover_60 | PPDK | -0.7 | 6.01E-18 |
| ndvi_40 | PPDK | -0.7 | 6.07E-18 |
| ndvi_60 | PPDK | -0.7 | 1.20E-17 |
| cired_40 | PPDK | -0.7 | 2.08E-17 |
| vegetation_cover_40 | PPDK | -0.7 | 2.36E-17 |
| cired_60 | PPDK | -0.69 | 5.03E-17 |

| | | | |
|---|---|---|---|
| ndvi_60 | GI2 | 0.68 | 1.77E-16 |
| ndvi_60 | BEL5 | -0.68 | 1.89E-16 |
| wdvi_60 | CDF1 | -0.68 | 1.99E-16 |
| cired_60 | BEL5 | -0.68 | 2.45E-16 |
| cired_60 | GI2 | 0.68 | 3.02E-16 |
| cired_40 | SIS | -0.68 | 3.11E-16 |
| vegetation_cover_40 | TPL | -0.68 | 3.92E-16 |
| vegetation_cover_120 | CDF1 | 0.68 | 4.09E-16 |
| wdvi_40 | CAT1 | -0.67 | 7.60E-16 |
| wdvi_40 | CO | -0.67 | 7.41E-16 |
| wdvi_60 | CO | -0.67 | 9.32E-16 |
| ndvi_40 | SIS | -0.66 | 1.88E-15 |
| vegetation_cover_60 | RBA | -0.66 | 4.44E-15 |
| ndvi_40 | CAT1 | -0.66 | 4.25E-15 |
| cired_40 | CAT1 | -0.66 | 5.56E-15 |
| cired_60 | FKF1 | 0.65 | 6.59E-15 |
| wdvi_60 | PPDK | -0.65 | 6.68E-15 |
| wdvi_60 | HSC70 | -0.65 | 6.68E-15 |
| cired_40 | CO | -0.65 | 7.92E-15 |
| wdvi_40 | HB20 | -0.65 | 8.55E-15 |
| wdvi_40 | PIF4 | -0.65 | 8.97E-15 |
| ndvi_60 | FKF1 | 0.65 | 8.84E-15 |
| wdvi_40 | RBA | -0.65 | 1.16E-14 |
| cired_40 | RBA | -0.65 | 1.36E-14 |
| cired_60 | SIS | -0.65 | 1.54E-14 |
| vegetation_cover_40 | HB20 | -0.65 | 1.53E-14 |
| ndvi_40 | PIF4 | -0.65 | 1.98E-14 |
| ndvi_40 | CO | -0.65 | 1.83E-14 |
| ndvi_40 | RBA | -0.64 | 2.22E-14 |

| | | | |
|---|---|---|---|
| vegetation_cover_40 | RBA | -0.65 | 2.10E-14 |
| cired_40 | PIF4 | -0.64 | 2.55E-14 |
| cired_40 | HB20 | -0.64 | 2.65E-14 |
| ndvi_40 | HB20 | -0.64 | 3.99E-14 |
| wdvi_60 | RBA | -0.64 | 4.44E-14 |
| ndvi_60 | SIS | -0.64 | 5.26E-14 |
| wdvi_60 | GI2 | 0.64 | 6.55E-14 |
| vegetation_cover_120 | CAT1 | 0.63 | 8.08E-14 |
| vegetation_cover_60 | TPL | -0.63 | 1.05E-13 |
| vegetation_cover_40 | CAT1 | -0.63 | 1.17E-13 |
| ndvi_60 | HSC70 | -0.63 | 1.84E-13 |
| vegetation_cover_40 | PIF4 | -0.63 | 2.16E-13 |
| vegetation_cover_40 | CDF1 | -0.62 | 2.50E-13 |
| ndvi_60 | RBA | -0.62 | 2.49E-13 |
| ndvi_60 | TPL | -0.62 | 2.98E-13 |
| cired_120 | BEL5 | 0.62 | 4.69E-13 |
| cired_60 | TPL | -0.62 | 4.39E-13 |
| cired_60 | CDF2 | -0.62 | 5.11E-13 |
| vegetation_cover_120 | SIS | 0.62 | 6.33E-13 |
| cired_60 | HSC70 | -0.61 | 9.54E-13 |
| cired_60 | RBA | -0.61 | 1.31E-12 |
| ndvi_60 | CDF2 | -0.6 | 2.03E-12 |
| vegetation_cover_60 | BEL5 | -0.6 | 2.16E-12 |
| wdvi_60 | BEL5 | -0.6 | 3.58E-12 |
| ndvi_60 | HB20 | -0.6 | 4.51E-12 |
| cired_90 | Znf | -0.59 | 6.69E-12 |
| cired_60 | HB20 | -0.59 | 1.39E-11 |
| ndvi_60 | PIF4 | -0.58 | 1.63E-11 |
| wdvi_90 | Znf | -0.58 | 1.63E-11 |

| | | | |
|---|---|---|---|
| wdvi_60 | FKF1 | 0.58 | 1.69E-11 |
| cired_60 | PIF4 | -0.58 | 2.11E-11 |
| vegetation_cover_120 | CO | 0.58 | 2.38E-11 |
| cired_120 | CAT1 | 0.58 | 2.73E-11 |
| ndvi_120 | BEL5 | 0.58 | 3.44E-11 |
| vegetation_cover_120 | GI2 | -0.57 | 4.24E-11 |
| vegetation_cover_40 | SIS | -0.57 | 6.00E-11 |
| vegetation_cover_60 | HSC70 | -0.57 | 7.57E-11 |
| vegetation_cover_60 | HB20 | -0.57 | 7.39E-11 |
| wdvi_40 | GI2 | 0.57 | 9.80E-11 |
| wdvi_40 | CDF2 | -0.57 | 9.30E-11 |
| wdvi_60 | HB20 | -0.57 | 1.01E-10 |
| vegetation_cover_120 | CDF2 | 0.56 | 1.25E-10 |
| ndvi_90 | Znf | -0.56 | 1.42E-10 |
| vegetation_cover_120 | RBA | 0.56 | 1.67E-10 |
| wdvi_40 | ACX3 | -0.56 | 2.42E-10 |
| vegetation_cover_120 | PIF4 | 0.56 | 2.52E-10 |
| vegetation_cover_40 | ACX3 | -0.55 | 2.79E-10 |
| vegetation_cover_120 | FKF1 | -0.55 | 3.93E-10 |
| cired_40 | ACX3 | -0.55 | 4.03E-10 |
| cired_40 | GI2 | 0.55 | 4.04E-10 |
| vegetation_cover_60 | CDF1 | -0.55 | 3.91E-10 |
| vegetation_cover_120 | PPDK | 0.55 | 4.50E-10 |
| ndvi_40 | ACX3 | -0.55 | 4.61E-10 |
| ndvi_40 | GI2 | 0.55 | 4.73E-10 |
| ndvi_120 | CAT1 | 0.54 | 6.64E-10 |
| vegetation_cover_40 | CO | -0.54 | 8.84E-10 |
| vegetation_cover_120 | TPL | 0.54 | 1.18E-09 |
| wdvi_120 | BEL5 | 0.54 | 1.31E-09 |

| | | | |
|---|---|---|---|
| vegetation_cover_40 | HSC70 | -0.54 | 1.42E-09 |
| wdvi_40 | TFL_SP | -0.53 | 1.80E-09 |
| cired_40 | CDF2 | -0.53 | 2.02E-09 |
| wdvi_60 | SIS | -0.53 | 2.05E-09 |
| wdvi_60 | TPL | -0.53 | 2.41E-09 |
| vegetation_cover_120 | HB20 | 0.53 | 2.52E-09 |
| wdvi_40 | FKF1 | 0.53 | 2.72E-09 |
| ndvi_40 | CDF2 | -0.52 | 4.30E-09 |
| cired_60 | TFL_SP | -0.52 | 4.39E-09 |
| wdvi_120 | CAT1 | 0.52 | 4.72E-09 |
| cired_40 | HSC70 | -0.52 | 4.43E-09 |
| ndvi_40 | HSC70 | -0.52 | 6.54E-09 |
| ndvi_60 | CAT1 | -0.52 | 6.54E-09 |
| vegetation_cover_40 | CYP704A2 | -0.51 | 7.91E-09 |
| cired_60 | CAT1 | -0.51 | 8.55E-09 |
| wdvi_60 | PIF4 | -0.51 | 9.07E-09 |
| cired_120 | CDF1 | 0.51 | 9.54E-09 |
| cired_40 | FKF1 | 0.51 | 9.92E-09 |
| cired_90 | TOC1 | 0.51 | 1.10E-08 |
| cired_40 | TFL_SP | -0.51 | 1.39E-08 |
| vegetation_cover_60 | PIF4 | -0.51 | 1.46E-08 |
| wdvi_40 | HSC70 | -0.51 | 1.50E-08 |
| ndvi_60 | TFL_SP | -0.51 | 1.54E-08 |
| wdvi_90 | CYP704A2 | 0.5 | 1.82E-08 |
| ndvi_40 | FKF1 | 0.5 | 1.99E-08 |
| vegetation_cover_60 | CAT1 | -0.5 | 2.00E-08 |
| cired_120 | RBA | 0.5 | 2.20E-08 |
| wdvi_60 | CDF2 | -0.5 | 2.27E-08 |
| ndvi_120 | CDF1 | 0.5 | 2.51E-08 |

| | | | |
|---|---|---|---|
| ndvi_90 | CYP704A2 | 0.5 | 2.59E-08 |
| vegetation_cover_60 | CYP704A2 | -0.5 | 2.90E-08 |

#### Section 5

Supplementary table S5.1: Set of the 32 selected non-highly correlated features with categories. For description of environmental features refer to Supplementary table S1.5.

| **Feature** | **Category** |
|---|---|
| num_days_min_temp_night_above_16_90-120 | Environmental (air temperature) |
| tempavg_median_90-120 | Environmental (air temperature) |
| tempmax_median_90-120 | Environmental (air temperature) |
| tempavg_median_0-40 | Environmental (air temperature) |
| tempmax_median_0-40 | Environmental (air temperature) |
| tempavg_median_40-60 | Environmental (air temperature) |
| daylength_median_90-120 | Environmental (daylength) |
| daylength_median_0-40 | Environmental (daylength) |
| humidity_min_90-120 | Environmental (humidity) |
| humidity_max_0-40 | Environmental (humidity) |
| humidity_median_0-40 | Environmental (humidity) |
| num_days_humidity_below_60_0-40 | Environmental (humidity) |
| humidity_max_40-60 | Environmental (humidity) |
| humidity_median_40-60 | Environmental (humidity) |
| num_days_humidity_below_60_40-60 | Environmental (humidity) |
| precip_median_0-40 | Environmental (precipitation) |
| precip_sum_0-40 | Environmental (precipitation) |
| avg_soil_moisture_max_90-120 | Environmental (soil moisture) |
| avg_soil_moisture_night_max_60-90 | Environmental (soil moisture) |
| num_days_avg_soil_moisture_below_20_90-120 | Environmental (soil moisture) |
| avg_soil_temp_day_median_90-120 | Environmental (soil temperature) |
| num_days_avg_soil_temp_night_above_20_60-90 | Environmental (soil temperature) |
| num_days_min_soil_temp_night_above_18_60-90 | Environmental (soil temperature) |
| num_days_min_soil_temp_night_above_20_90-120 | Environmental (soil temperature) |
| solarradiation_median_60-90 | Environmental (solar radiation) |
| maturity_score | Maturity Score |
| ndvi_40 | Vegetation metrics |
| ndvi_60 | Vegetation metrics |
| vegetation_cover_60 | Vegetation metrics |
| ndvi_90 | Vegetation metrics |

| | |
|---|---|
| vegetation_cover_90 | Vegetation metrics |
| ndvi_120 | Vegetation metrics |

Supplementary table S5.2: Performance, expressed as coefficient of determination (R2), of kernel ridge regression models on the test set for predicting the four targets: yield per hectare (YHA), number of tubers per plant (TP), overall impression (OI), and underwater weight (UW), evaluated using data restricted to features available up to 40, 60, 90, and 120 days post planting (dpp) during training.

| **Threshold (dpp)** | **Target** | **R2** |
|---|---|---|
| 40 | YHA | 0.71 |
| | TP | 0.16 |
| | OI | 0.12 |
| | UW | 0.06 |
| 60 | YHA | 0.77 |
| | TP | 0.14 |
| | OI | 0.16 |
| | UW | 0.08 |
| 90 | YHA | 0.80 |
| | TP | 0.16 |
| | OI | 0.16 |
| | UW | 0.11 |
| 120 | YHA | 0.78 |
| | TP | 0.17 |
| | OI | 0.16 |
| | UW | 0.11 |

Supplementary table S5.3: Predictive performance, expressed as the coefficient of determination ($R^2$), for yield per hecate using kernel ridge regression models trained on different combinations of input feature groups.

| **Feature subset(s)** | **Number of features** | **Test R2** |
|---|---|---|
| maturity score | 1 | -0.08 |

| | | |
|---|---|---|
| vegetation metrics | 6 | 0.69 |
| vegetation metrics + environmental features | 31 | 0.79 |
| vegetation metrics + maturity score | 7 | 0.69 |
| vegetation metrics + gene expression | 47 | 0.6 |
| vegetation metrics + maturity score + gene expression | 48 | 0.6 |
| vegetation metrics + environmental features + maturity score | 32 | 0.78 |
| vegetation metrics + environmental features + gene expression | 72 | 0.7 |
| vegetation metrics + environmental features + maturity score + gene expression | 73 | 0.7 |
| environmental features | 25 | 0.72 |
| environmental features + maturity score | 26 | 0.72 |
| environmental features + gene expression | 66 | 0.5 |
| environmental features + maturity score + gene expression | 67 | 0.5 |
| gene expression | 41 | 0.37 |
| gene expression + maturity score | 42 | 0.37 |

### Section 6

Supplementary table S6.1: Spearman correlation coefficients and p-values between the five selected features for linear regression equation for yield per hectare.

| **Feature 1** | **Feature 2** | **Spearman rho** | **p-value** |
|---|---|---|---|
| ndvi_40 | vegetation_cover_60 | 0.71 | 5.7E-86 |
| ndvi_40 | num_days_humidity_below_60_40-60 | -0.54 | 2.6E-42 |
| vegetation_cover_60 | precip_sum_0-40 | 0.43 | 2.8E-26 |
| vegetation_cover_60 | num_days_humidity_below_60_40-60 | -0.4 | 1.2E-22 |
| humidity_max_0-40 | precip_sum_0-40 | 0.4 | 2.1E-22 |
| ndvi_40 | precip_sum_0-40 | 0.29 | 8.9E-12 |
| humidity_max_0-40 | num_days_humidity_below_60_40-60 | -0.16 | 2.2E-04 |
| precip_sum_0-40 | num_days_humidity_below_60_40-60 | -0.09 | 0.026 |
| ndvi_40 | humidity_max_0-40 | -0.08 | 0.055 |
| vegetation_cover_60 | humidity_max_0-40 | 0.08 | 0.078 |

### Methods

Supplementary table SM1. Description of field trial parameters across locations.

| **Field trial** | **Planting date** | **Trial design** | **Irrigation type** | **Irrigation freq.** | **Plot size** | **Number of plants per plot** | **Number of plots per variety** | **Soil type** | **Drone provider** | **Management type** |
|---|---|---|---|---|---|---|---|---|---|---|
| Valencia 21 (irr.) | 3.02.21 | Randomized block | Flood irrigation | irregular | 1,63 | 8 | 2 | Clay loam | Aurea Imaging | Conventional |
| Valencia 21 (non-irr.) | 14.04.21 | Randomized block | - | - | 1,63 | 8 | 2 | Clay loam | Aurea Imaging | Conventional |
| Valencia 22 (irr.) | 28.01.22 | Randomized block | Flood irrigation | irregular | 1,63 | 8 | 2 | Clay loam | Aurea Imaging | Conventional |
| Vojvodina 23 (irr.) | 28.04.23 | Randomized block | Sprinkler irrigation | irregular | 2,7 | 12 | 2 | clay | Aurea Imaging | conventional |
| Zeeland 21 (irr.) | 3.05.21 | Randomized block | Drip irrigation | irregular | 1,65 m x 1,5 m | 8 plants (+4 borderplants) | 2 | Light clay | Aurea Imaging | Conventional |
| Zeeland 21 (non-irr.) | 3.05.21 | Randomized block | - | - | 1,65 m x 1,5 m | 8 plants (+4 borderplants) | 2 | Light clay | Aurea Imaging | Conventional |
| Zeeland 22 (irr.) | 2.05.22 | Randomized block | Drip irrigation | irregular | 1,65 m x 1,5 m | 8 plants (+4 borderplants) | 2 | Light clay | Aurea Imaging | Conventional |
| Zeeland 22 (non-irr.) | 2.05.22 | Randomized block | - | - | 1,65 m x 1,5 m | 8 plants (+4 borderplants) | 2 | Light clay | Aurea Imaging | Conventional |
| Zeeland 23 (irr.) | 22.05.23 | Randomized block | Drip irrigation | Irregular | 1,65 m x 1,5 m | 8 plants (+4 borderplants) | 2 | Light clay | Aurea Imaging | Conventional |
| Zeeland 23 (non-irr.) | 22.05.23 | Randomized block | - | - | | 8 plants (+4 borderplants) | 2 | Light clay | Aurea Imaging | Conventional |
| Fuchsenbigl 22 (irr.) | 21.04.22 | Randomized block | Sprinkler irrigation | 4 times | 12m2 | 50 | 4 | Black | Blickwinkel Agrarconsulting, M. Treiblmeier | Conventional |
| Fuchsenbigl 22 (non-irr.) | 21.04.22 | Randomized block | - | - | 12m2 | 50 | 4 | Black | Blickwinkel Agrarconsulting, M. Treiblmeier | Conventional |

| | | | | | | | | | | |
|---|---|---|---|---|---|---|---|---|---|---|
| Fuchsenbigl 23 (irr.) | 26.04.23 | Randomized block | Sprinkler irrigation | 3 times | 12m2 | 50 | 4 | Black | Blickwinkel Agrarconsulting, M. Treiblmeier | Conventional |
| Fuchsenbigl 23 (non-irr.) | 26.04.23 | Randomized block | - | - | 12m2 | 50 | 4 | Black | Blickwinkel Agrarconsulting, M. Treiblmeier | Conventional |
| Großnondorf 22 (non-irr.) | 14.04.22 | Randomized block | - | - | 12m2 | 50 | 4 | Black | Blickwinkel Agrarconsulting, M. Treiblmeier | Conventional |
| Großnondorf 23 (non-irr.) | 26.04.23 | Randomized block | - | - | 12m2 | 50 | 4 | Black | Blickwinkel Agrarconsulting, M. Treiblmeier | Conventional |

Supplementary Table SM2. Geographic coordinates of the Visual Crossing Weather Data Services stations used to retrieve weather data for each field trial site.

| **Location** | **Latitude** | **Longitude** |
|---|---|---|
| Fuchsenbigl | 48.197125 | 16.752882 |
| Großnondorf | 48.649207 | 16.023260 |
| Valencia | 39.214 | -0.313 |
| Vojvodina | 45.406588 | 19.511469 |
| Zeeland | 51.495427 | 3.758976 |

Supplementary Table SM3. Tested hyperparameters for regression models used in this study. The table lists the range of values explored for each model during grid search in the inner loops of the nested cross-validation.

| **Model** | **Tested hyperparameters** |
|---|---|
| LinearRegression | fit_intercept: [True] |
| Ridge | alpha: [0.1, 1.0, 10.0, 50.0, 100.0, 500.0] |
| Lasso | alpha: [0.01, 0.1, 1.0, 10.0] |
| ElasticNet | alpha: [0.1, 1.0, 10.0, 50.0, 100.0]; l1_ratio: [0.01, 0.1, 0.5, 0.9] |
| RandomForest | n_estimators: [50, 100, 150, 200]; max_depth: [None, 3, 4, 5, 10]; min_samples_split: [5, 10, 20, 30]; min_samples_leaf: [1, 2, 10]; max_features: [None, 'sqrt', 'log2'] |
| XGBoost | n_estimators: [100, 150, 200]; max_depth: [3, 6]; learning_rate: [0.01, 0.05, 0.1]; subsample: [0.2, 0.4, 0.6, 0.8]; colsample_bytree: [0.2, 0.4, 0.6, 0.8]; lambda: [0, 0.01, 1.0, 5.0] |

| | |
|---|---|
| KernelRidge | alpha: [0.01, 0.1, 1.0, 10.0]; kernel: ['rbf', 'poly']; gamma: [0.001, 0.01, 0.1, 1.0]; degree: [2, 3, 4] |
| SVR | C: [0.1, 1.0, 10.0, 100]; epsilon: [0.01, 0.1, 0.5]; kernel: ['linear', 'rbf', 'poly']; gamma: [0.001, 0.01, 0.1] |

Supplementary Table SM4. Selected hyperparameters for the kernel ridge models for each target variable

| **Target** | **Hyperparameters** |
|---|---|
| YHA | alpha: 0.1;<br>gamma: 0.01;<br>kernel: poly;<br>degree: 3 |
| TP | alpha: 0.1;<br>gamma: 0.001;<br>kernel: poly;<br>degree: 4 |
| OI | alpha: 0.01;<br>gamma: 0.001;<br>kernel: poly;<br>degree: 2 |
| UW | alpha: 0.01;<br>gamma: 0.001;<br>kernel: poly;<br>degree: 3 |
